# Minimally invasive classification of pediatric solid tumors using reduced representation bisulfite sequencing of cell-free DNA: a proof-of-principle study

**DOI:** 10.1101/795047

**Authors:** Ruben Van Paemel, Andries De Koker, Charlotte Vandeputte, Lieke van Zogchel, Tim Lammens, Geneviève Laureys, Jo Vandesompele, Gudrun Schleiermacher, Mathieu Chicard, Nadine Van Roy, Ales Vicha, G.A.M. Tytgat, Nico Callewaert, Katleen De Preter, Bram De Wilde

## Abstract

In the clinical management of pediatric solid tumors, histological examination of tumor tissue obtained by a biopsy remains the gold standard to establish a conclusive pathological diagnosis. The DNA methylation pattern of a tumor is known to correlate with the histopathological diagnosis across cancer types and is showing promise in the diagnostic workup of tumor samples. This methylation pattern can be detected in the cell-free DNA. Here, we provide proof-of-concept of histopathologic classification of pediatric tumors using cell-free reduced representation bisulfite sequencing (cf-RRBS) from retrospectively collected plasma and cerebrospinal fluid samples. We determined the correct tumor type in 49 out of 60 (81.6%) samples starting from minute amounts (less than 10 ng) of cell-free DNA. We demonstrate that the majority of misclassifications were associated with sample quality and not with the extent of disease. Our approach has the potential to help tackle some of the remaining diagnostic challenges in pediatric oncology in a cost-effective and minimally invasive manner.

**Translational relevance:** Obtaining a correct diagnosis in pediatric oncology can be challenging in some tumor types, especially in renal tumors or central nervous system tumors. Furthermore, the diagnostic odyssey can result in anxiety and discomfort for these children. By applying a novel technique, reduced representation bisulfite sequencing on cell-free DNA (cf-RRBS), we show the feasibility of obtaining the histopathological diagnosis with a minimally invasive test on either plasma or cerebrospinal fluid. Furthermore, we were able to derive the copy number profile or tumor subtype from the same assay. Given that primary tumor material might be difficult to obtain, in particular in critically ill children or depending on the tumor location, and might be limited in terms of quantity or quality, our assay could become complementary to the classical tissue biopsy in difficult cases.

## Introduction

The diagnostic work-up of pediatric cancer patients with solid tumors requires a number of investigations to obtain a full diagnosis and a complete staging. Most imaging and tumor marker investigations are of low specificity and do not result in a definitive diagnosis. Thus, in most cases, a histopathological examination of a surgically derived tumor biopsy is required. Many of the pediatric tumors share a “small blue” histology and show few lineage-specific morphological features. Hence, extensive immunohistochemistry staining is often required to establish a diagnosis. Nevertheless, some types of embryonic tumors or sarcomas remain hard to classify correctly. In the context of international frontline clinical trial protocols tumor specimens are often sent for central pathology review. In a small percentage of patients, discrepancy in either stage or diagnosis can result in suboptimal treatment^1,2^. In pediatric renal tumors, the rate of misdiagnosis was estimated to range from 3.5% to 17% based on multicenter clinical trials^1^. Limited tissue availability due to fine needle biopsy sampling can further complicate the diagnostic work-up. Furthermore, for some types of pediatric tumors, sampling of the tumor is not routinely recommended in some treatment protocols (e.g. SIOP protocol for treatment of renal tumors^3,4^) or is deemed too risky due to the location (e.g. diffuse intrinsic pontine glioma) or clinical condition of the patient (e.g. contra-indications for anesthesia due to extensive disease). Thus, therapy needs to be started empirically based on imaging and clinical characteristics alone, increasing the odds for misdiagnosis.

The epigenetic signal of primary tissue samples is currently being explored to complement classical histopathological analysis. This epigenetic signal seems to be highly tissue-specific^5–7^ as opposed to most mutational or copy number alterations. This tissue specificity is the dominant remaining signal after malignant transformation, resulting in a unique fingerprint of DNA for different tumor entities^8,9^. For example, the DNA methylation signal contributed to obtaining the correct diagnosis in 25 out of 30 cases in a cohort of translocation-negative Ewing sarcoma samples^10^. Among brain tumors, primitive neuro-ectodermal tumors of the central nervous system (CNS-PNETs) are notoriously difficult to diagnose based on histology alone. DNA methylation revealed that this entity is actually composed of multiple known and previously unknown tumor types^11^. Furthermore, in a large cohort of 1155 prospectively collected brain tumor samples, 12% of histopathological diagnoses could be adjusted based on tumor DNA methylation characteristics^12^. In medulloblastoma, prospective clinical trials are currently ongoing to test whether the adaptation of treatment based on new multi-omics (including DNA methylation) improves event-free survival (NCT02066220).

Recently, analysis of cell-free tumor-derived DNA (ctDNA) isolated from liquid biopsies including blood has emerged as a complementary assay for tumor tissue genomic profiling^13^. Several groups have investigated the use of cell-free DNA methylation for noninvasive diagnosis of both benign^14^ and malignant^15^ conditions. For example, Kang *et al*. developed CancerLocator^8^, an algorithm based on the epigenetic signature determined on whole genome bisulfite sequencing (WGBS) to classify cell-free DNA of adult cancer patients to their corresponding tumor tissue. Through data deconvolution, the CancerLocator method was able to correctly classify samples, even if they had a low percentage of circulating tumor DNA, outperforming standard machine-learning approaches.

Although tumor classification from the epigenetic cfDNA profile is possible, the high cost of WGBS limits its use in routine clinical diagnostics and follow-up. Reduced representation bisulfite sequencing (RRBS) is a cost-effective alternative whereby only a small informative fraction of the genome is surveyed ^16^. Enrichment of CpG-rich sites by digesting DNA with the MspI restriction enzyme (cuts 5’-C/CGG-3’), followed by DNA fragment size-selection is a commonly applied strategy of RRBS. This method was redesigned by De Koker *et al*.^17^ to allow single tube RRBS library preparation on low input levels (10 ng) of low-quality and/or fragmented DNA.

Our study investigates whether the methylome profile generated with RRBS on cell-free DNA (cf-RRBS) is capable of classifying pediatric tumors according to their histopathological diagnosis. To this end, we profiled plasma or cerebrospinal fluid of 59 patients with pediatric cancer. This profile was used to classify the sample according to a reference dataset built from publicly available methylome data of pediatric tumors of known histology. We further explored the tumor copy number profile extracted from cf-RRBS comparing it against the copy number profile obtained from shallow whole genome sequencing data.

## Materials and methods

### Patients and samples

Pediatric cancer patients (median age 46 months [18.5 – 114.5]) were included retrospectively if plasma or CSF was available at diagnosis (n = 58) or at relapse (n = 1). For one case (patient_36, clear cell sarcoma of the kidney), two plasma samples were available at different timepoints, resulting in a total of 60 samples. Samples were collected at Ghent University Hospital (Ghent, Belgium, n = 20), Prinses Máxima Centrum (Utrecht, The Netherlands, n = 15), Institut Curie (Paris, France, n = 6) and Prague Motol University Hospital (Prague, Czech Republic, n = 19). Blood was collected from patients with neuroblastoma (n = 10), nephroblastoma (n = 16), malignant rhabdoid tumor of the kidney (n = 1), clear cell sarcoma of the kidney (n = 2), alveolar rhabdomyosarcoma (n = 9), embryonal rhabdomyosarcoma (n = 8), osteosarcoma (n = 4) and Ewing sarcoma (n = 6). Cerebrospinal fluid was collected from patients with medulloblastoma (n = 3) and atypical teratoid-rhabdoid tumor (n = 1). The local ethical committee approved the study and written consent was obtained from all patients enrolled in this study or their representatives. Per-sample information is available in supplementary table 2. Descriptive statistics were performed with R v3.5.1 and Python v3.6.3 and are expressed by median [Q25 – Q75]. Unless otherwise stated, the non-parametric Wilcoxon rank sum test was used to test for significance with alpha specified at 0.05.

### Sample collection and processing

#### Ghent University Hospital

Whole blood was collected in citrate tubes (Greiner Bio-One, n = 16). Plasma was obtained after 1 × 8 min centrifugation at 1885 g. The time between blood collection and plasma preparation was not documented. All plasma samples were stored at −80 °C until processing for cfDNA extraction. CSF was collected into tubes without additives. The CSF was stored at −80 °C until further processing for cfDNA extraction.

#### Prinses Máxima Center

Whole blood from all patients was collected in EDTA Vacutainer tubes (BD Biosciences). Plasma was obtained after 1 × 10 min centrifugation at 1375 g with centrifuge acceleration and without deceleration. The plasma was stored at −20 °C until processing for cfDNA extraction. Plasma was prepared within 24 hours after blood collection.

#### Prague Motol University Hospital

Whole blood from all patients was collected in EDTA Vacutainer tubes (BD Biosciences). Plasma was obtained after 1 × 10 min centrifugation at 1600 g at room temperature, centrifuge acceleration, and deceleration was set to two, followed by a subsequent centrifugation step of 1 × 10 minutes at 16,000 g at room temperature. The plasma was stored at −80 °C until processing for cfDNA extraction. Before the first centrifugation step, 2.5% (v/v) of a 10% formaldehyde solution was added. Plasma was prepared within 4 hours after blood draw.

#### Institut Curie

Whole blood was collected in EDTA tubes. Plasma was obtained after 1 × 20 min centrifugation at 2000 g at room temperature, centrifuge acceleration, and deceleration was set to two, followed by a subsequent centrifugation step of 1 × 2 minutes at 16,000 g at room temperature. The plasma was stored at −80 °C until processing for cfDNA extraction.

### Cell-free DNA extraction

#### Ghent University Hospital and Prinses Maxima Center

cfDNA was extracted using the Maxwell RSC LV ccfDNA kit (Promega). Isolation of cfDNA was done starting from 200 μL to 2 mL of plasma or CSF, and if less volume was available, volumes were adjusted to 2 mL by adding 1x PBS (Gibco). DNA extraction was performed according to the manufacturer’s instructions. DNA was eluted in 75 μL of elution buffer (Promega).

#### Prague Motol University Hospital

cfDNA was extracted using the QIAamp Circulating Nucleic Acid kit (Qiagen). Isolation of cfDNA was done from 1 mL of plasma. DNA extraction was performed according to the manufacturer’s instructions. DNA was eluted in 25 μL of AVE buffer (Qiagen).

#### Institut Curie

cfDNA was extracted using QIAamp Circulating Nucleic Acid Kit (Qiagen) with the Qiavac24s system, according to the manufacturers’ recommendations. Isolation of cfDNA was done starting from 200 μL to 1.5 mL of plasma. Volumes were adjusted to 2 mL by adding 1x PBS (Gibco). DNA was eluted in 36 μL of AVE buffer (Qiagen).

### Cell-free DNA quality control

DNA concentration was measured using the Qubit high-sensitivity kit (Thermo Fisher Scientific). Size distribution of the cfDNA was measured using the FEMTO Pulse Automated Pulsed-Field CE Instrument (Agilent) according to the manufacturer’s instructions (NGS Kit, FP-1101-0275). The ratio between cell-free DNA and high-molecular weight DNA (HMW) was calculated with the Prosize 3.0 software (Agilent). To this end, two blinded independent observers visually selected the cfDNA regions and high molecular weight (HMW) regions, excluding the upper marker. Then, the percentage of cfDNA was divided by the percentage of HMW DNA to obtain the cfDNA/HMW ratio. Detailed figures are shown in supplementary figures. Because this interpretation is inherently subjective, the coefficient of determination (R^2^) and Cohen’s Kappa coefficient between both observers was calculated. R^2^ between the two observers was 0.91. Samples with a cfDNA/HMW ratio less than 1 were labeled as “high HMW contamination”, samples with a cfDNA/HMW ratio between 1 and 5 “medium HMW contamination” and samples with a cfDNA/HMW ratio >5 as “low HMW contamination”. Cohen’s Kappa coefficient with equal weights between the two observers was 0.798 (p < 0.001). We then evaluated these cut-offs by collecting blood from 3 healthy volunteers in EDTA tubes. The tubes were either processed immediately or left standing at room temperature for 24 or 72 hours. The blood tubes that were processed immediately all showed a cfDNA/HMW ratio above 5 (11.48, 9.01 and 36.03, respectively). The cfDNA/HMW ratio of the EDTA tubes that were left standing for 24h were all between 1 and 5 (5.02, 1.37 and 4.05, respectively) and the cfDNA/HMW ratio of the tubes that were left standing for 72h were all lower than 1 (0.33, 0.20 and 0.42, respectively).

### RRBS library construction

Median input DNA for the library construction was 10.00 ng [7.65 – 10.19]. Library construction was performed according to the methods described by De Koker *et al*. ^17^ with the following modifications: [1] If DNA concentration was lower than 0.2 ng/μL, samples were concentrated with a vacuum centrifuge (SpeedVac, Thermo Fischer Scientific) at 35°C and H_2_O was added to a volume of 10.6 or 11.1 μL, depending on the amount of lambda spike-in (1 μL or 0.5 μL of a 0.01 ng/μL solution). [2] Libraries prepared using the cf-RRBS protocol were cleaned by magnetic bead selection (AMPure XT beads – NEB) and eluted in 0.1X TE buffer. The libraries were visualized with the Fragment Analyzer (Advanced Analytical Technologies) and quantified using the Kapa library quantification kit for Illumina platforms (Kapa Biosystems). [3] Based on the concentration, the libraries were equimolarly pooled and were sequenced on a NextSeq500 instrument with a NextSeq 500/550 High Output Kit V2.5 (75 cycles) using 5% PhiX without dark cycles (n = 11) or 10-20% PhiX with 7 dark cycles (protocol provided by Illumina, n = 49). A maximum of 12 samples were pooled in one sequencing run resulting in 19.05 million [17.05 – 21.72] single-end reads per sample on average (supplementary table 3).

### Sequencing quality control and mapping

After sequencing, bcl files were demultiplexed using bcl2fastq v2.19.1.403. The raw fastq files were first quality checked with FastQC v0.11.5^18^. During this check, many reads did not pass QC due to severe adaptor contamination. These adaptors were removed with Trim Galore v0.4.4^19^ (with --rrbs flag for RRBS data) and CutAdapt v1.16^20^ with default parameters except for “--three_prime_clip_R1 1”, and processed fastq files were again quality checked with FastQC. Mapping to GRCh37 was done with Bismark v0.19.0^21^ with default parameters. Mapping efficiency of the cf-RRBS samples was 50.25% [46.00 – 52.62]. Bisulfite conversion percentage was assessed by mapping to the lambda genome and was 98.05% [98.65 – 97.50]. The bait regions were defined as the MspI regions between 20-200 bp in GRCh37 (see ‘Feature Generation’) and the number of reads in these regions were counted with picard tools v2.18.27 (hsmetrics module; http://broadinstitute.github.io/picard/). The percentage of bases mapping to these “bait” regions was 87.43% [85.29 – 90.19]. Per-sample information is available in supplementary table 3. Visualizations were made with the R programming language v3.2.1. and ggplot2 v3.0.0.

### Copy number alteration profiling

For each sample, shallow whole genome sequencing (sWGS) data was available to assess copy number alterations (CNAs) in cfDNA. After DNA isolation, cfDNA was processed as previously described by Raman *et al*. ^22^. After sequencing, fastq files were mapped to GRCh38 with bwa 0.7.17 and duplicate reads were removed after mapping with picard tools v2.1.1. WisecondorX (https://github.com/CenterForMedicalGeneticsGhent/WisecondorX) with 400 kb binsize was used to call copy number variations. In addition, we used WisecondorX to detect CNA in the cf-RRBS data with 400 kb bins after mapping to a bisulfite converted GRCh38 reference genome. Duplicate reads were not removed in the cf-RRBS data in accordance with the Bismark user guide^21^. All samples were normalized with an in-house dataset from data obtained from healthy volunteers for both sWGS and cf- RRBS. Copy number profiles from all samples are available in supplementary data.

#### IchorCNA

To estimate the circulating tumor percentage in the cfDNA from the sWGS data, we used ichorCNA (https://github.com/broadinstitute/ichorCNA) with 500 kb binsize and the following parameters: “--chrs ‘c(1:22)’ --chrTrain ‘c(1:22)’ --scStates ‘c(1,3)’ --txnE 0.9999 --txnStrength 10000 --normal ‘c(0.2,0.35,0.5,0.65,0.8)’ --maxCN 5”.

### Building classifier with publicly available data

We used publicly available methylation profiling data from neuroblastoma (n = 220), osteosarcoma (n = 86), Wilms tumor (n = 131), clear cell sarcoma of the kidney (CCSK, n = 11) (TARGET, https://ocg.cancer.gov/programs/target), rhabdomyosarcoma^23^ (n = 53), Ewing sarcoma^12,24,25^ (n = 38), malignant rhabdoid tumors^26^ (MRT, n = 26), prepubertal white blood cells^27^ (WBCs, n = 52) and non-malignant cfDNA^15,28^ (n = 24) to build the reference set. The methylation signature of 2801 brain tumors was obtained from Capper *et al*.^12^.

Neuroblastoma cases from TARGET with unspecified *MYCN* status, and rhabdomyosarcoma cases with unspecified subgroup (embryonal/alveolar) were removed from the datamatrix (supplementary table 1).

Within these published datasets, the epigenetic profile of neuroblastoma, osteosarcoma, Wilms tumor, CCSK, WBC, rhabdomyosarcoma and Ewing sarcoma was determined with Infinium HumanMethylation450K and MRT samples were determined with Infinium HumanMethylationEPIC. For non-malignant controls by Chan *et al*.^28^ (n = 24), the authors used low-coverage whole genome bisulfite sequencing. Processing of raw fastq files from sequencing data was done similar as described above. WGBS files were deduplicated after mapping with Bismark, RRBS samples were not deduplicated. Accession numbers for published datasets are available in supplementary table 1. Chromosomes X and Y were removed from downstream analysis.

### Feature selection

CpGs were grouped in order to use more mappable reads. Input data (cf-RRBS test set & public reference set) was prepared similar to Kang *et al*. However, we adjusted the target regions to make them more suited for RRBS data. First, we used mkrrgenome^29^ to extract all MspI regions between 20-200 bp from GRCh37. Then, we merged all remaining regions within 1 bp from each other with BEDtools^30^. Finally, clusters were retained if they contain at least 3 CpGs covered on the Illumina HM450K array, resulting in 14,103 clusters covering 61,750 probes on the HM450K array. In our cohort, these regions were consistently covered across all samples, except for 8 regions that were not covered in any of the samples (supplementary data).

### t-SNE visualizations

Clustered and processed beta values from the 9 reference entities were grouped in a single data matrix with Python v3.6.3 and pandas v0.20.3. A t-SNE plot was generated on the full data matrix (634 samples and 4811 CpG clusters), after removing missing values (9292 CpG clusters, 65.88%) with sklearn v0.19.1 and matplotlib v2.1.0. The t-SNE parameters were n_components=2, perplexity=30, n_iter=2000. Mean sigma was 1.678887, error after 2000 iterations was 0.466.

### NNLS classification

Classification of plasma and CSF cfDNA samples was done with non-negative least squares (NNLS) matrix decomposition as described by Moss and colleagues^14^ (https://github.com/nloyfer/meth_atlas). The reference sets were identical as previously described (one reference set for intracranial and one reference set for extracranial tumor entities) and CpGs were grouped in 14,103 clusters as well for both reference and test samples. As normal class, the shallow whole genome bisulfite sequencing (sWGBS) cfDNA data from Chan *et al*. was used for both reference sets. For each tumor entity in the reference dataset, the median beta value of the CpG cluster was calculated, resulting in one column with beta values per tumor or normal entity. The sample class was determined by extracting the tumor entity with the highest fraction (after normal and WBC fractions) after running the meth_atlas NNLS wrapper. Samples with no estimated tumor fraction were labeled as inconclusive.

## Results

### The methylation signature is pediatric tumor type specific

Similar to the t-SNE plots as described by Capper *et al*.^12^, we applied t-SNE dimensionality reduction (Figure 1) to the extracranial reference dataset to assess the feasibility of establishing the diagnosis of extracranial pediatric cancers based on the methylation signature from public data. Separate clusters based on the histopathological diagnosis could be visually identified. Furthermore, Ewing sarcoma samples from three different public datasets cluster together, indicating that laboratory and/or method batch effects from different publicly available datasets are not dominating the classification. In addition, some individual tumor samples cluster with white blood cells (WBC). We hypothesize that either these biopsies may have been contaminated with leukocytes during surgery, or the tumor might have been rich in infiltrating lymphocytes. Nevertheless, to avoid bias, these samples were not excluded from the reference dataset.

**Figure 1:**
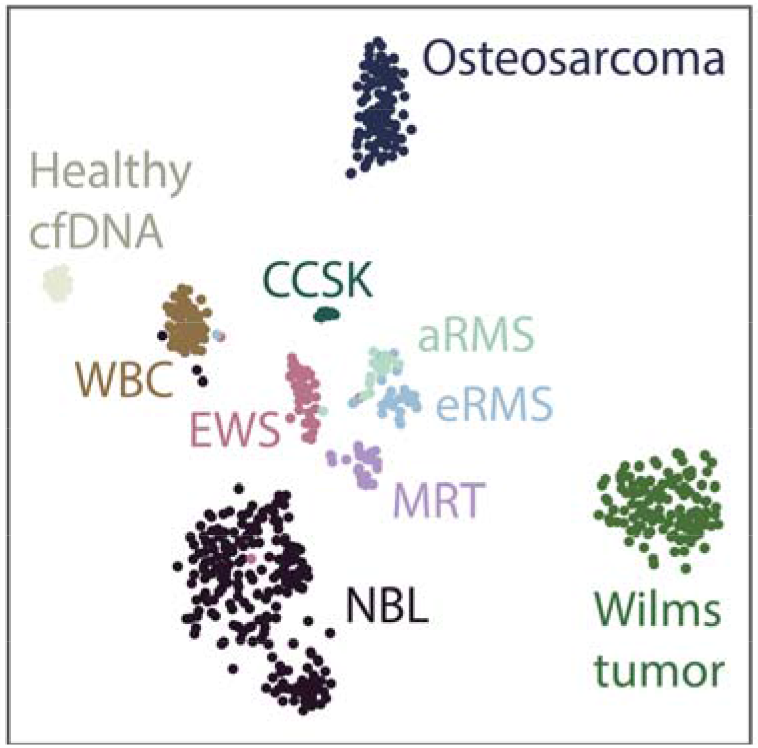
2-component t-SNE plot derived from the samples in the extracranial tumor reference dataset after grouping CpGs into 14,103 regions.

### Classification of cell-free DNA according to histopathological diagnosis

We profiled the methylation pattern of 59 pediatric cancer cases (60 samples), of which 56 plasma samples and 4 CSF samples and we were able to classify 49 samples (81.66%) correctly according to the histopathologic diagnosis.

Within the extracranial tumor entities, we could correctly classify neuroblastoma (n = 9/10), Wilms tumor (n = 13/16), rhabdomyosarcoma (n = 14/17), MRT (n = 1/1), CCSK (n = 1/2), Ewing sarcoma (n = 3/6) and osteosarcoma (n = 4/4) from the cfDNA methylation profile (Figure 2). Subclasses within the rhabdomyosarcoma group (i.e. alveolar and embryonal) are correct in 13 out of 14 of the correctly classified rhabdomyosarcoma cases. The two clear cell sarcoma samples derived from the same patient but sampled 24 hours apart misclassified once as an osteosarcoma sample with an estimated tumor fraction of 2.5%, and the second time correctly classified as clear cell sarcoma of the kidney with 2.5% estimated tumor fraction. In addition, we were able to distinguish medulloblastoma (n = 3/3) from an atypical teratoid-rhabdoid tumor using the methylation profile of CSF cfDNA.

**Figure 2:**
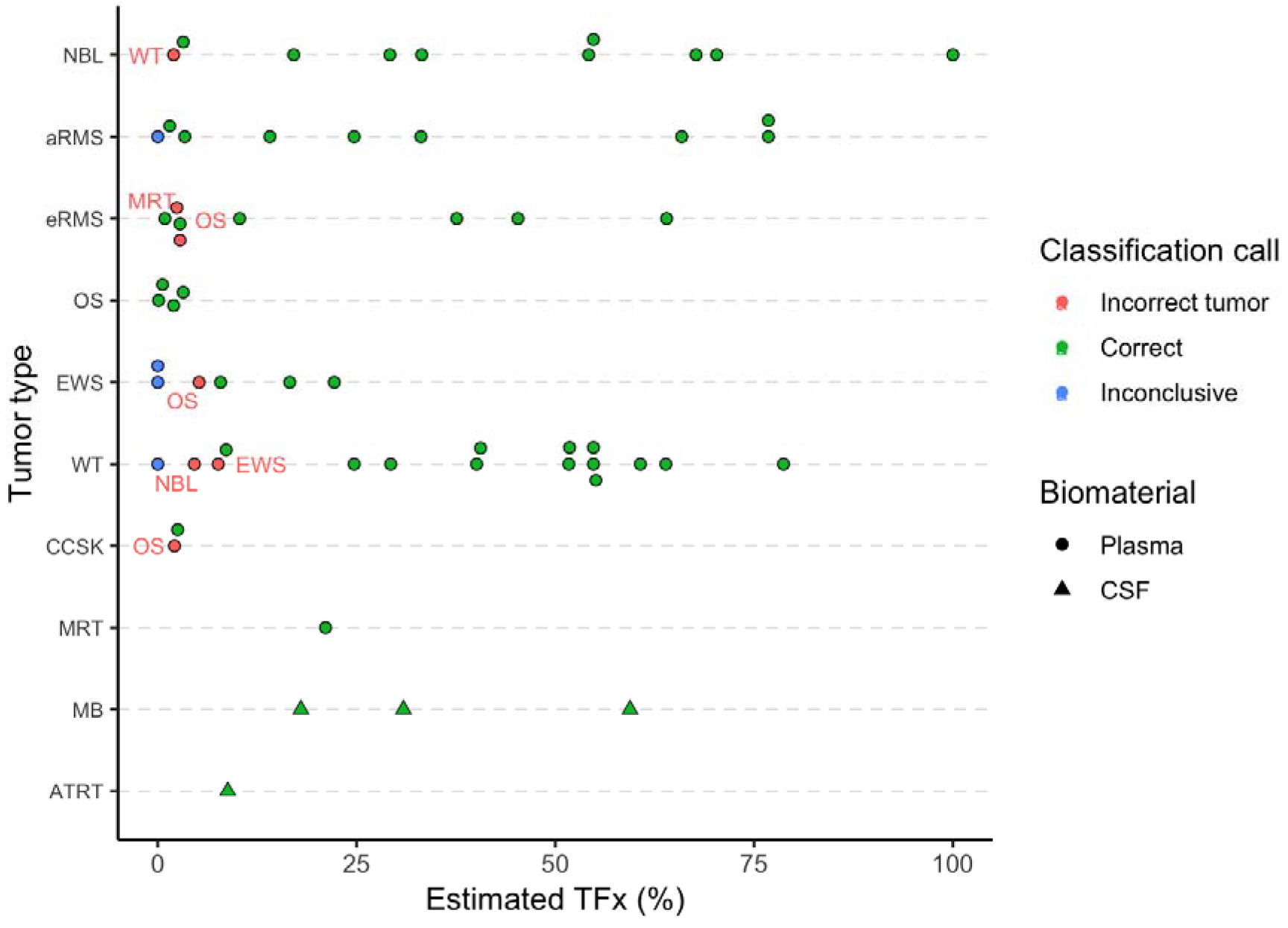
Overview of the results after classification based on the cfDNA methylation profile of the respective samples. Red labels near the dots on the scatter plot indicate what entity the sample is misclassified as. TFx, tumor fraction. Inconclusive indicates that no tumor fraction was detected in the cfDNA; NBL, neuroblastoma; aRMS, alveolar rhabdomyosarcoma; eRMS, embryonal rhabdomyosarcoma; OS, osteosarcoma; EWS, Ewing sarcoma; WT, Wilms tumor; CCSK, clear cell sarcoma of the kidney; MRT, malignant rhabdoid tumor; MB, medulloblastoma; ATRT, atypical teratoid-rhabdoid tumor.

### Classification accuracy is correlated with sample quality

Misclassified samples had significantly lower estimated tumor fraction in the cfDNA based on methylation data (2.10% [0.00 – 3.70] vs 30.90% [8.80 – 54.80], p < 0.001). Furthermore, if the estimated tumor fraction was above 10%, all samples classified correctly (n = 36). If the estimated tumor fraction was below 10%, 13 out of 24 samples (54.17%) classified correctly (Figure 5). To investigate whether a low tumor fraction correlates to disease extent, we grouped patients by metastatic status. Between these two groups, the estimated tumor fraction was not significantly different (30.90% [3.00 – 54.50] vs 16.60% [3.20 – 51.80], p = 0.39, Figure 5). Further investigation into the misclassified samples revealed that the majority of these samples had evidence for medium to high HMW contamination (cfDNA/HMW DNA ratio less than 5). In samples with low HMW DNA contamination, the classification accuracy was 94.59% (n = 35/37) (Figure 5). In samples with medium to high HMW DNA contamination, the classification accuracy was 60.86% (n = 14/23).

### Copy number analysis is feasible using cf-RRBS data

Copy number alterations could be detected with shallow WGS in 42 out of 60 samples (Figure 3, Figure 4). The concordance between estimated tumor fraction using methylation profiling data and the estimated tumor fraction by ichorCNA analysis of sWGS profiles was moderate for samples with low to medium HMW DNA contamination (Spearman r 0.77 and 0.78, respectively, Figure 6). Importantly, *MYCN* amplification could be observed with both the sWGS and cf-RRBS CNA profiles in the three neuroblastoma samples with confirmed *MYCN* amplification. If CNAs were present in cf-RRBS (n = 32), the correlation of the CNA profile with sWGS was high (0.87 [0.82 - 0.94]). However, the reverse was not true; several samples (n = 10) showed CNAs according to sWGS but none using cf-RRBS. In the group with discordant CNA profiles between cf-RRBS and sWGS, 5 out of 10 samples (50%) had evidence for high HMW contamination. In contrast, in the group showing CNA with both methods, only 4 out of 46 samples (8.69%) had evidence for high HMW DNA contamination (p = 0.004, Fisher’s Exact Test). None of the 18 (out of 60) samples that showed no CNA using sWGS had evidence for CNA based on cf-RRBS data (Figure 6).

**Figure 3:**
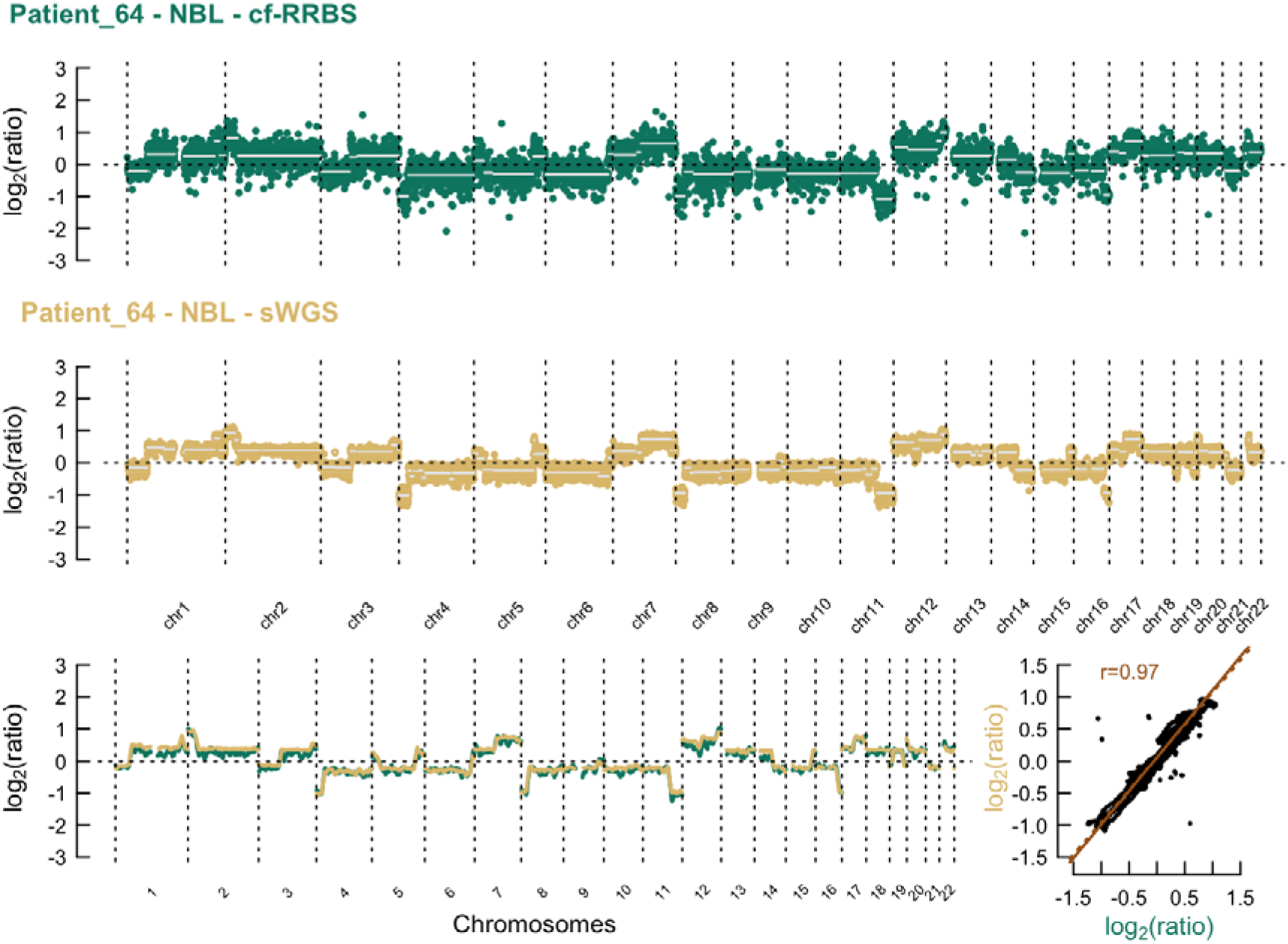
**Top & middle:** Example of the comparison of copy number profiles derived from (top) cf-RRBS data and (middle) sWGS data with 400 kb binsize of a high-quality sample. Lower left: sliding window of 10 Mb average log2 ratio. **Lower right:** scatterplot between cf-RRBS and sWGS with Pearson r. The dotted line equals least squares fit; the solid line equals the orthogonal regression fit. Cf-RRBS, cell-free reduced representation bisulfite sequencing; sWGS, shallow whole genome sequencing.

**Figure 4:**
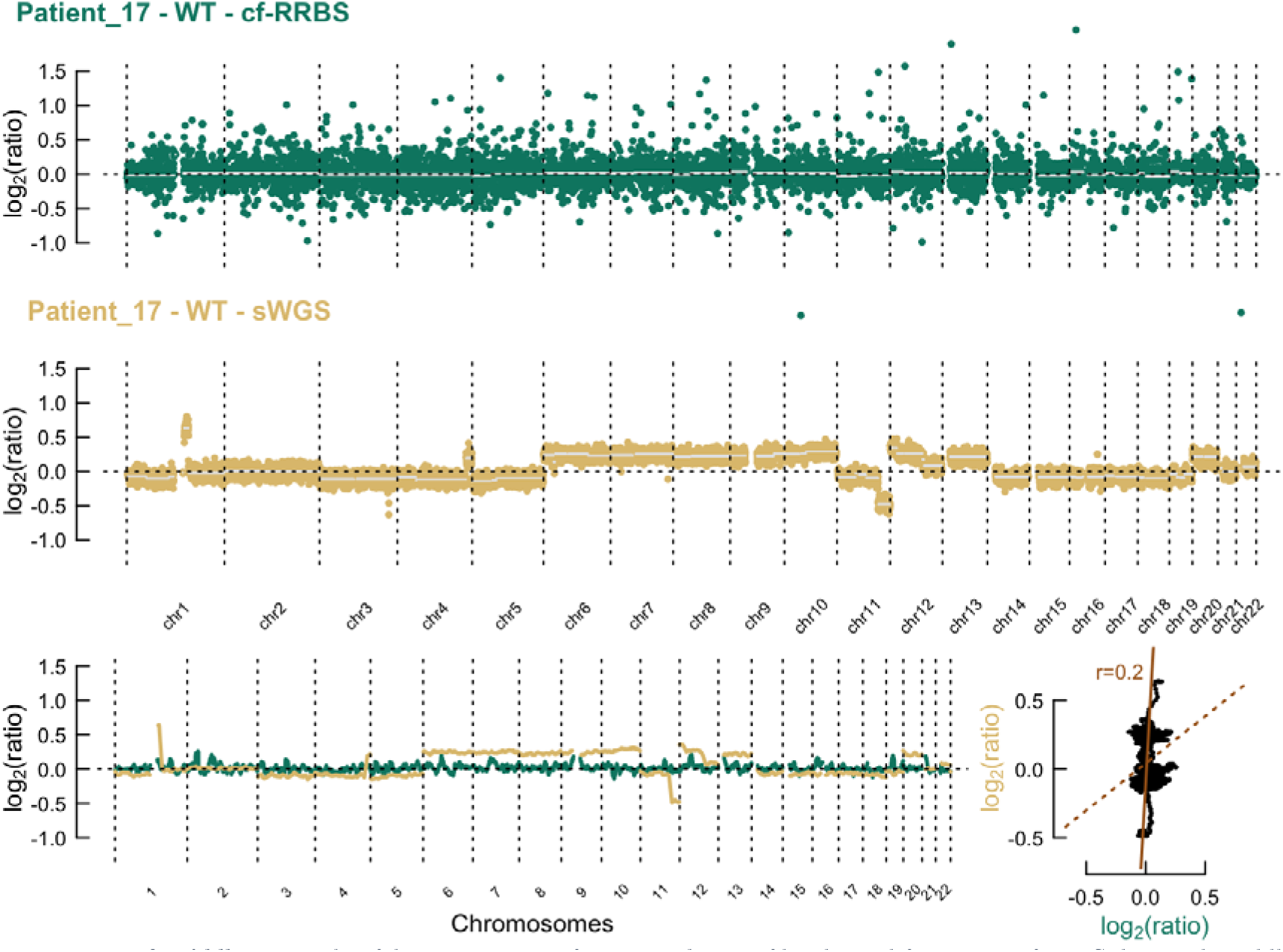
**Top & middle:** Example of the comparison of copy number profiles derived from (top) cf-RRBS data and (middle) sWGS data with 400 kb binsize of a low-quality sample. Lower left: sliding window of 10 Mb average log2 ratio. **Lower right:** scatterplot between cf-RRBS and sWGS with Pearson r. The dotted line equals least squares fit; the solid line equals the orthogonal regression fit. Cf-RRBS, cell-free reduced representation bisulfite sequencing; sWGS, shallow whole genome sequencing.

**Figure 5:**
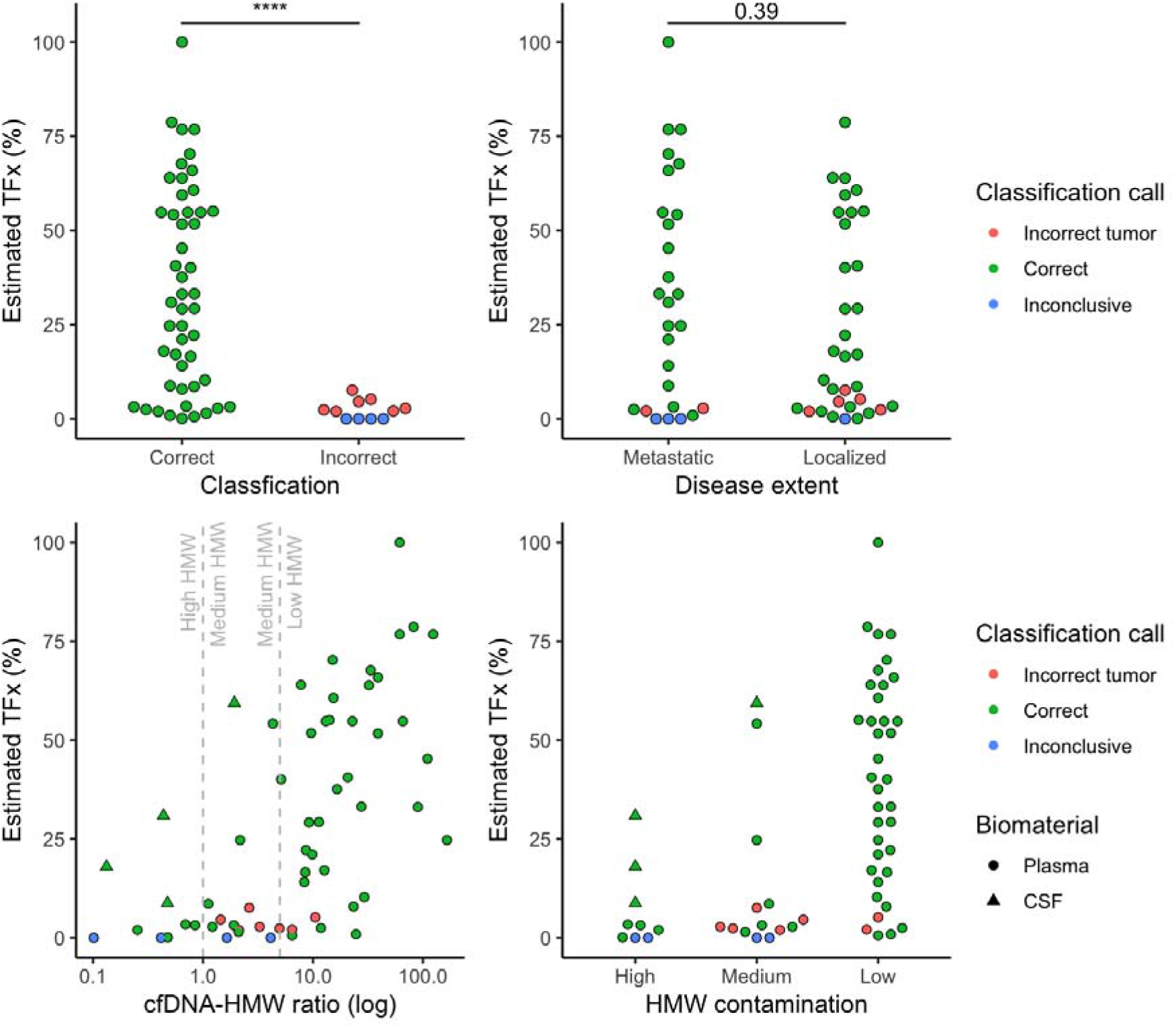
**Top left:** the estimated tumor fraction is significantly lower in the misclassified group (p < 0.001, Wilcoxon rank sum test)). **Top right:** The estimated tumor fraction does not differ significantly for disease extent across all tumor types (p > 0.05, Wilcoxon rank sum test). **Bottom left:** The majority of the misclassifications are situated in the medium and high HMW group (cfDNA/HMW ratio less than 5). **Bottom right:** In the group with low HMW DNA contamination, classification accuracy reaches 94% (n = 35/37), indicating that pre-analytical variables or sample quality might influence the classification accuracy. TFx, tumor fraction; HMW, high molecular weight; cfDNA, cell-free DNA; CSF, cerebrospinal fluid; **** = p < 0.001.

**Figure 6:**
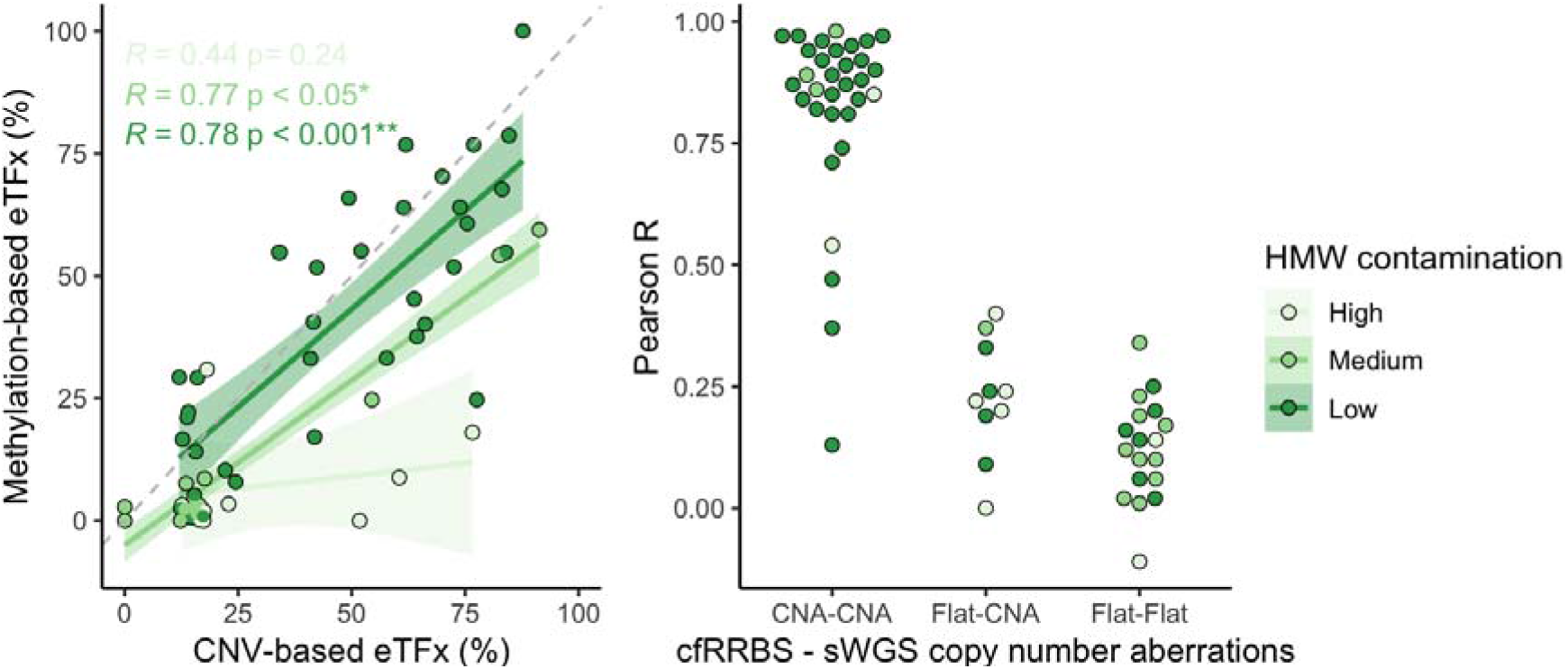
**Left:** scatterplot of the estimated tumor fraction derived from cf-RRBS data vs. estimated tumor fraction derived from sWGS data. In samples with low to medium HMW contamination, Spearman correlation between methylation and CNA-based tumor fraction estimation is moderate to high (r = 0.78 and 0.77, respectively). Solid line equals the linear regression fit. Dashed line equals y = x. **Right:** if CNAs are present in both cf-RRBS and sWGS data, Pearson correlation between both methods is high. “Flat” indicates copy neutral (i.e. no gains or losses) samples. eTFx, estimated tumor fraction; CNV, copy number variations; CNA, copy number aberrations, HMW contamination, high molecular weight DNA contamination; **** = p < 0.001.

## Discussion

We show that classification based on cfDNA methylation profile as determined by cf-RRBS is feasible for the subset of extra-and intracranial pediatric cancer entities included in this study. Overall, we are able to correctly classify 49 out of 60 (81.6%) samples. We are able to identify specific recurrent genomic alterations such as *MYCN* amplification in neuroblastoma and differentiate between alveolar and embryonal rhabdomyosarcoma subtype from these same data. Importantly, only 1-10 ng of cfDNA is required for the cf-RRBS analysis, making clinical implementation in pediatric practice feasible from just 200 μl of plasma. Furthermore, cost-per-sample was calculated at a conservative 180 euro, including sample collection, DNA isolation, library preparation and sequencing (excluding personnel costs), favoring our techniques routine application over the substantially more expensive gold-standard of whole genome bisulfite sequencing.

In our study, misclassifications seemed to be predominantly associated with sample quality. Samples of low quality had a higher amount of high molecular weight, non-cell-free DNA and this artificially reduced the estimated tumor fraction, resulting in a more challenging classification. Inherent to the retrospective multicenter nature of our study is a lack of control over the pre-analytical variables such as the blood tube used, the handling of the collection tube, any delay between collection and downstream processing and the SOP used in cfDNA isolation. Preservation tubes (e.g. Streck Cell-Free DNA BCT, Streck) have been developed to stabilize the cfDNA and avoid high molecular weight DNA contamination. However, it is currently unknown to what extent the preservation medium in these tubes alters the methylation profile of the cfDNA or hampers its downstream processing for cf-RRBS. Furthermore, classification accuracy may be improved for samples with a very low ctDNA fraction given a more sensitive computational deconvolution algorithm, a more robust reference dataset or a combination of genomic and epigenomics-based classification. Interestingly, in our study, the estimated ctDNA fraction was not associated with extent of disease. This is in contrast to the adult oncology field, where it has often been suggested that cfDNA based assays are mainly of value in high stage, metastatic disease^31^. This may depend on the tumor entity, but should further be explored in future, prospective studies. Furthermore, we could correctly classify 9 out of 18 samples with a neutral cfDNA CNA profile. This hints at an improved sensitivity of methylation profiling over CNA profiling in cfDNA samples with a low tumor fraction.

Other efforts exploring cell-free DNA in pediatric cancer have focused on detection of copy number aberrations, structural variants^32,33^ and mutations^34^. However, these alterations come with low specificity for classification of pediatric solid tumors according to their histopathological diagnosis^35^. In addition, there is a high interest to develop a screening test based on cfDNA methylation in adult oncology^36^. Due to the rarity of cancer in children, we do not see our test as a screening tool, but rather as a complementary diagnostic test for children for whom a classic biopsy is unfeasible or of low quality. Indeed, it could improve the diagnosis of tumor types with very similar histology (e.g. small round blue cell tumors) as illustrated by the distinction of MRT, CCSK and Wilms tumor in this study. This could potentially impact on clinical decision making for patients with renal tumors who are usually not biopsied before therapy start. Another major application is in the minimally-invasive classification of brain tumors, where tumor tissue methylation profiling is being pitched as a way to improve tumor classification^11,12^ but high quality and sufficiently large biopsies are hard to obtain.

While we provide proof-of-concept for the applicability of the cf-RRBS assay, there are several limitations to our study. Building a classifier from public datasets from many different laboratories will result in batch and platform effects that will inevitably bias the results of this study. In addition, our current classifier is also limited to entities of whom public data is available. The public data available was generated with different strategies (array, RRBS, WGBS) potentially resulting in a suboptimal classifier. Ideally, a reference atlas of RRBS or WGBS data from a wide array of pediatric cancers and healthy children should be generated to further improve our classifier. Implementation into clinical trials or routine diagnostics will require additional adaptations to our method. In future validation studies, the inclusion of children without cancer will be necessary to calculate the specificity of the assay. Furthermore, in its current iteration our assay has a theoretical turn-around-time of 5 working days which is too long for its use in real-time clinical decision making.

In recent years, a number of alternatives to WGBS for cfDNA methylation profiling were developed. The Roche SeqCap Epi assay is a capture-based technique and is able to assess the methylation profile of cfDNA by enriching for CpG dense genomic areas. However, this technique has a lower cost-effectiveness compared to cf-RRBS, is difficult to implement in a routine clinical setting and the protocol is cumbersome and time-consuming^17^. Single cell RRBS (scRRBS) has also been suggested as a cost-effective alternative to classical RRBS and WGBS for low-input samples, but is more labor-intensive^15^. More recently, cfMeDIP-seq^37^ was used to detect and classify adult cancers in an early stage. However, as this method is based on immunoprecipitation it has its own drawbacks^38^. These techniques all cover different parts of the methylome, and the optimal technique for sensitive tumor detection remains yet to be demonstrated.

In conclusion, while a thorough prospective evaluation is required to delineate and establish the true value of the technique and its clinical application, this proof-of-concept study suggests a role for cf-RRBS as a cost-effective addition to the diagnostic toolbox in pediatric oncology.

## Acknowledgements

A.D.K and R.V.P were funded by a predoctoral fellowship from the Research Foundation Flanders (FWO). This project was partially funded by Kom op tegen Kanker, a CRIG (Cancer Research Institute Ghent) Young Investigators Proof-of-Concept grant, vzw Kinderkankerfonds - a non-profit childhood cancer foundation under Belgian law (grant to T.L) and by the VIB Grand Challenges Program (N.C. lab). We thank Lennart Raman for assistance with WisecondorX and Peter Stockwell for assistance with mkrrgenome. We thank the Center for Medical Genetics Ghent lab technicians for assistance with plasma preparation, cfDNA extraction, shallow WGS library preparation and sequencing (Thalia Van Laethem, Eveline Vanden Eynde, Kimberly Verniers, Peter Degrave, Dimitri Broucke, Machteld Baetens, Aline Eggermont, Els De Smet). We thank Landric Vautmans for assistance with the FEMTO Pulse. In France, this study was supported by the Annenberg Foundation, the Association Hubert Gouin Enfance et Cancer, the Association Enfants et Santé, and the SiRIC programme (Site de Recherche Integrée sur le Cancer). This study makes use of data generated by dr. Junko Takita, Department of Pediatrics, The University of Tokyo and The Chinese University of Hong Kong (CUHK) Circulating Acids Research Group, as reported by Chan KC in Proc Natl Acad Sci USA (2013 Nov 19;110(47):18761-8), the TARGET database, Gene Expression Omnibus (GEO) identifiers GSE89041, GSE97529, GSE90496, GSE107946, GSE90496 and ArrayExpress identifier E-MTAB-4187.

## Code and data availability

Scripts and supporting files are available at https://github.com/rmvpaeme/cfRRBS_manuscript. Raw data is available at EGA identifier EGAD00001005928. Processed data is available at ArrayExpress identifier E-MTAB-8770. The cf-RRBS protocol can be found at https://www.protocols.io/private/4098389D37C151C92AA19EC574AD9201.

Supplementary figures depicting library preparation and sequencing quality control metrics, as well as classification results and comparative copy number plots are available at https://github.com/rmvpaeme/cfRRBS_manuscript/blob/master/Markdowns/Analysis_paper_public.html.

