## Supplementary figures for "Minimally invasive classification of pediatric solid tumors using reduced representation bisulfite sequencing of cell-free DNA: a proof-of-principle study"

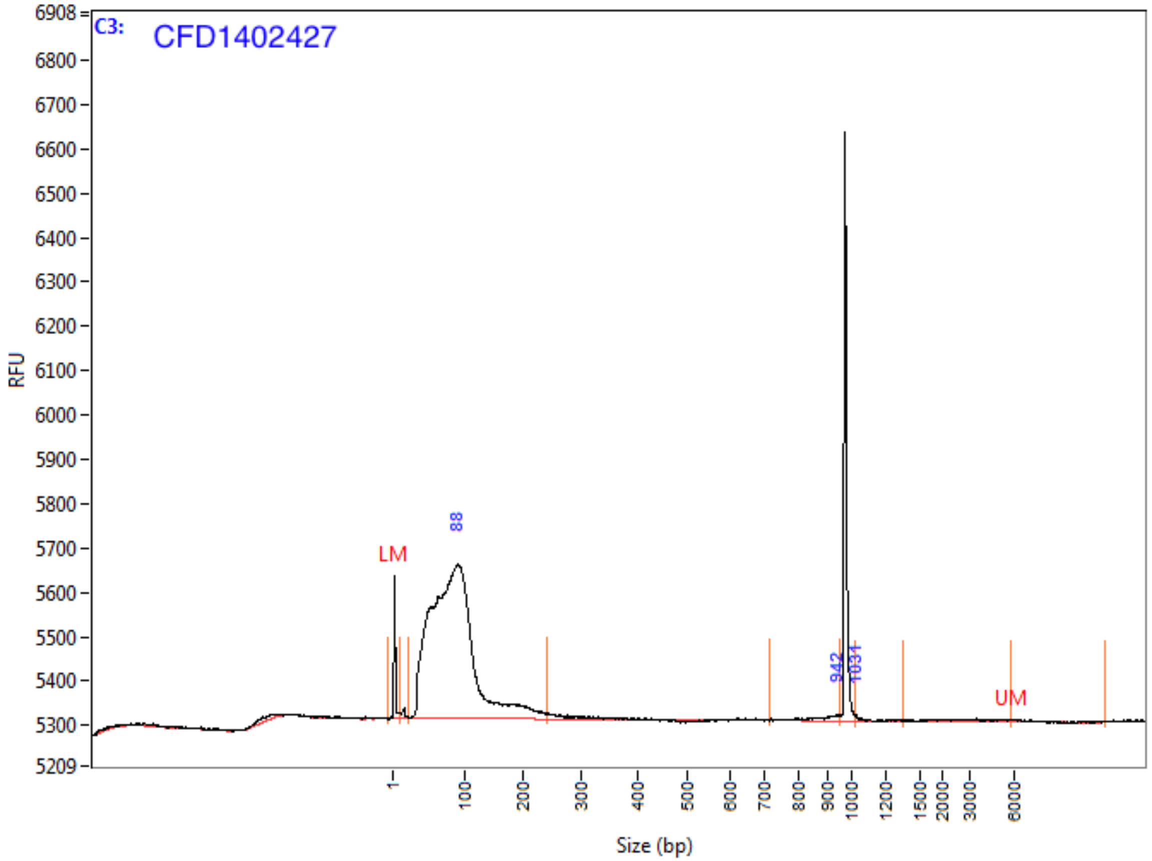

**Peak Table**

| | Size (bp) | ng/uL | % (Conc.) (ng/uL) | nmole/L | From (bp) | To (bp) | Avg. Size | CV% |
|---|---|---|---|---|---|---|---|---|
| 1 | 1 (LM) | 0.0004 | | 0.5401 | 0 | 10 | 1 | 170.25 |
| 2 | 88 | 0.1046 | 99.1 | 2.0445 | 23 | 241 | 84 | 43.91 |
| 3 | 942 | 0.0008 | 0.7 | 0.0015 | 713 | 952 | 877 | 6.83 |
| 4 | 1031 | 0.0002 | 0.2 | 0.0003 | 1025 | 1357 | 1118 | 9.82 |
| 5 | 6000 (UM) | 0.0000 | | 0.0000 | 5861 | 12258 | 8086 | 26.03 |
| | TIC: | 0.1056 | ng/uL | | | | | |
| | TIM: | 2.0463 | nmole/L | | | | | |
| | Total Conc.: | 0.1193 | ng/uL | | | | | |

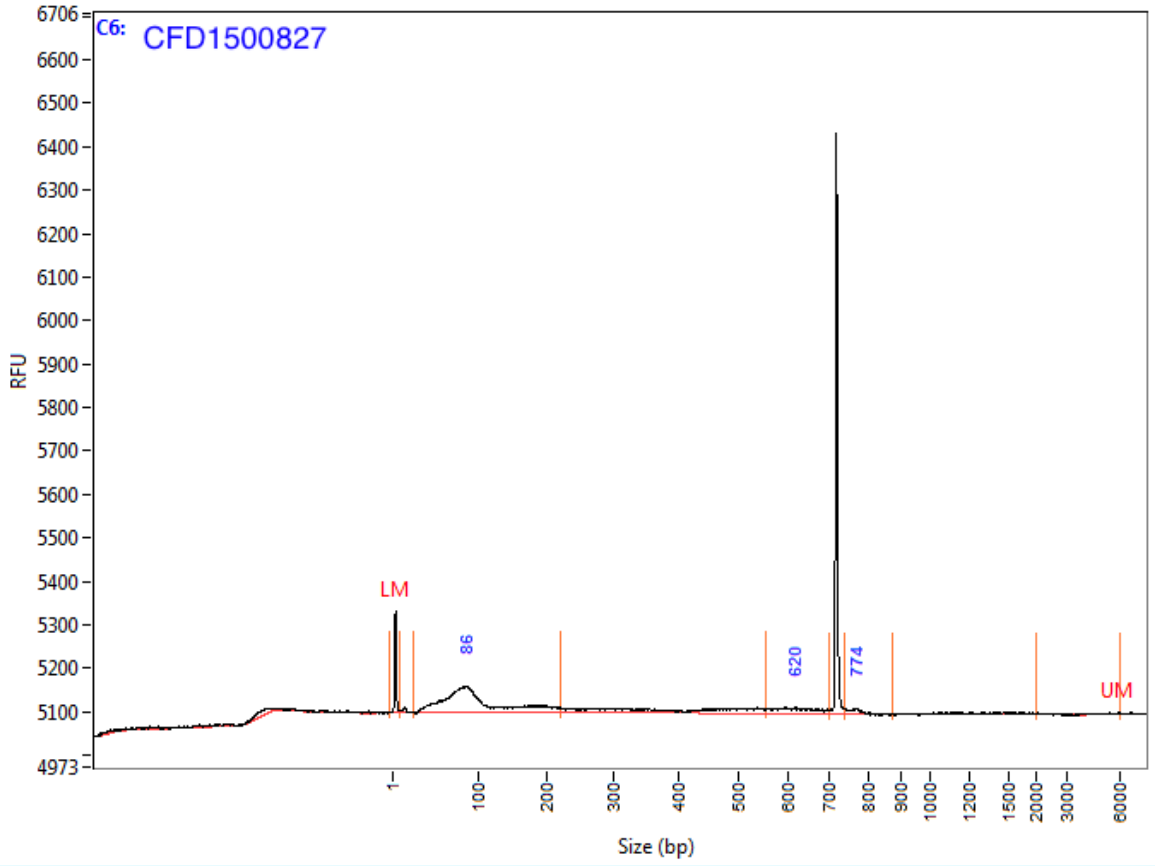

**Peak Table**

| | Size (bp) | ng/uL | % (Conc.) (ng/uL) | nmole/L | From (bp) | To (bp) | Avg. Size | CV% |
|---|---|---|---|---|---|---|---|---|
| 1 | 1 (LM) | 0.0004 | | 0.5401 | 0 | 9 | 1 | 189.44 |
| 2 | 86 | 0.0246 | 89.3 | 0.4073 | 23 | 220 | 99 | 46.41 |
| 3 | 620 | 0.0024 | 8.7 | 0.0064 | 555 | 702 | 622 | 6.60 |
| 4 | 774 | 0.0005 | 2.0 | 0.0012 | 740 | 874 | 763 | 1.69 |
| 5 | 6000 (UM) | 0.0000 | | 0.0000 | 2027 | 6115 | 4471 | 26.32 |
| | TIC: | 0.0275 | ng/uL | | | | | |
| | TIM: | 0.4148 | nmole/L | | | | | |
| | Total Conc.: | 0.0483 | ng/uL | | | | | |

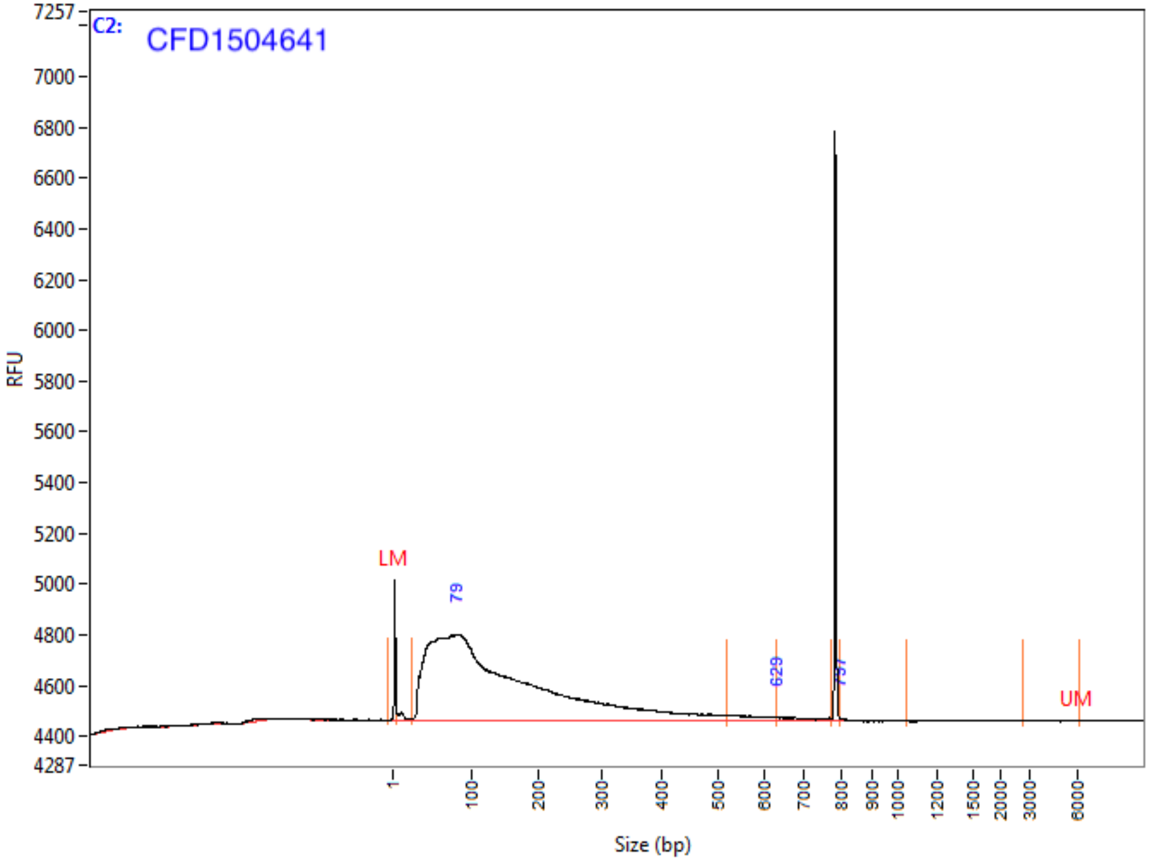

**Peak Table**

| | Size (bp) | ng/uL | % (Conc.) (ng/uL) | nmole/L | From (bp) | To (bp) | Avg. Size | CV% |
|---|---|---|---|---|---|---|---|---|
| 1 | 1 (LM) | 0.0004 | | 0.5401 | 0 | 5 | 1 | 174.44 |
| 2 | 79 | 0.1396 | 99.4 | 1.6381 | 24 | 517 | 140 | 71.85 |
| 3 | 629 | 0.0008 | 0.6 | 0.0019 | 629 | 775 | 690 | 6.27 |
| 4 | 797 | 0.0000 | 0.0 | 0.0001 | 797 | 1043 | 806 | 0.90 |
| 5 | 6000 (UM) | 0.0000 | | 0.0000 | 2818 | 6162 | 5307 | 23.25 |
| | TIC: | 0.1404 | ng/uL | | | | | |
| | TIM: | 1.6401 | nmole/L | | | | | |
| | Total Conc.: | 0.1514 | ng/uL | | | | | |

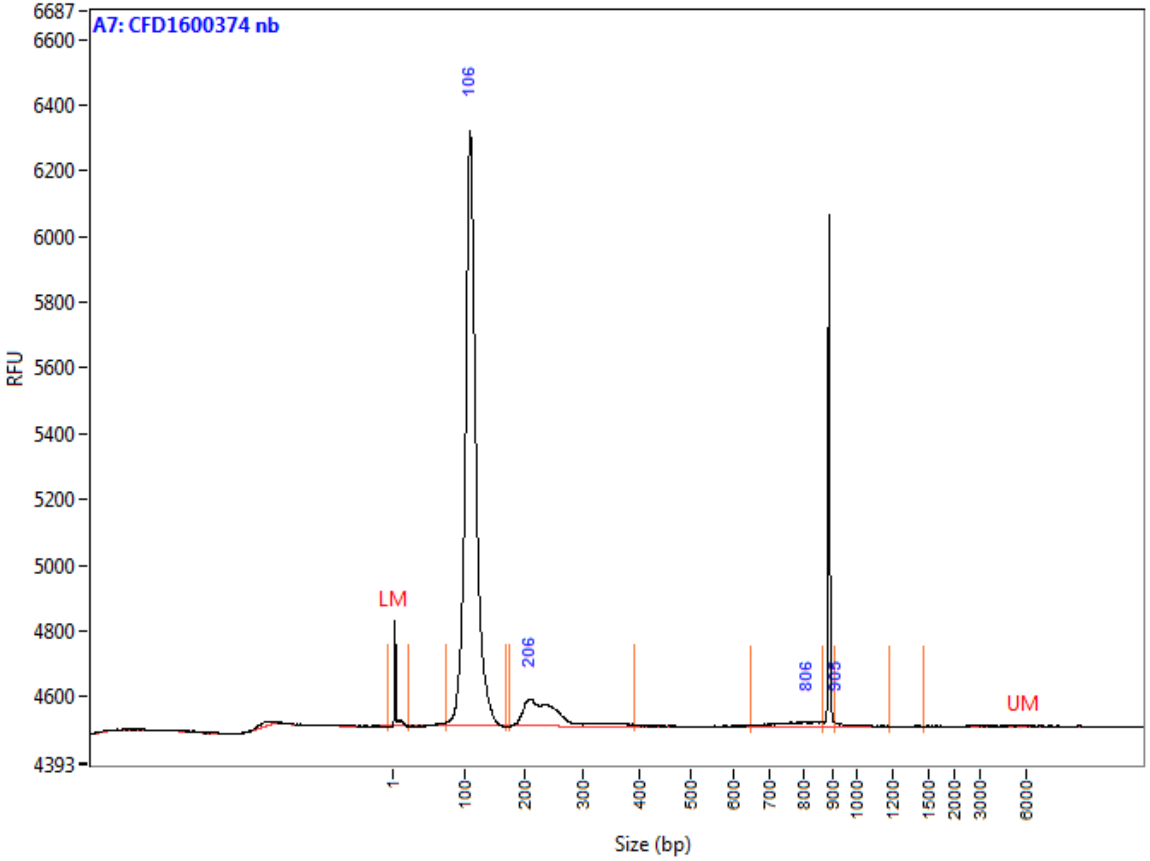

**Peak Table**

| | Size (bp) | ng/uL | % (Conc.) (ng/uL) | nmole/L | From (bp) | To (bp) | Avg. Size | CV% |
|---|---|---|---|---|---|---|---|---|
| 1 | 1 (LM) | 0.0007 | | 0.5202 | 0 | 22 | 2 | 181.14 |
| 2 | 106 | 0.2281 | 88.3 | 3.5005 | 72 | 167 | 107 | 9.72 |
| 3 | 206 | 0.0265 | 10.2 | 0.1836 | 172 | 392 | 237 | 17.41 |
| 4 | 806 | 0.0031 | 1.2 | 0.0067 | 647 | 864 | 774 | 7.19 |
| 5 | 905 | 0.0007 | 0.3 | 0.0012 | 905 | 1179 | 961 | 5.50 |
| 6 | 6000 (UM) | 0.0001 | | 0.0000 | 1462 | 14059 | 6082 | 49.49 |
| | TIC: | 0.2584 | ng/uL | | | | | |
| | TIM: | 3.6920 | nmole/L | | | | | |
| | Total Conc.: | 0.2771 | ng/uL | | | | | |

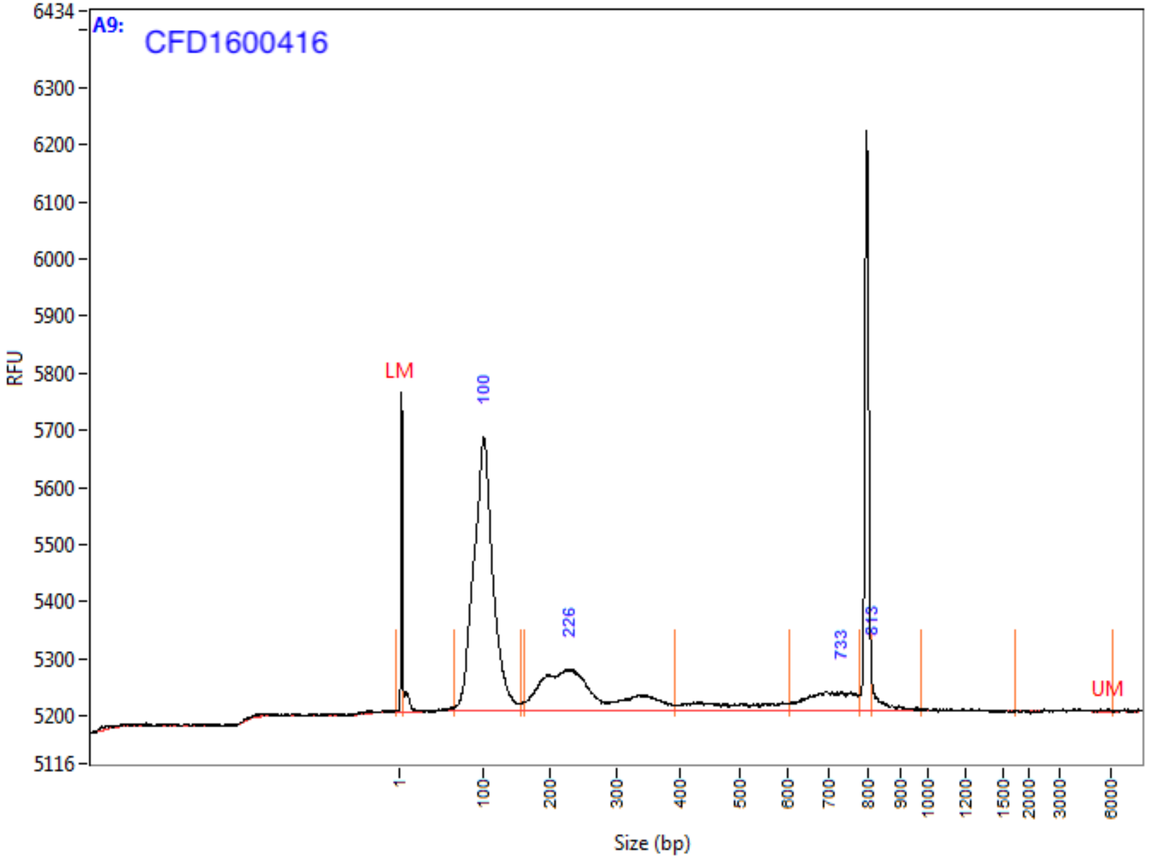

**Peak Table**

| | Size (bp) | ng/uL | % (Conc.) (ng/uL) | nmole/L | From (bp) | To (bp) | Avg. Size | CV% |
|---|---|---|---|---|---|---|---|---|
| 1 | 1 (LM) | 0.0011 | | 1.4680 | 0 | 5 | 1 | 179.29 |
| 2 | 100 | 0.1060 | 67.2 | 1.7231 | 66 | 157 | 101 | 14.17 |
| 3 | 226 | 0.0402 | 25.5 | 0.2708 | 162 | 390 | 244 | 23.05 |
| 4 | 733 | 0.0093 | 5.9 | 0.0218 | 605 | 780 | 700 | 6.70 |
| 5 | 813 | 0.0023 | 1.4 | 0.0044 | 813 | 980 | 852 | 4.58 |
| 6 | 6000 (UM) | 0.0000 | | 0.0000 | 1758 | 6169 | 5091 | 18.00 |
| | TIC: | 0.1577 | ng/uL | | | | | |
| | TIM: | 2.0200 | nmole/L | | | | | |
| | Total Conc.: | 0.1911 | ng/uL | | | | | |

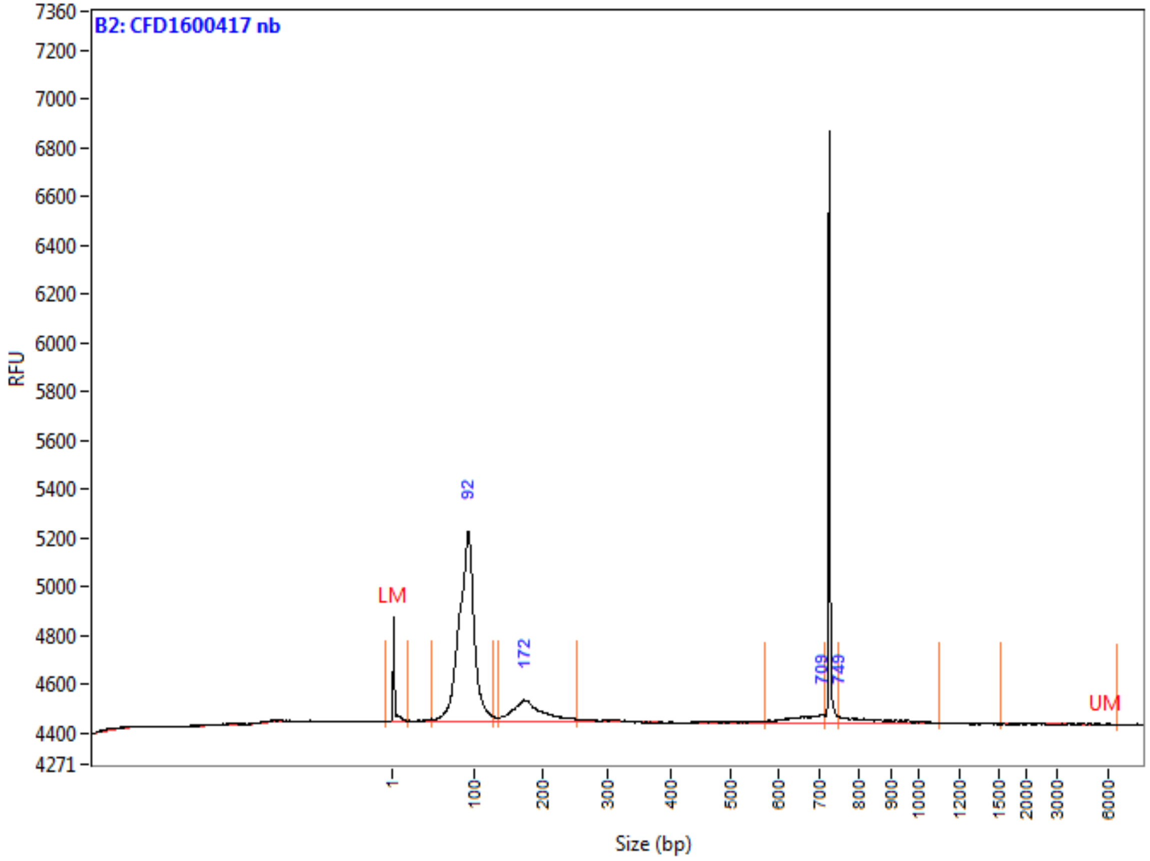

**Peak Table**

| | Size (bp) | ng/uL | % (Conc.) (ng/uL) | nmole/L | From (bp) | To (bp) | Avg. Size | CV% |
|---|---|---|---|---|---|---|---|---|
| 1 | 1 (LM) | 0.0005 | | 0.3937 | 0 | 19 | 2 | 188.49 |
| 2 | 92 | 0.0759 | 78.8 | 1.4165 | 50 | 129 | 88 | 12.25 |
| 3 | 172 | 0.0144 | 14.9 | 0.1301 | 136 | 253 | 182 | 14.14 |
| 4 | 709 | 0.0032 | 3.3 | 0.0080 | 571 | 715 | 659 | 5.77 |
| 5 | 749 | 0.0028 | 2.9 | 0.0054 | 749 | 1102 | 851 | 9.45 |
| 6 | 6000 (UM) | 0.0000 | | 0.0000 | 1538 | 6596 | 5328 | 14.98 |
| | TIC: | 0.0964 | ng/uL | | | | | |
| | TIM: | 1.5600 | nmole/L | | | | | |
| | Total Conc.: | 0.1133 | ng/uL | | | | | |

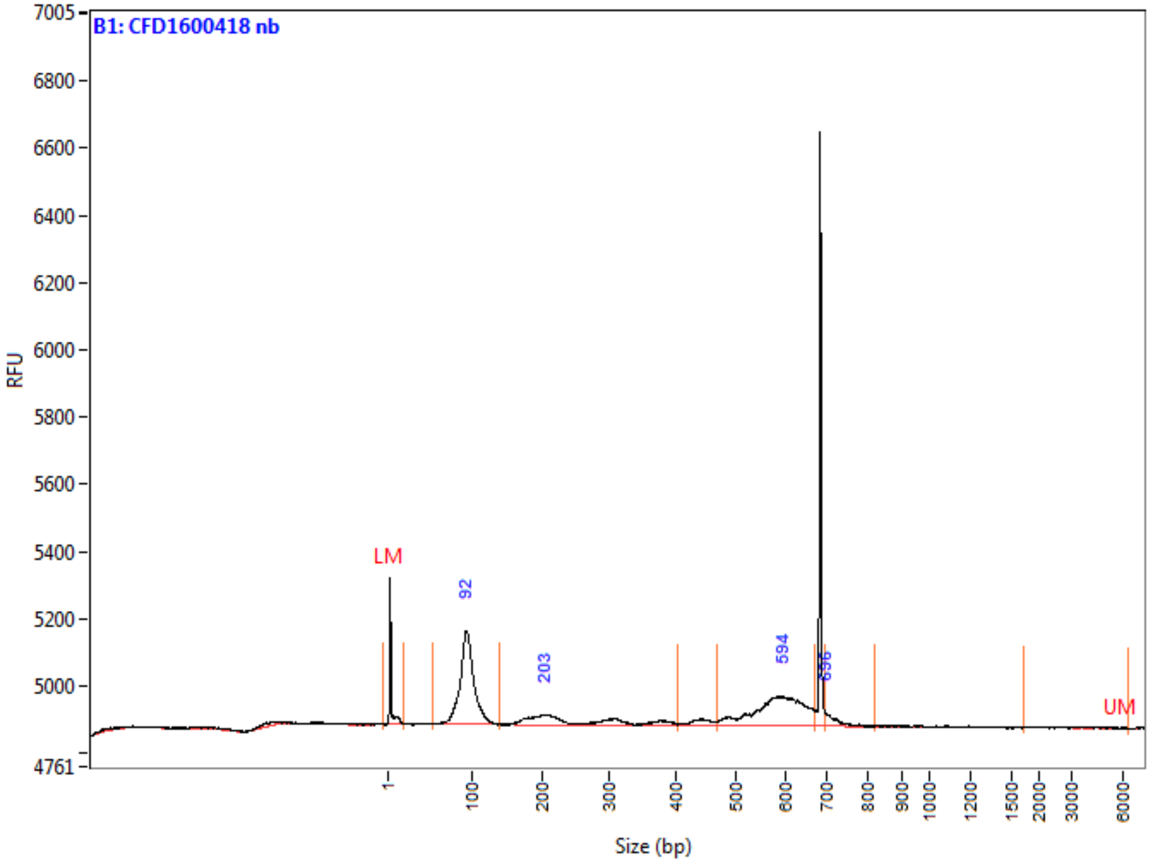

**Peak Table**

| | Size (bp) | ng/uL | % (Conc.) (ng/uL) | nmole/L | From (bp) | To (bp) | Avg. Size | CV% |
|---|---|---|---|---|---|---|---|---|
| 1 | 1 (LM) | 0.0005 | | 0.3937 | 0 | 18 | 2 | 159.35 |
| 2 | 92 | 0.0232 | 50.0 | 0.4135 | 53 | 140 | 92 | 10.62 |
| 3 | 203 | 0.0072 | 15.6 | 0.0473 | 140 | 403 | 251 | 27.75 |
| 4 | 594 | 0.0147 | 31.7 | 0.0416 | 469 | 671 | 582 | 8.47 |
| 5 | 696 | 0.0012 | 2.7 | 0.0029 | 696 | 822 | 720 | 2.98 |
| 6 | 6000 (UM) | 0.0000 | | 0.0000 | 1736 | 6344 | 4714 | 27.40 |
| | TIC: | 0.0464 | ng/uL | | | | | |
| | TIM: | 0.5053 | nmole/L | | | | | |
| | Total Conc.: | 0.0582 | ng/uL | | | | | |

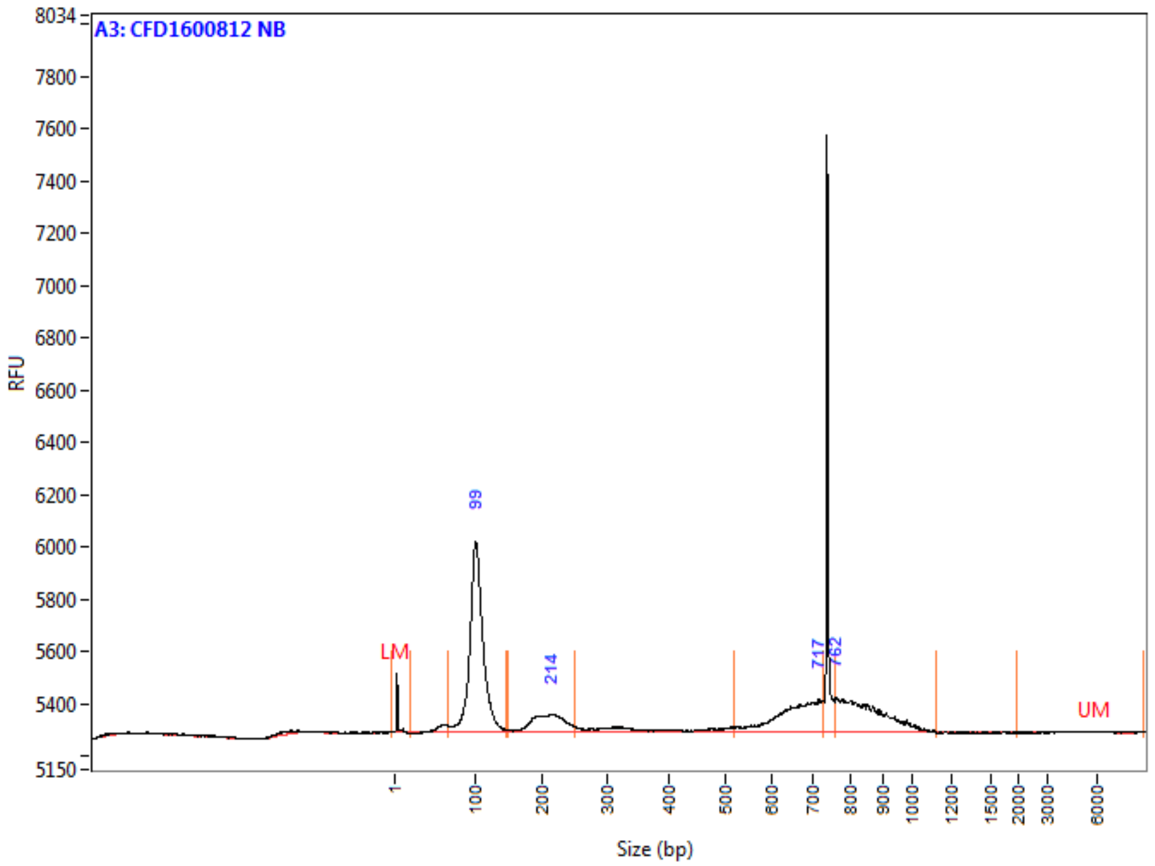

**Peak Table**

| | Size (bp) | ng/uL | % (Conc.) (ng/uL) | nmole/L | From (bp) | To (bp) | Avg. Size | CV% |
|---|---|---|---|---|---|---|---|---|
| 1 | 1 (LM) | 0.0003 | | 0.3525 | 0 | 19 | 1 | 168.54 |
| 2 | 99 | 0.0623 | 56.7 | 1.0223 | 66 | 146 | 100 | 11.00 |
| 3 | 214 | 0.0124 | 11.3 | 0.0986 | 149 | 250 | 206 | 10.52 |
| 4 | 717 | 0.0174 | 15.8 | 0.0438 | 518 | 727 | 652 | 7.95 |
| 5 | 762 | 0.0179 | 16.3 | 0.0346 | 759 | 1126 | 850 | 8.23 |
| 6 | 6000 (UM) | 0.0000 | | 0.0000 | 1987 | 9002 | 6735 | 16.89 |
| | TIC: | 0.1098 | ng/uL | | | | | |
| | TIM: | 1.1994 | nmole/L | | | | | |
| | Total Conc.: | 0.1338 | ng/uL | | | | | |

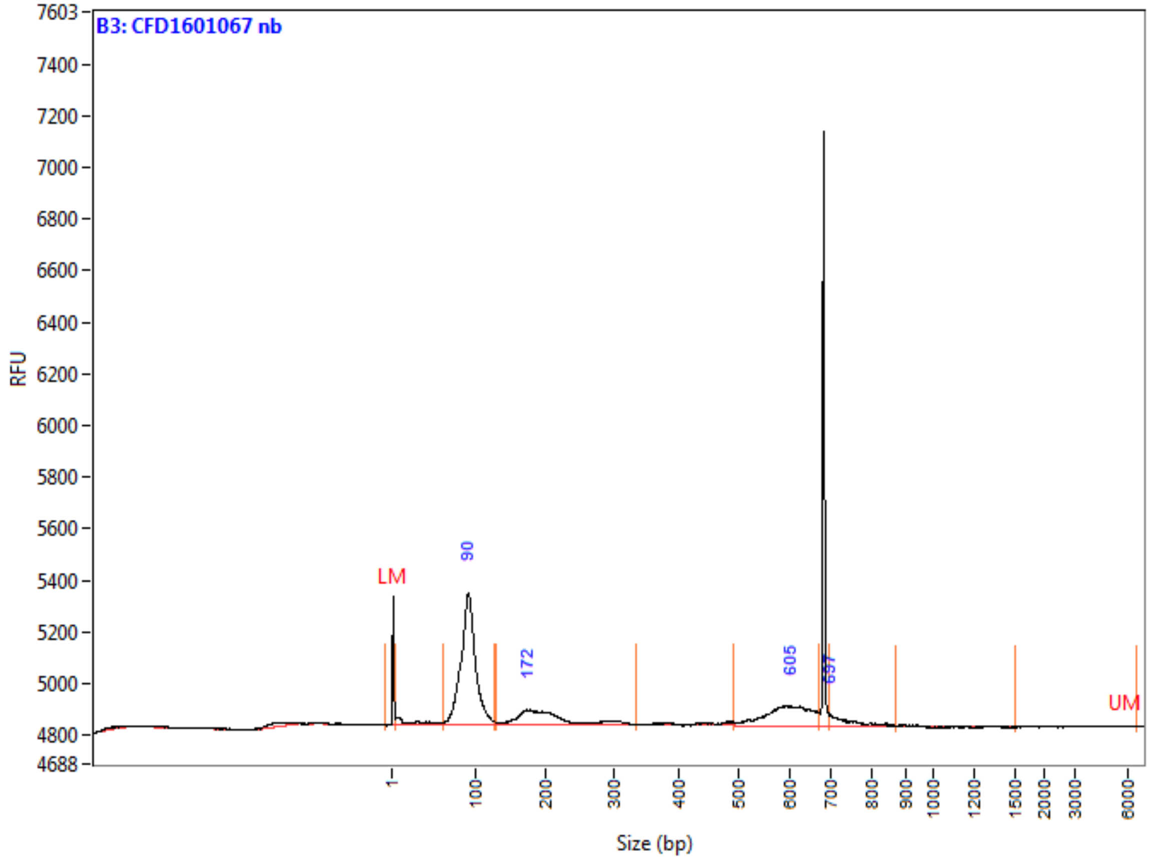

**Peak Table**

| | Size (bp) | ng/uL | % (Conc.) (ng/uL) | nmole/L | From (bp) | To (bp) | Avg. Size | CV% |
|---|---|---|---|---|---|---|---|---|
| 1 | 1 (LM) | 0.0005 | | 0.7062 | 0 | 4 | 1 | 191.91 |
| 2 | 90 | 0.0485 | 63.1 | 0.8942 | 62 | 128 | 89 | 11.23 |
| 3 | 172 | 0.0139 | 18.1 | 0.1124 | 127 | 336 | 203 | 22.12 |
| 4 | 605 | 0.0122 | 15.9 | 0.0337 | 491 | 672 | 596 | 7.20 |
| 5 | 697 | 0.0023 | 2.9 | 0.0050 | 697 | 874 | 741 | 5.27 |
| 6 | 6000 (UM) | 0.0000 | | 0.0000 | 1508 | 6596 | 5009 | 20.21 |
| | TIC: | 0.0768 | ng/uL | | | | | |
| | TIM: | 1.0453 | nmole/L | | | | | |
| | Total Conc.: | 0.0953 | ng/uL | | | | | |

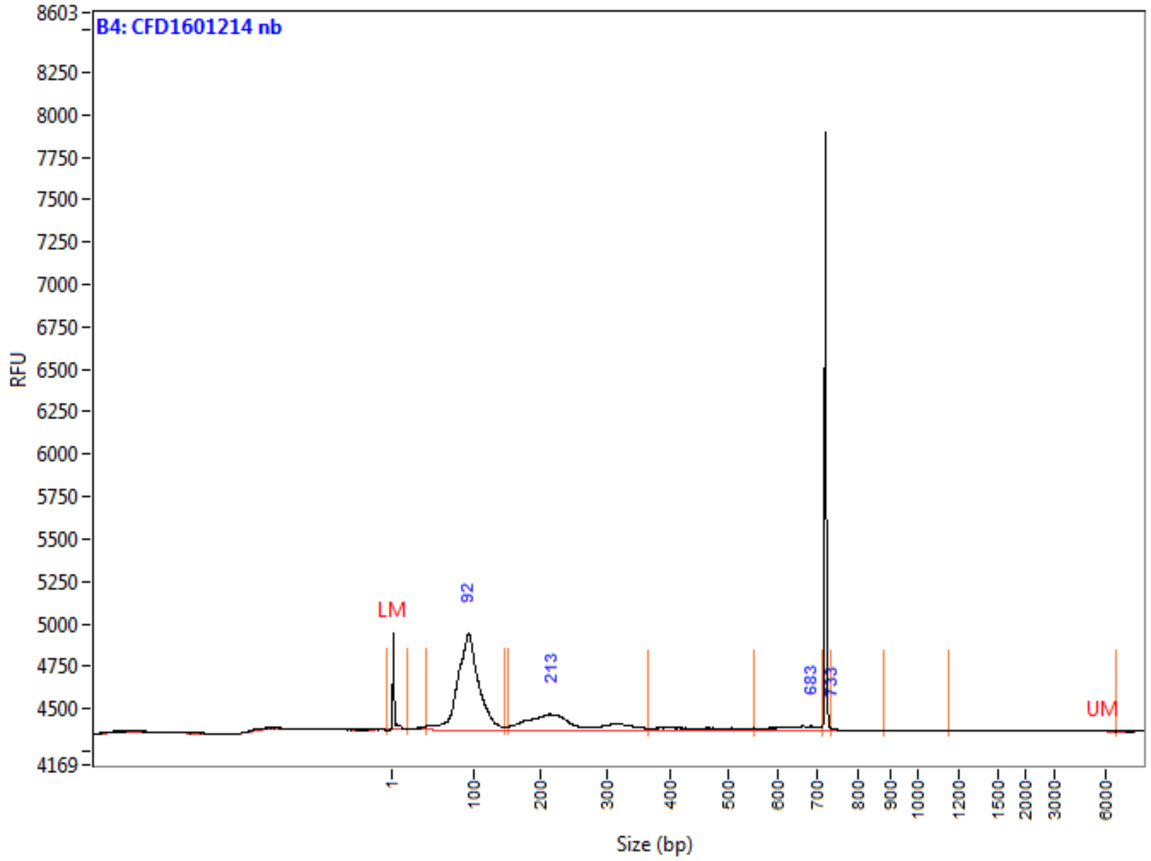

**Peak Table**

| | Size (bp) | ng/uL | % (Conc.) (ng/uL) | nmole/L | From (bp) | To (bp) | Avg. Size | CV% |
|---|---|---|---|---|---|---|---|---|
| 1 | 1 (LM) | 0.0005 | | 0.3937 | 0 | 20 | 2 | 187.72 |
| 2 | 92 | 0.0565 | 72.4 | 1.0088 | 41 | 146 | 92 | 17.18 |
| 3 | 213 | 0.0188 | 24.1 | 0.1337 | 150 | 366 | 231 | 22.91 |
| 4 | 683 | 0.0026 | 3.3 | 0.0067 | 551 | 712 | 636 | 7.15 |
| 5 | 733 | 0.0002 | 0.2 | 0.0004 | 733 | 884 | 745 | 1.79 |
| 6 | 6000 (UM) | 0.0000 | | 0.0000 | 1157 | 6665 | 4668 | 34.87 |
| | TIC: | 0.0781 | ng/uL | | | | | |
| | TIM: | 1.1495 | nmole/L | | | | | |
| | Total Conc.: | 0.0966 | ng/uL | | | | | |

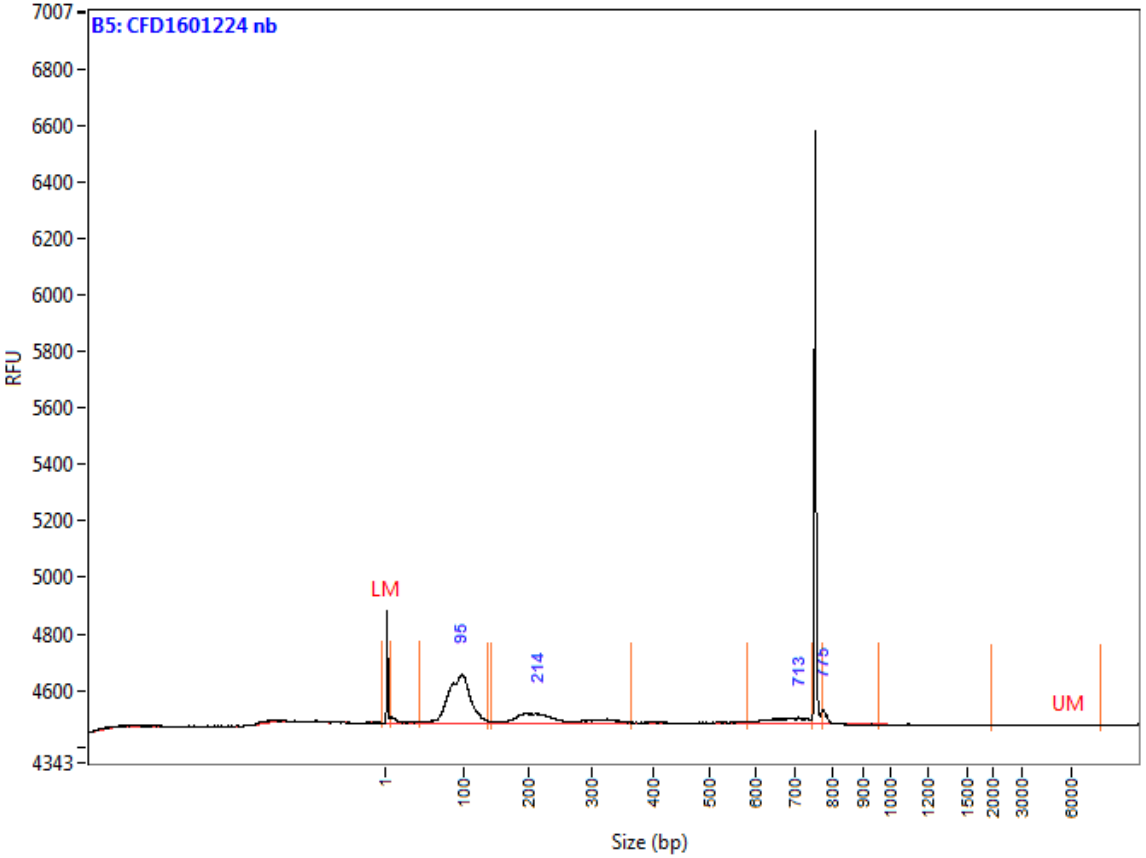

**Peak Table**

| | Size (bp) | ng/uL | % (Conc.) (ng/uL) | nmole/L | From (bp) | To (bp) | Avg. Size | CV% |
|---|---|---|---|---|---|---|---|---|
| 1 | 1 (LM) | 0.0005 | | 0.7062 | 0 | 8 | 1 | 173.23 |
| 2 | 95 | 0.0293 | 67.5 | 0.5232 | 44 | 137 | 92 | 15.45 |
| 3 | 214 | 0.0099 | 22.7 | 0.0700 | 142 | 363 | 232 | 22.22 |
| 4 | 713 | 0.0034 | 7.8 | 0.0082 | 581 | 748 | 673 | 6.60 |
| 5 | 775 | 0.0009 | 2.0 | 0.0019 | 775 | 958 | 784 | 1.05 |
| 6 | 6000 (UM) | 0.0000 | | 0.0000 | 1993 | 7925 | 7163 | 9.39 |
| | TIC: | 0.0434 | ng/uL | | | | | |
| | TIM: | 0.6033 | nmole/L | | | | | |
| | Total Conc.: | 0.0605 | ng/uL | | | | | |

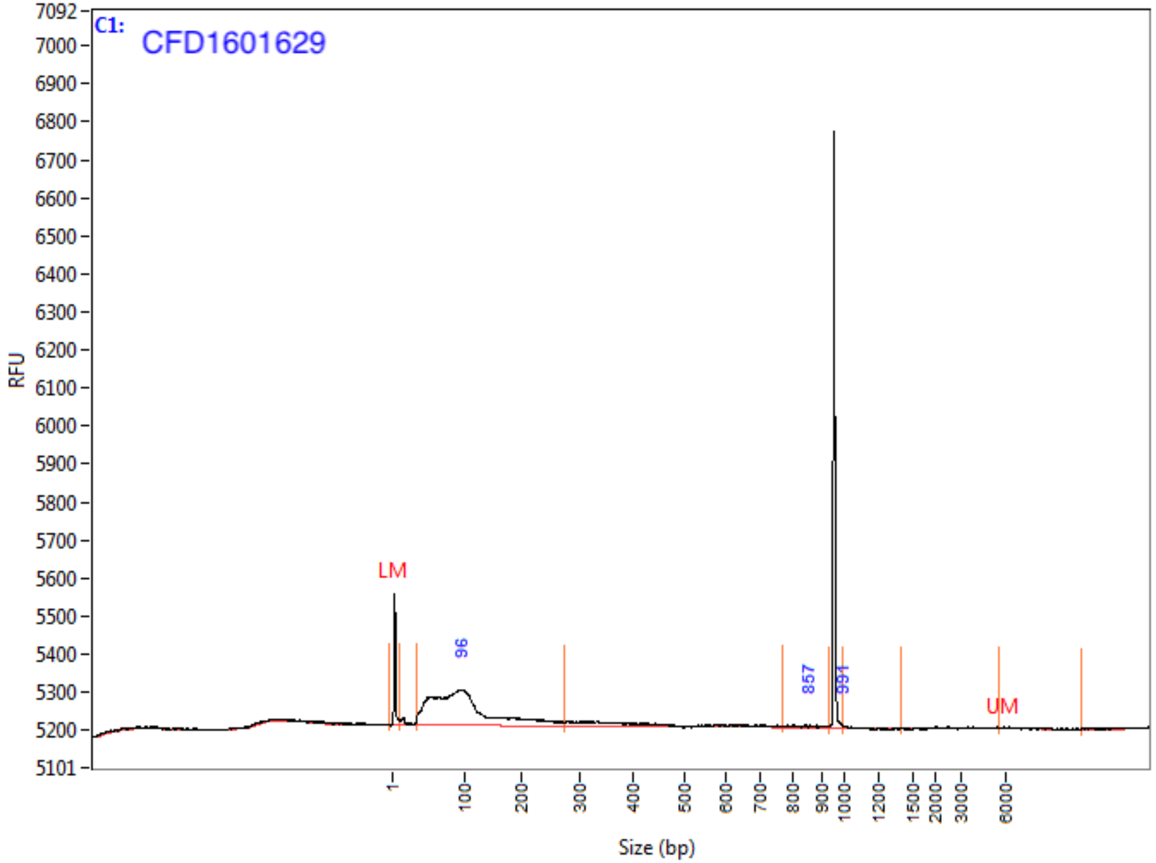

**Peak Table**

| | Size (bp) | ng/uL | % (Conc.) (ng/uL) | nmole/L | From (bp) | To (bp) | Avg. Size | CV% |
|---|---|---|---|---|---|---|---|---|
| 1 | 1 (LM) | 0.0004 | | 0.5401 | 0 | 9 | 1 | 169.22 |
| 2 | 96 | 0.0314 | 98.9 | 0.5111 | 34 | 272 | 101 | 52.29 |
| 3 | 857 | 0.0003 | 1.0 | 0.0006 | 768 | 932 | 840 | 5.95 |
| 4 | 991 | 0.0000 | 0.1 | 0.0001 | 991 | 1402 | 1005 | 1.65 |
| 5 | 6000 (UM) | 0.0000 | | 0.0000 | 5605 | 11288 | 6766 | 16.16 |
| | TIC: | 0.0318 | ng/uL | | | | | |
| | TIM: | 0.5118 | nmole/L | | | | | |
| | Total Conc.: | 0.0456 | ng/uL | | | | | |

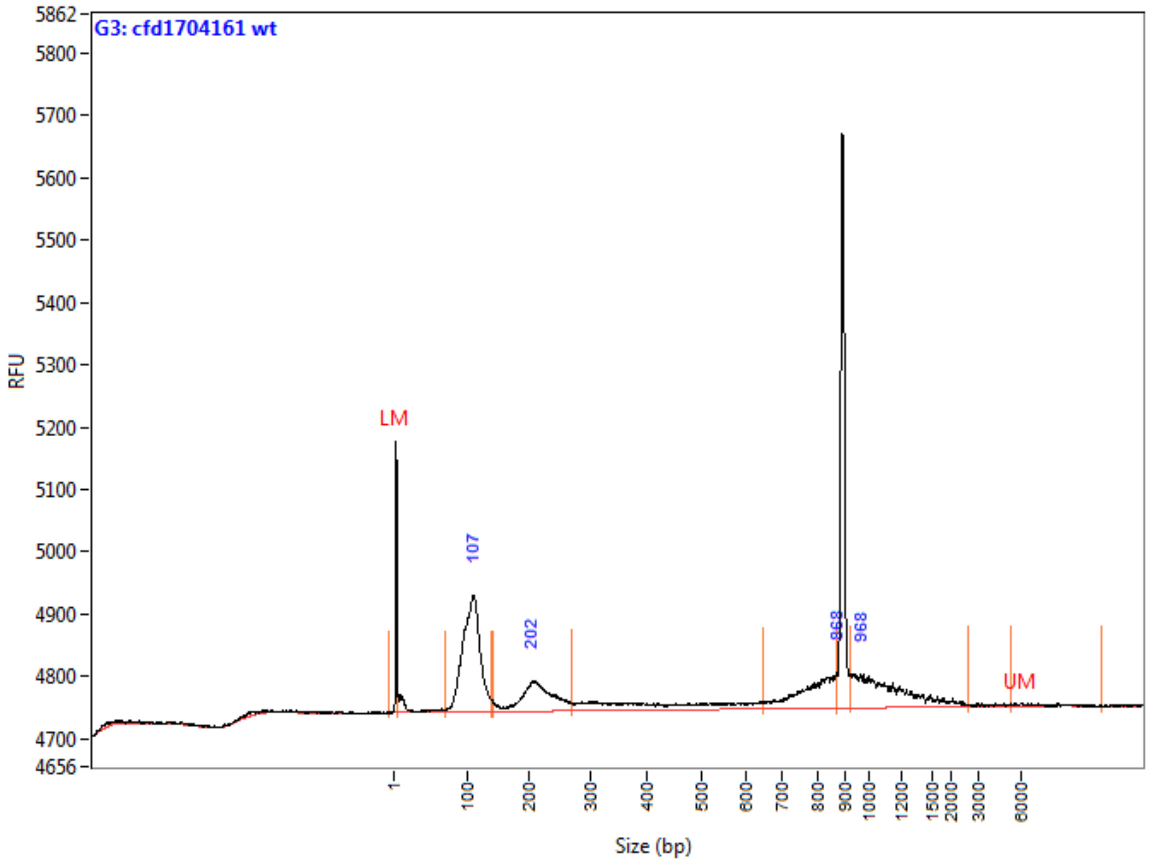

**Peak Table**

| | Size (bp) | ng/uL | % (Conc.) (ng/uL) | nmole/L | From (bp) | To (bp) | Avg. Size | CV% |
|---|---|---|---|---|---|---|---|---|
| 1 | 1 (LM) | 0.0007 | | 0.9609 | 0 | 5 | 1 | 198.47 |
| 2 | 107 | 0.0324 | 49.5 | 0.5164 | 69 | 136 | 103 | 12.12 |
| 3 | 202 | 0.0125 | 19.0 | 0.0971 | 142 | 271 | 211 | 13.24 |
| 4 | 868 | 0.0090 | 13.7 | 0.0188 | 647 | 869 | 783 | 7.40 |
| 5 | 968 | 0.0117 | 17.8 | 0.0164 | 920 | 2647 | 1169 | 24.77 |
| 6 | 6000 (UM) | 0.0000 | | 0.0000 | 5313 | 11641 | 7414 | 26.00 |
| | TIC: | 0.0655 | ng/uL | | | | | |
| | TIM: | 0.6487 | nmole/L | | | | | |
| | Total Conc.: | 0.0960 | ng/uL | | | | | |

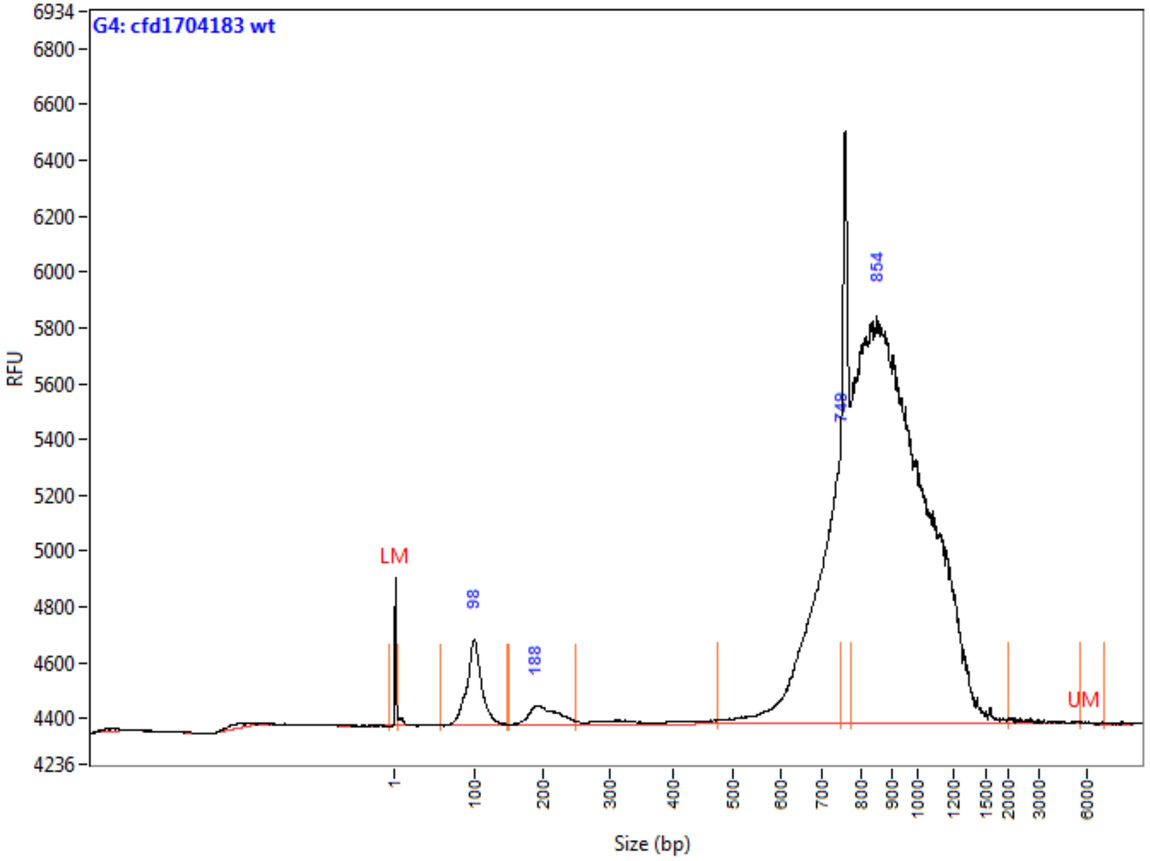

**Peak Table**

| | Size (bp) | ng/uL | % (Conc.) (ng/uL) | nmole/L | From (bp) | To (bp) | Avg. Size | CV% |
|---|---|---|---|---|---|---|---|---|
| 1 | 1 (LM) | 0.0007 | | 0.9609 | 0 | 5 | 1 | 145.75 |
| 2 | 98 | 0.0374 | 6.8 | 0.6274 | 59 | 148 | 98 | 11.27 |
| 3 | 188 | 0.0133 | 2.4 | 0.1083 | 149 | 248 | 202 | 9.79 |
| 4 | 749 | 0.1076 | 19.4 | 0.2577 | 474 | 750 | 687 | 7.82 |
| 5 | 854 | 0.3953 | 71.4 | 0.6914 | 776 | 2036 | 941 | 14.86 |
| 6 | 6000 (UM) | 0.0000 | | 0.0000 | 5725 | 7160 | 6375 | 7.19 |
| | TIC: | 0.5536 | ng/uL | | | | | |
| | TIM: | 1.6848 | nmole/L | | | | | |
| | Total Conc.: | 0.6173 | ng/uL | | | | | |

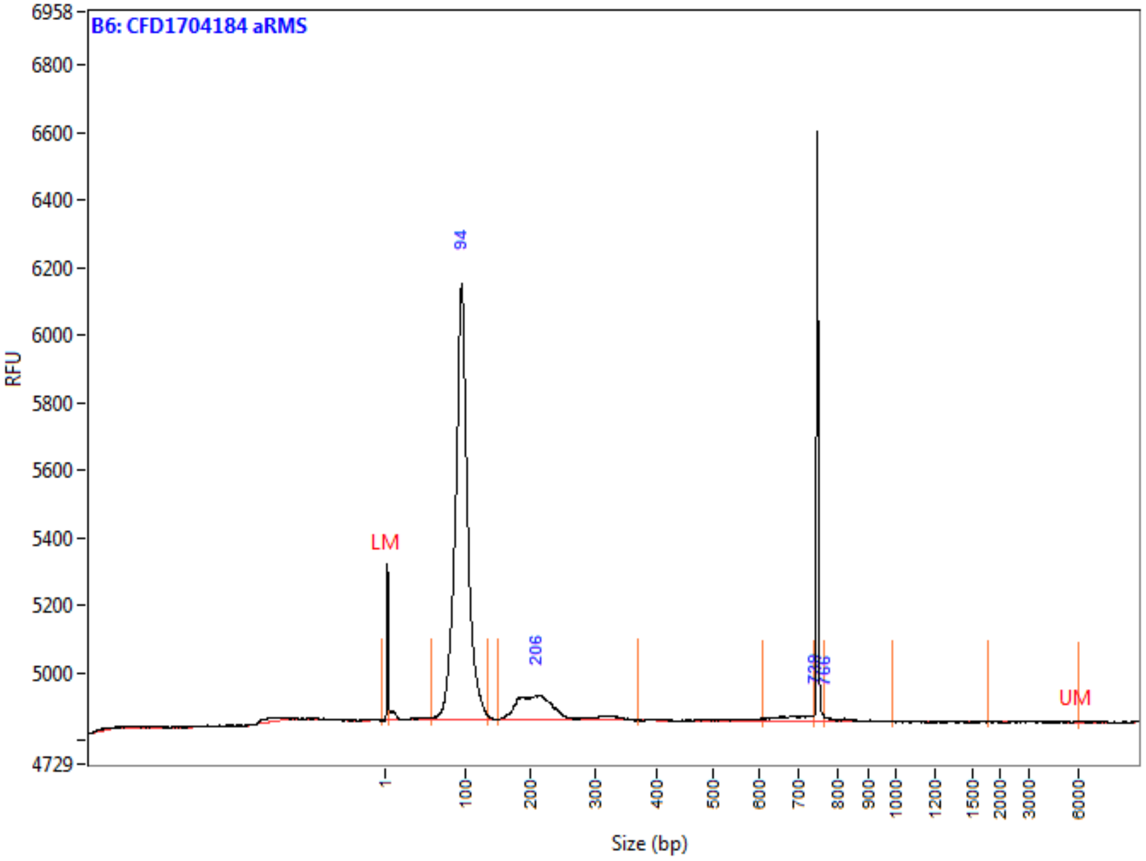

**Peak Table**

| | Size (bp) | ng/uL | % (Conc.) (ng/uL) | nmole/L | From (bp) | To (bp) | Avg. Size | CV% |
|---|---|---|---|---|---|---|---|---|
| 1 | 1 (LM) | 0.0005 | | 0.7062 | 0 | 5 | 1 | 176.80 |
| 2 | 94 | 0.1121 | 85.7 | 1.9582 | 58 | 134 | 94 | 10.40 |
| 3 | 206 | 0.0167 | 12.8 | 0.1278 | 149 | 372 | 215 | 18.83 |
| 4 | 739 | 0.0017 | 1.3 | 0.0040 | 607 | 740 | 681 | 5.31 |
| 5 | 766 | 0.0004 | 0.3 | 0.0008 | 766 | 990 | 801 | 4.07 |
| 6 | 6000 (UM) | 0.0000 | | 0.0000 | 1795 | 6092 | 5784 | 8.18 |
| | TIC: | 0.1309 | ng/uL | | | | | |
| | TIM: | 2.0908 | nmole/L | | | | | |
| | Total Conc.: | 0.1444 | ng/uL | | | | | |

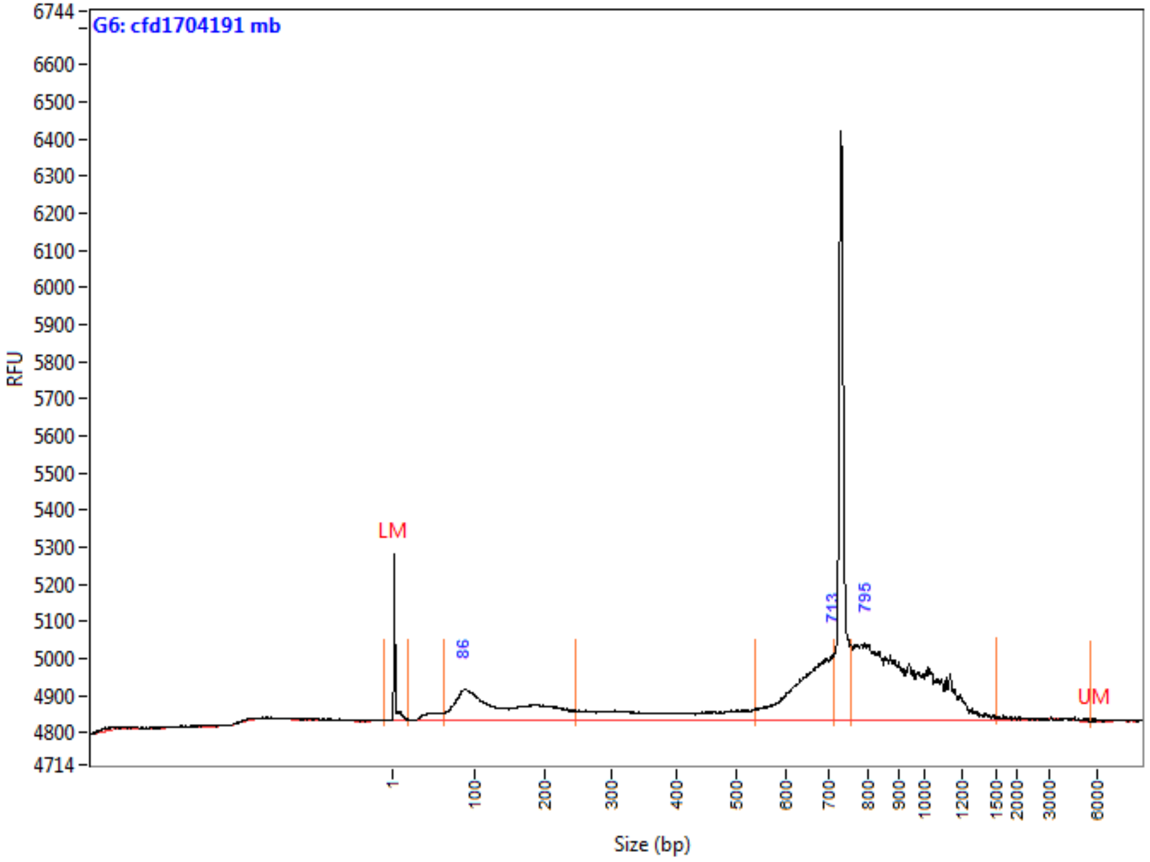

**Peak Table**

| | Size (bp) | ng/uL | % (Conc.) (ng/uL) | nmole/L | From (bp) | To (bp) | Avg. Size | CV% |
|---|---|---|---|---|---|---|---|---|
| 1 | 1 (LM) | 0.0007 | | 0.5357 | 0 | 20 | 2 | 160.33 |
| 2 | 86 | 0.0354 | 30.4 | 0.4272 | 63 | 245 | 136 | 37.87 |
| 3 | 713 | 0.0272 | 23.4 | 0.0692 | 536 | 715 | 647 | 7.25 |
| 4 | 795 | 0.0537 | 46.2 | 0.0959 | 759 | 1527 | 921 | 14.88 |
| 5 | 6000 (UM) | 0.0000 | | 0.0000 | 5702 | 8866 | 6777 | 14.10 |
| | TIC: | 0.1162 | ng/uL | | | | | |
| | TIM: | 0.5923 | nmole/L | | | | | |
| | Total Conc.: | 0.1710 | ng/uL | | | | | |

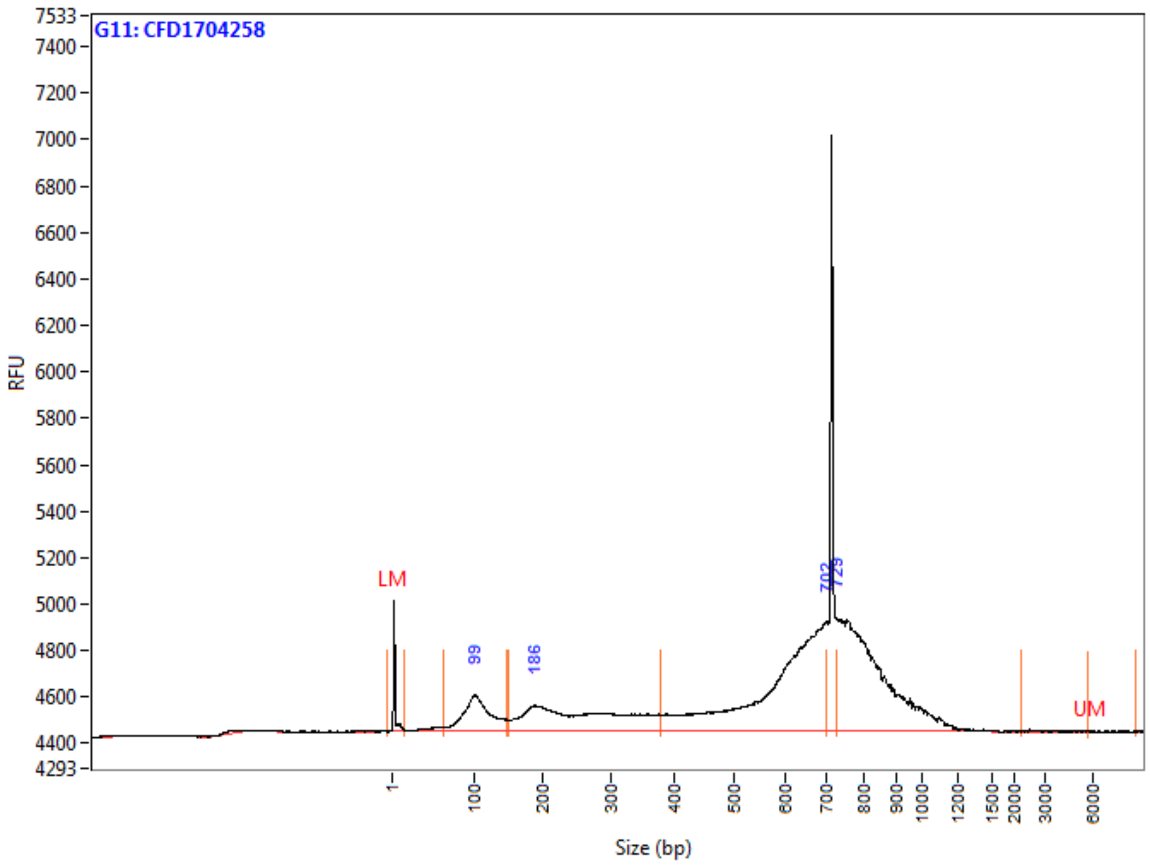

**Peak Table**

| | Size (bp) | ng/uL | % (Conc.) (ng/uL) | nmole/L | From (bp) | To (bp) | Avg. Size | CV% |
|---|---|---|---|---|---|---|---|---|
| 1 | 1 (LM) | 0.0007 | | 0.5357 | 0 | 16 | 2 | 157.74 |
| 2 | 99 | 0.0270 | 11.2 | 0.4229 | 63 | 150 | 105 | 18.98 |
| 3 | 186 | 0.0506 | 21.0 | 0.3275 | 148 | 377 | 254 | 25.57 |
| 4 | 702 | 0.0917 | 38.1 | 0.2584 | 377 | 703 | 584 | 15.63 |
| 5 | 729 | 0.0715 | 29.7 | 0.1413 | 729 | 2271 | 833 | 12.32 |
| 6 | 6000 (UM) | 0.0000 | | 0.0000 | 5771 | 8729 | 6862 | 12.63 |
| | TIC: | 0.2409 | ng/uL | | | | | |
| | TIM: | 1.1502 | nmole/L | | | | | |
| | Total Conc.: | 0.2687 | ng/uL | | | | | |

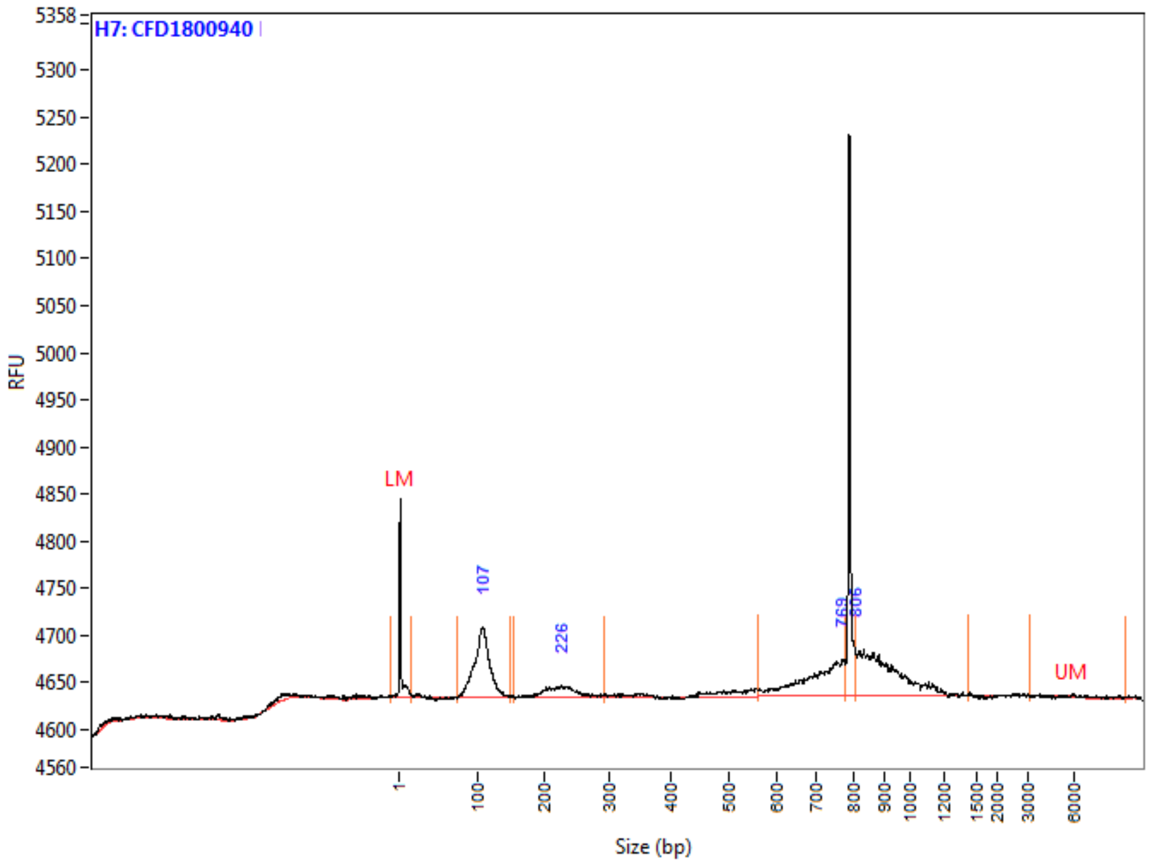

**Peak Table**

| | Size (bp) | ng/uL | % (Conc.) (ng/uL) | nmole/L | From (bp) | To (bp) | Avg. Size | CV% |
|---|---|---|---|---|---|---|---|---|
| 1 | 1 (LM) | 0.0007 | | 0.5357 | 0 | 18 | 2 | 184.61 |
| 2 | 107 | 0.0173 | 37.4 | 0.2713 | 75 | 150 | 105 | 10.91 |
| 3 | 226 | 0.0039 | 8.4 | 0.0288 | 155 | 292 | 223 | 9.54 |
| 4 | 769 | 0.0108 | 23.3 | 0.0254 | 561 | 778 | 700 | 8.16 |
| 5 | 806 | 0.0143 | 30.8 | 0.0258 | 806 | 1425 | 912 | 10.09 |
| 6 | 6000 (UM) | 0.0000 | | 0.0000 | 3184 | 9389 | 7472 | 14.83 |
| | TIC: | 0.0463 | ng/uL | | | | | |
| | TIM: | 0.3513 | nmole/L | | | | | |
| | Total Conc.: | 0.0604 | ng/uL | | | | | |

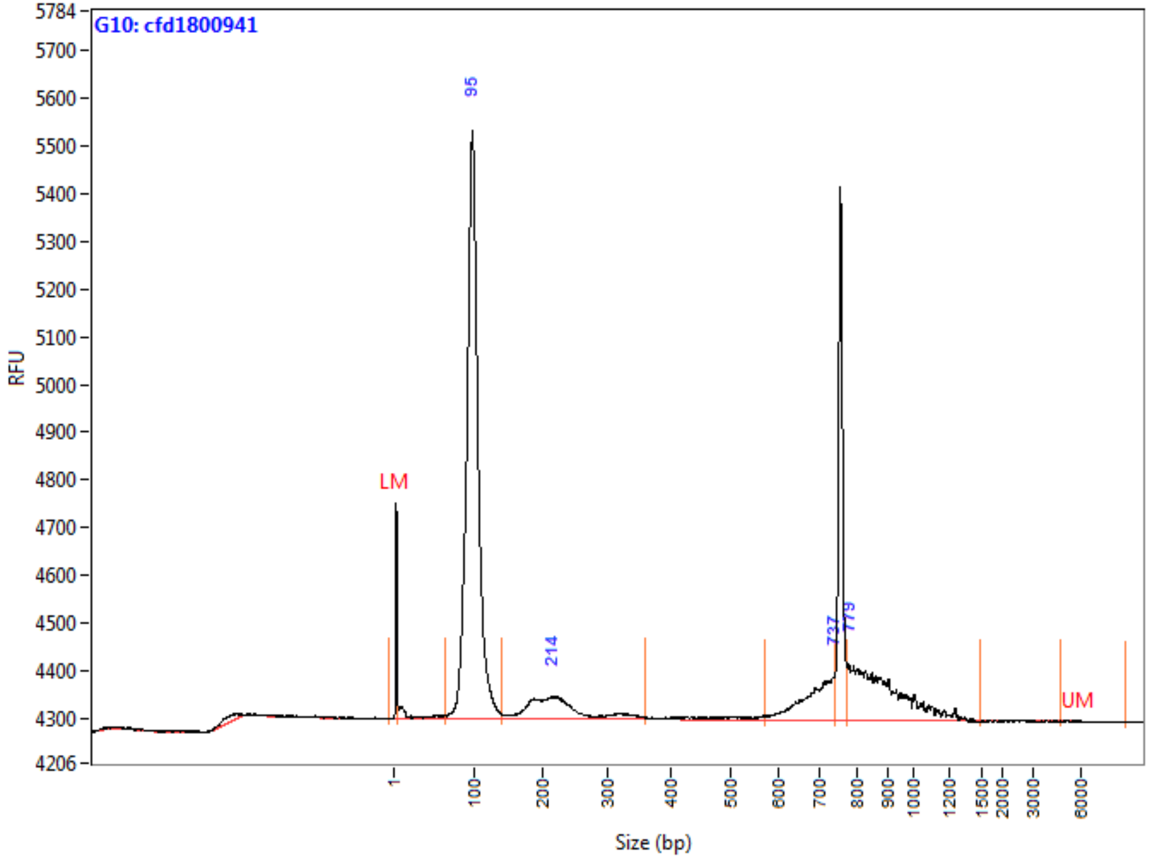

**Peak Table**

| | Size (bp) | ng/uL | % (Conc.) (ng/uL) | nmole/L | From (bp) | To (bp) | Avg. Size | CV% |
|---|---|---|---|---|---|---|---|---|
| 1 | 1 (LM) | 0.0007 | | 0.9609 | 0 | 5 | 1 | 150.17 |
| 2 | 95 | 0.1383 | 71.2 | 2.3658 | 63 | 139 | 96 | 9.51 |
| 3 | 214 | 0.0179 | 9.2 | 0.1337 | 140 | 357 | 220 | 20.65 |
| 4 | 737 | 0.0129 | 6.6 | 0.0309 | 571 | 742 | 684 | 6.16 |
| 5 | 779 | 0.0251 | 12.9 | 0.0461 | 775 | 1490 | 897 | 12.21 |
| 6 | 6000 (UM) | 0.0000 | | 0.0000 | 4787 | 8911 | 6982 | 17.40 |
| | TIC: | 0.1942 | ng/uL | | | | | |
| | TIM: | 2.5766 | nmole/L | | | | | |
| | Total Conc.: | 0.2227 | ng/uL | | | | | |

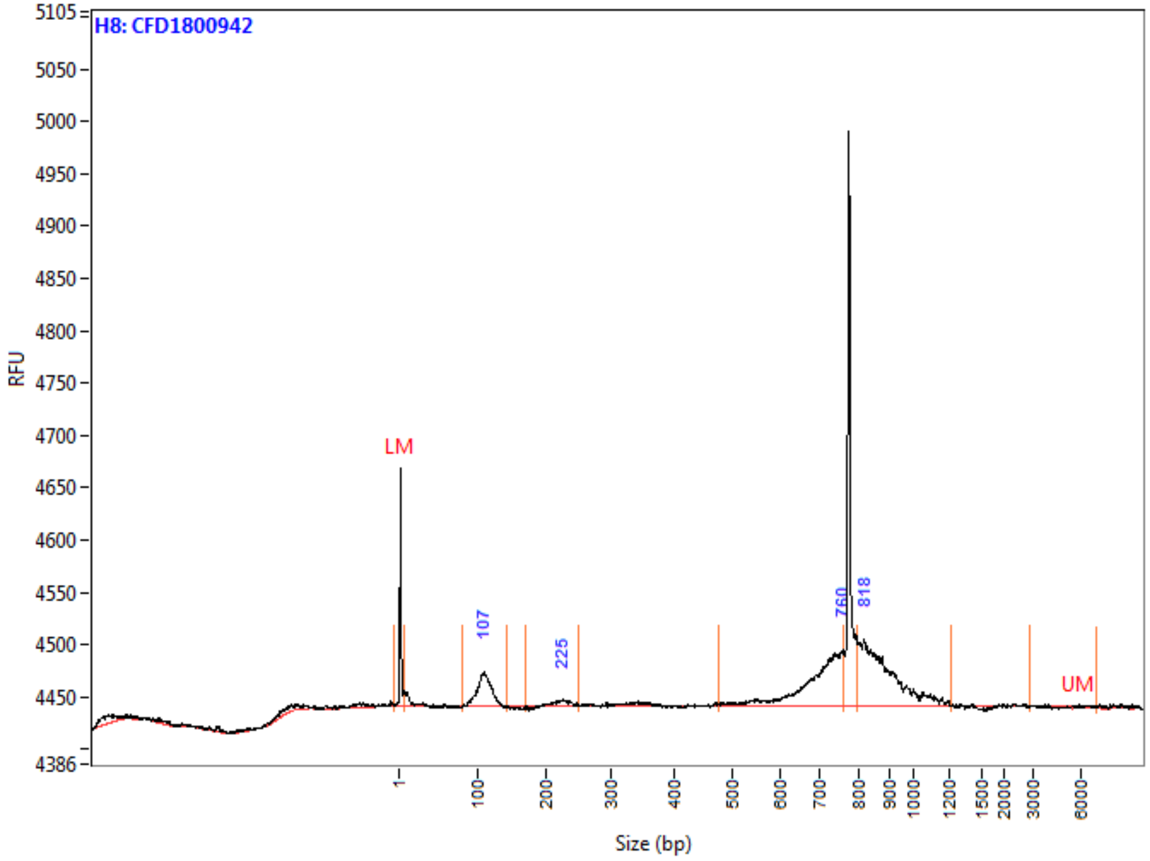

**Peak Table**

| | Size (bp) | ng/uL | % (Conc.) (ng/uL) | nmole/L | From (bp) | To (bp) | Avg. Size | CV% |
|---|---|---|---|---|---|---|---|---|
| 1 | 1 (LM) | 0.0007 | | 0.9609 | 0 | 7 | 1 | 199.41 |
| 2 | 107 | 0.0070 | 17.8 | 0.1079 | 80 | 143 | 106 | 8.66 |
| 3 | 225 | 0.0009 | 2.4 | 0.0069 | 171 | 252 | 222 | 5.18 |
| 4 | 760 | 0.0146 | 37.3 | 0.0350 | 476 | 761 | 687 | 9.09 |
| 5 | 818 | 0.0166 | 42.5 | 0.0309 | 797 | 1217 | 887 | 9.11 |
| 6 | 6000 (UM) | 0.0000 | | 0.0000 | 2894 | 7069 | 6233 | 10.11 |
| | TIC: | 0.0391 | ng/uL | | | | | |
| | TIM: | 0.1807 | nmole/L | | | | | |
| | Total Conc.: | 0.0559 | ng/uL | | | | | |

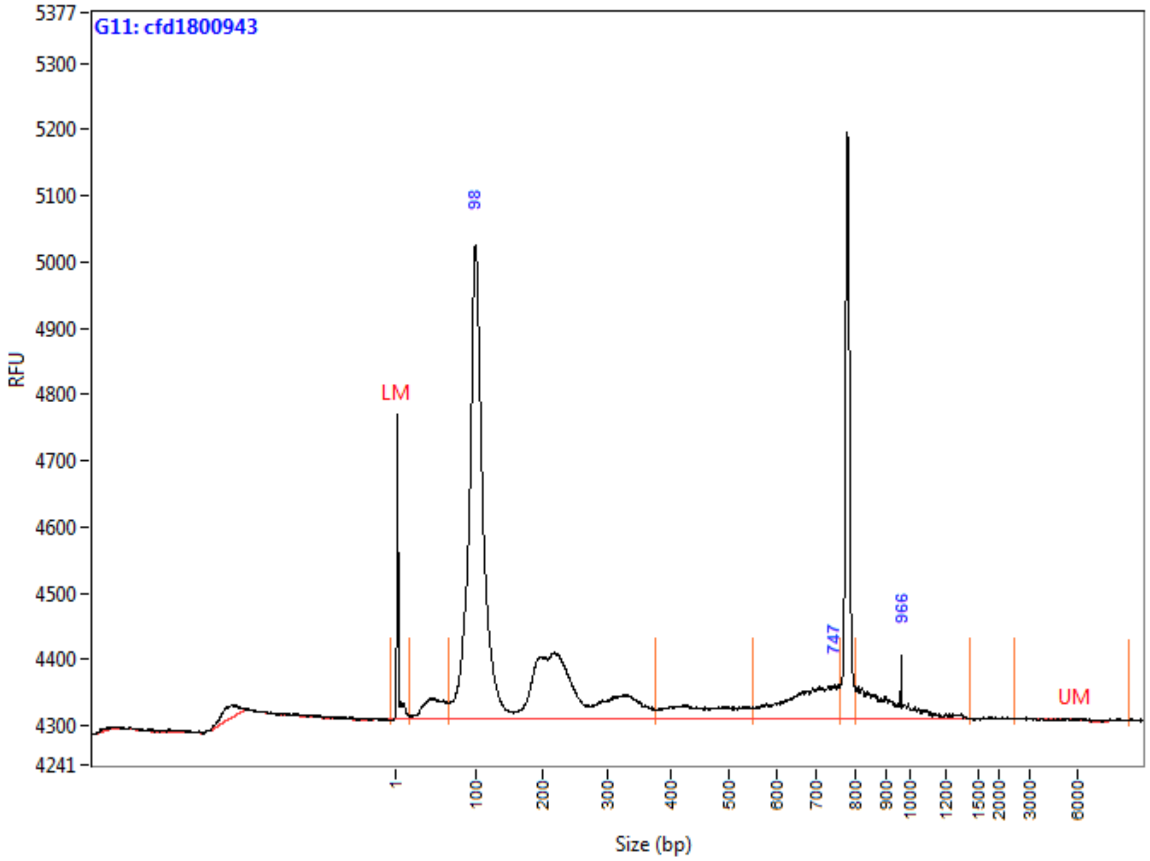

**Peak Table**

| | Size (bp) | ng/uL | % (Conc.) (ng/uL) | nmole/L | From (bp) | To (bp) | Avg. Size | CV% |
|---|---|---|---|---|---|---|---|---|
| 1 | 1 (LM) | 0.0007 | | 0.5357 | 0 | 18 | 2 | 177.69 |
| 2 | 98 | 0.1135 | 86.6 | 1.3044 | 67 | 375 | 143 | 50.16 |
| 3 | 747 | 0.0107 | 8.2 | 0.0263 | 550 | 762 | 670 | 8.77 |
| 4 | 966 | 0.0069 | 5.2 | 0.0122 | 804 | 1428 | 926 | 12.23 |
| 5 | 6000 (UM) | 0.0000 | | 0.0000 | 2518 | 9435 | 7040 | 18.10 |
| | TIC: | 0.1311 | ng/uL | | | | | |
| | TIM: | 1.3429 | nmole/L | | | | | |
| | Total Conc.: | 0.1599 | ng/uL | | | | | |

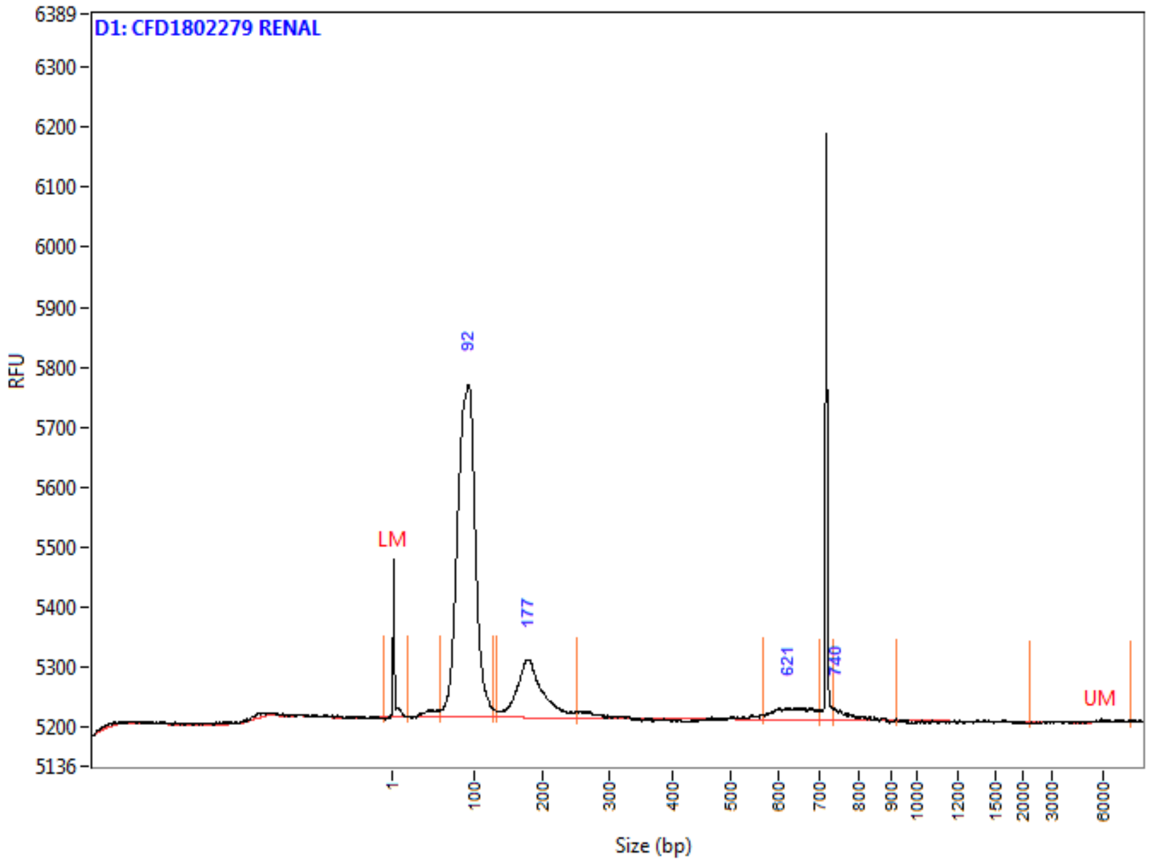

**Peak Table**

| | Size (bp) | ng/uL | % (Conc.) (ng/uL) | nmole/L | From (bp) | To (bp) | Avg. Size | CV% |
|---|---|---|---|---|---|---|---|---|
| 1 | 1 (LM) | 0.0005 | | 0.3740 | 0 | 20 | 2 | 155.44 |
| 2 | 92 | 0.1042 | 79.4 | 1.9432 | 58 | 126 | 88 | 11.54 |
| 3 | 177 | 0.0216 | 16.5 | 0.1954 | 133 | 250 | 182 | 12.19 |
| 4 | 621 | 0.0040 | 3.1 | 0.0105 | 567 | 701 | 636 | 5.75 |
| 5 | 740 | 0.0014 | 1.1 | 0.0030 | 736 | 921 | 775 | 4.58 |
| 6 | 6000 (UM) | 0.0000 | | 0.0000 | 2299 | 7683 | 5678 | 16.73 |
| | TIC: | 0.1312 | ng/uL | | | | | |
| | TIM: | 2.1520 | nmole/L | | | | | |
| | Total Conc.: | 0.1445 | ng/uL | | | | | |

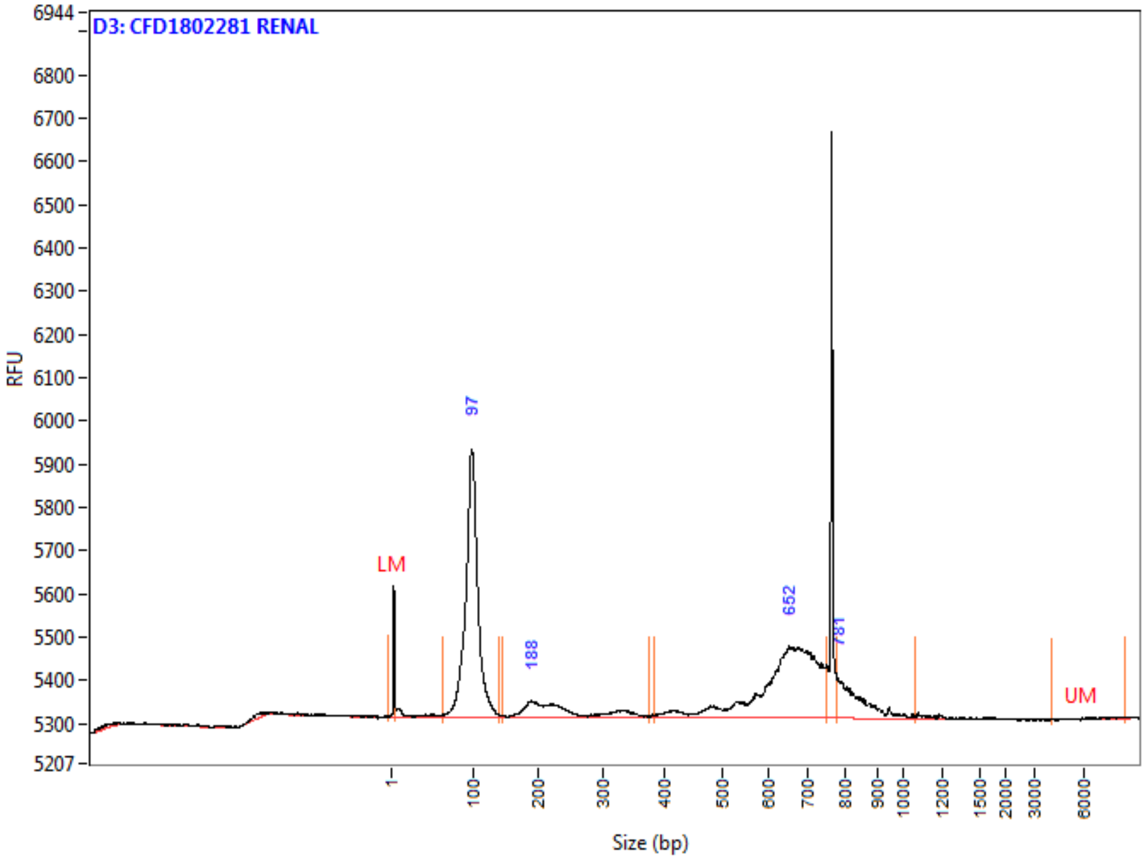

**Peak Table**

| | Size (bp) | ng/uL | % (Conc.) (ng/uL) | nmole/L | From (bp) | To (bp) | Avg. Size | CV% |
|---|---|---|---|---|---|---|---|---|
| 1 | 1 (LM) | 0.0005 | | 0.6708 | 0 | 4 | 1 | 153.77 |
| 2 | 97 | 0.0772 | 50.2 | 1.3067 | 62 | 138 | 97 | 10.18 |
| 3 | 188 | 0.0138 | 9.0 | 0.0965 | 146 | 376 | 235 | 23.57 |
| 4 | 652 | 0.0520 | 33.8 | 0.1350 | 384 | 750 | 634 | 12.99 |
| 5 | 781 | 0.0109 | 7.1 | 0.0213 | 776 | 1064 | 840 | 6.94 |
| 6 | 6000 (UM) | 0.0000 | | 0.0000 | 4146 | 8570 | 6877 | 13.91 |
| | TIC: | 0.1539 | ng/uL | | | | | |
| | TIM: | 1.5596 | nmole/L | | | | | |
| | Total Conc.: | 0.1721 | ng/uL | | | | | |

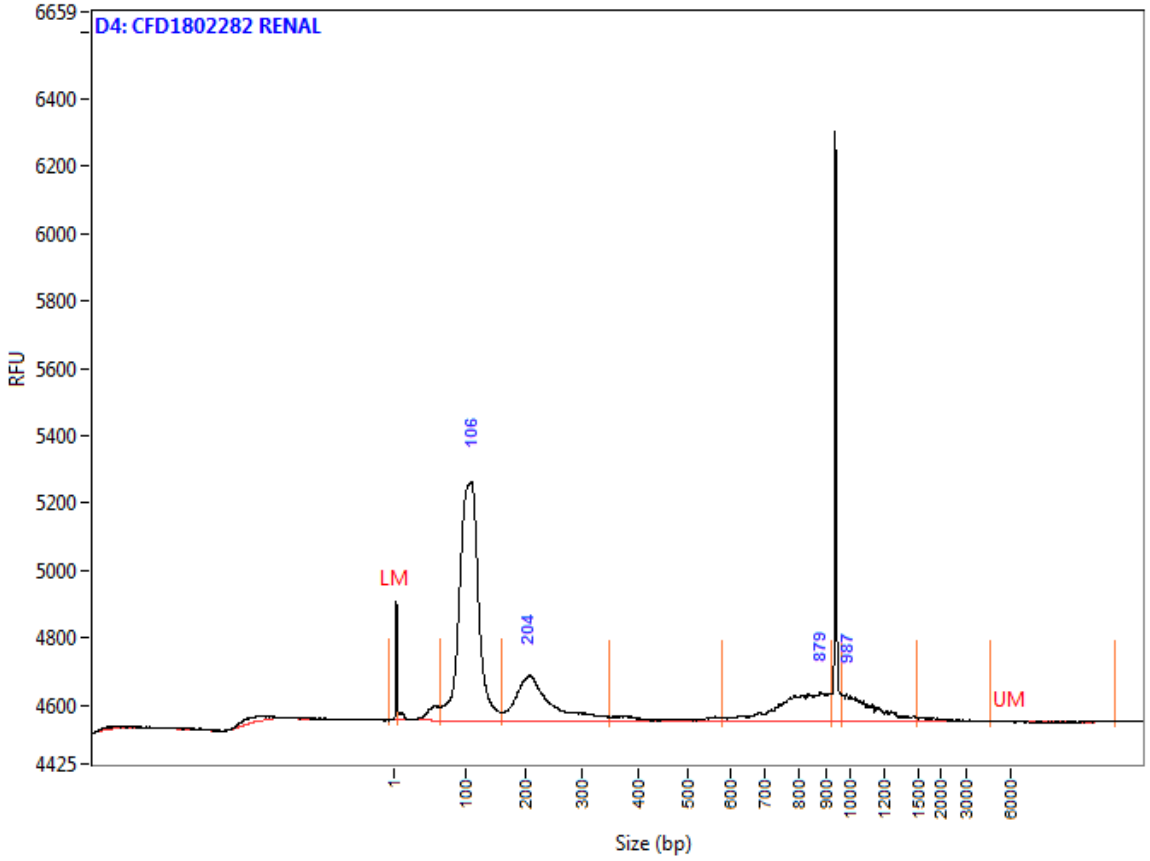

**Peak Table**

| | Size (bp) | ng/uL | % (Conc.) (ng/uL) | nmole/L | From (bp) | To (bp) | Avg. Size | CV% |
|---|---|---|---|---|---|---|---|---|
| 1 | 1 (LM) | 0.0005 | | 0.6708 | 0 | 5 | 1 | 197.16 |
| 2 | 106 | 0.1199 | 65.4 | 1.9110 | 64 | 158 | 103 | 14.36 |
| 3 | 204 | 0.0335 | 18.3 | 0.2472 | 160 | 350 | 223 | 18.16 |
| 4 | 879 | 0.0195 | 10.6 | 0.0403 | 581 | 920 | 797 | 10.03 |
| 5 | 987 | 0.0105 | 5.7 | 0.0156 | 965 | 1497 | 1107 | 11.37 |
| 6 | 6000 (UM) | 0.0000 | | 0.0000 | 4649 | 13164 | 9402 | 34.07 |
| | TIC: | 0.1834 | ng/uL | | | | | |
| | TIM: | 2.2141 | nmole/L | | | | | |
| | Total Conc.: | 0.2094 | ng/uL | | | | | |

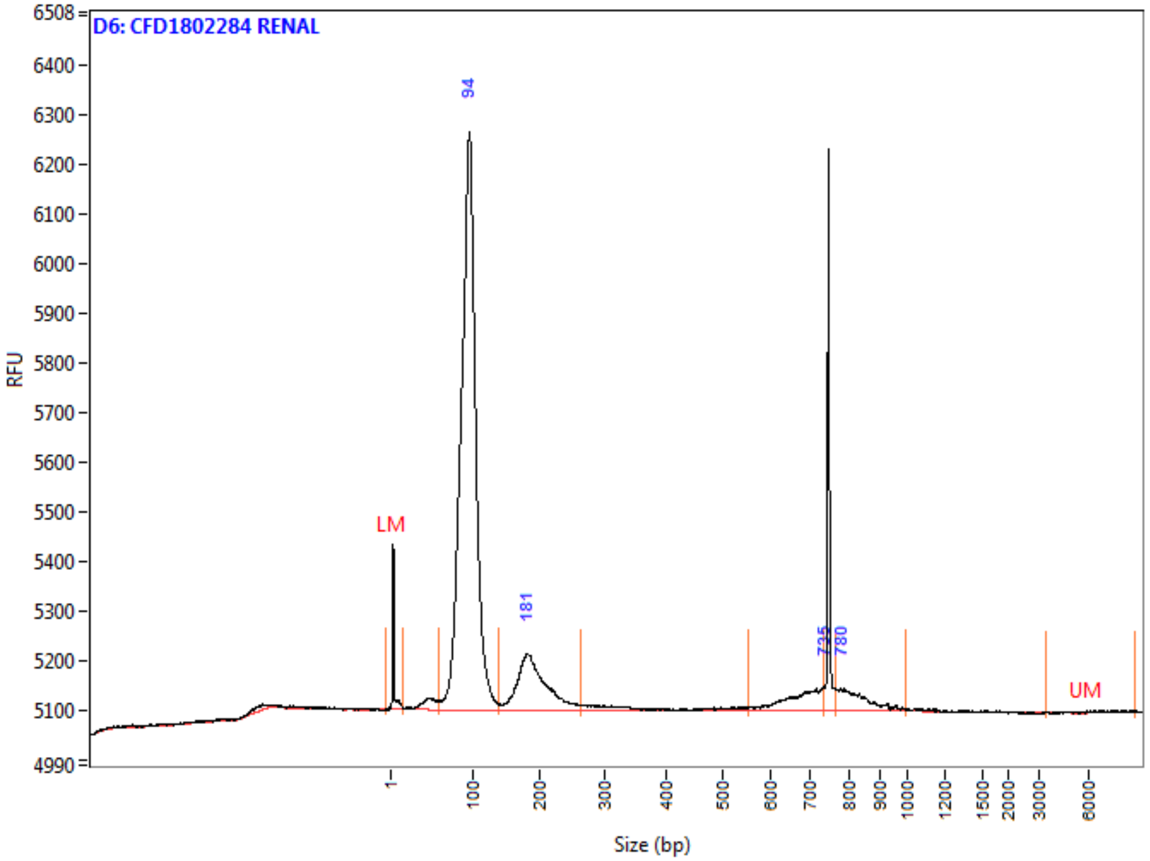

**Peak Table**

| | Size (bp) | ng/uL | % (Conc.) (ng/uL) | nmole/L | From (bp) | To (bp) | Avg. Size | CV% |
|---|---|---|---|---|---|---|---|---|
| 1 | 1 (LM) | 0.0005 | | 0.3740 | 0 | 14 | 2 | 166.91 |
| 2 | 94 | 0.1357 | 80.9 | 2.3960 | 59 | 139 | 93 | 11.29 |
| 3 | 181 | 0.0210 | 12.5 | 0.1808 | 140 | 265 | 191 | 12.82 |
| 4 | 735 | 0.0061 | 3.6 | 0.0150 | 553 | 736 | 670 | 6.94 |
| 5 | 780 | 0.0050 | 3.0 | 0.0099 | 766 | 999 | 829 | 6.24 |
| 6 | 6000 (UM) | 0.0000 | | 0.0000 | 3505 | 8888 | 7012 | 17.14 |
| | TIC: | 0.1678 | ng/uL | | | | | |
| | TIM: | 2.6017 | nmole/L | | | | | |
| | Total Conc.: | 0.1832 | ng/uL | | | | | |

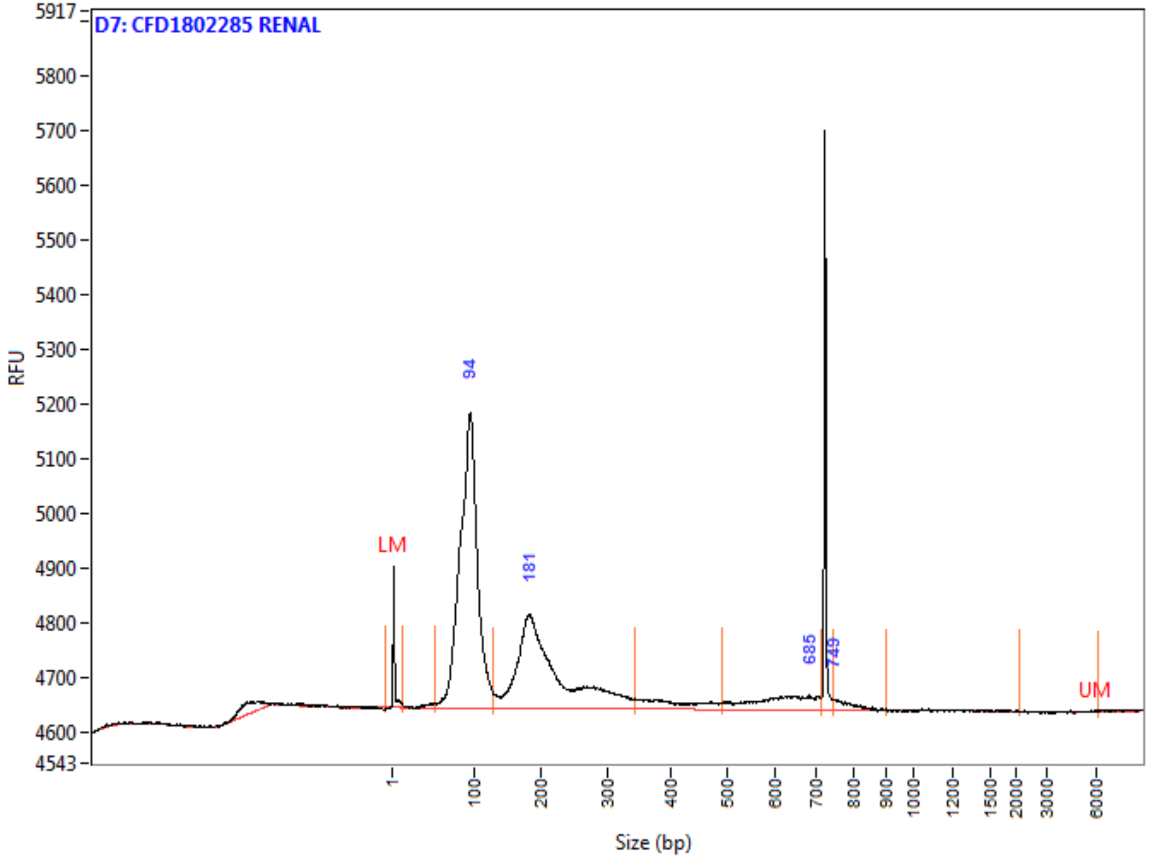

**Peak Table**

| | Size (bp) | ng/uL | % (Conc.) (ng/uL) | nmole/L | From (bp) | To (bp) | Avg. Size | CV% |
|---|---|---|---|---|---|---|---|---|
| 1 | 1 (LM) | 0.0005 | | 0.6708 | 0 | 14 | 1 | 182.66 |
| 2 | 94 | 0.0972 | 58.8 | 1.7537 | 53 | 128 | 91 | 12.92 |
| 3 | 181 | 0.0582 | 35.2 | 0.4579 | 129 | 344 | 209 | 23.21 |
| 4 | 685 | 0.0084 | 5.1 | 0.0226 | 489 | 715 | 611 | 10.25 |
| 5 | 749 | 0.0016 | 1.0 | 0.0034 | 746 | 900 | 783 | 3.96 |
| 6 | 6000 (UM) | 0.0000 | | 0.0000 | 2118 | 6136 | 5549 | 10.19 |
| | TIC: | 0.1655 | ng/uL | | | | | |
| | TIM: | 2.2377 | nmole/L | | | | | |
| | Total Conc.: | 0.1828 | ng/uL | | | | | |

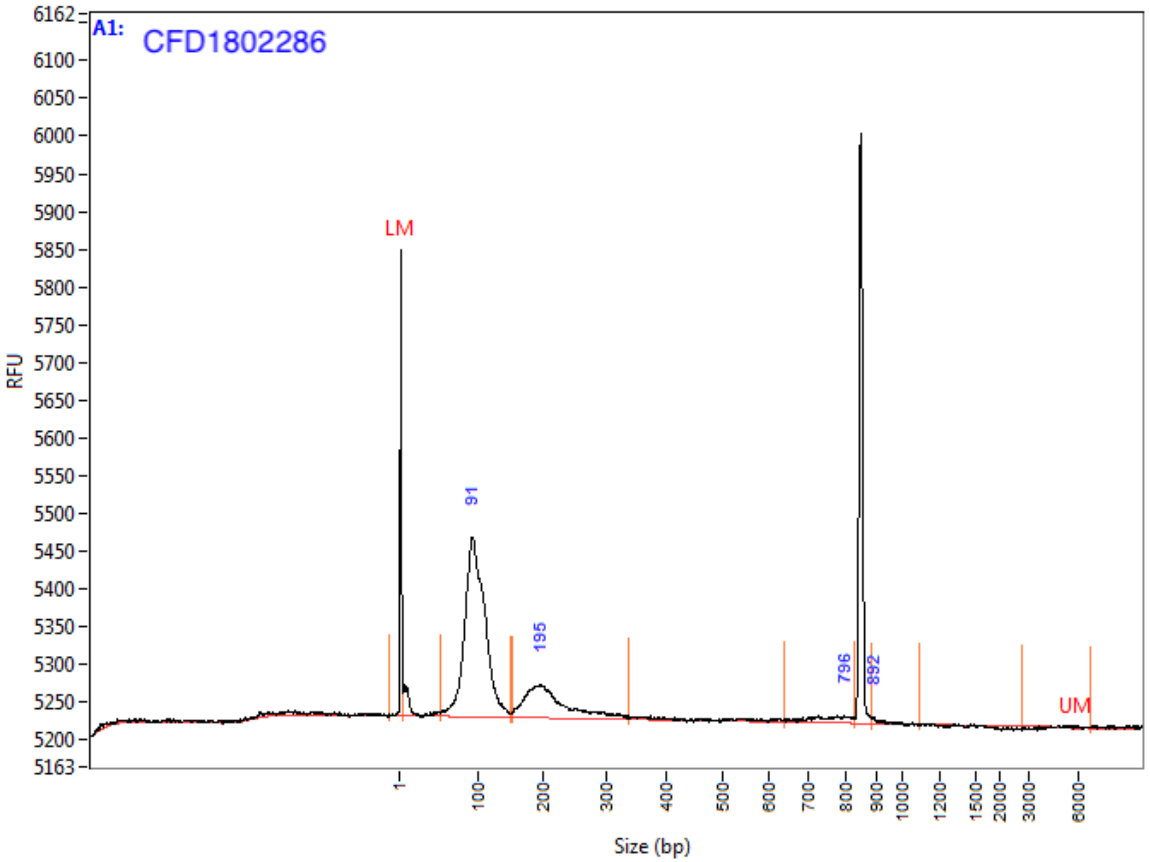

**Peak Table**

| | Size (bp) | ng/uL | % (Conc.) (ng/uL) | nmole/L | From (bp) | To (bp) | Avg. Size | CV% |
|---|---|---|---|---|---|---|---|---|
| 1 | 1 (LM) | 0.0011 | | 1.4680 | 0 | 5 | 1 | 148.28 |
| 2 | 91 | 0.0522 | 75.8 | 0.8930 | 54 | 151 | 96 | 15.20 |
| 3 | 195 | 0.0149 | 21.7 | 0.1162 | 154 | 336 | 211 | 18.55 |
| 4 | 796 | 0.0015 | 2.2 | 0.0033 | 639 | 829 | 754 | 6.14 |
| 5 | 892 | 0.0002 | 0.3 | 0.0004 | 886 | 1097 | 914 | 3.24 |
| 6 | 6000 (UM) | 0.0000 | | 0.0000 | 2788 | 6846 | 6072 | 5.68 |
| | TIC: | 0.0689 | ng/uL | | | | | |
| | TIM: | 1.0130 | nmole/L | | | | | |
| | Total Conc.: | 0.0879 | ng/uL | | | | | |

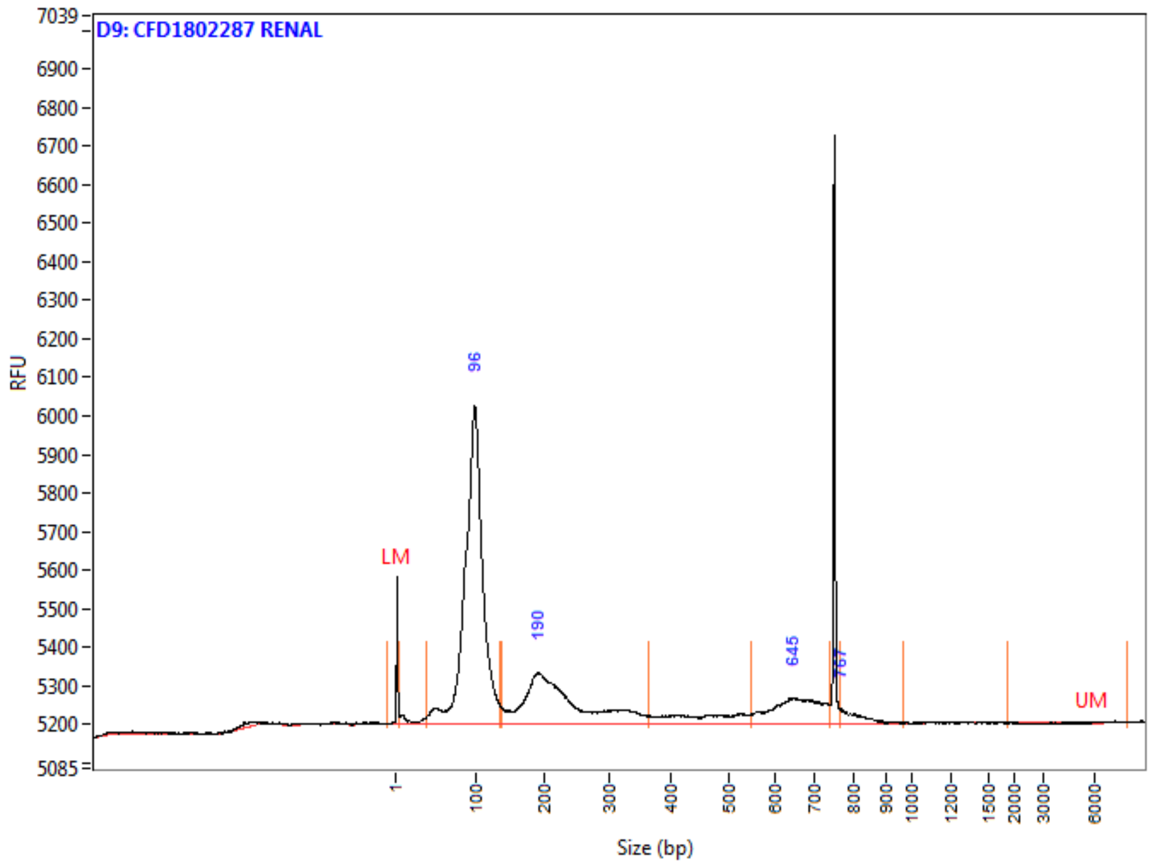

**Peak Table**

| | Size (bp) | ng/uL | % (Conc.) (ng/uL) | nmole/L | From (bp) | To (bp) | Avg. Size | CV% |
|---|---|---|---|---|---|---|---|---|
| 1 | 1 (LM) | 0.0005 | | 0.6708 | 0 | 5 | 1 | 186.93 |
| 2 | 96 | 0.1164 | 66.3 | 2.0332 | 39 | 136 | 94 | 16.13 |
| 3 | 190 | 0.0429 | 24.4 | 0.3164 | 136 | 366 | 223 | 24.34 |
| 4 | 645 | 0.0138 | 7.8 | 0.0349 | 548 | 739 | 650 | 7.79 |
| 5 | 767 | 0.0024 | 1.4 | 0.0049 | 765 | 969 | 807 | 4.13 |
| 6 | 6000 (UM) | 0.0000 | | 0.0000 | 1896 | 8070 | 6332 | 12.54 |
| | TIC: | 0.1755 | ng/uL | | | | | |
| | TIM: | 2.3893 | nmole/L | | | | | |
| | Total Conc.: | 0.1966 | ng/uL | | | | | |

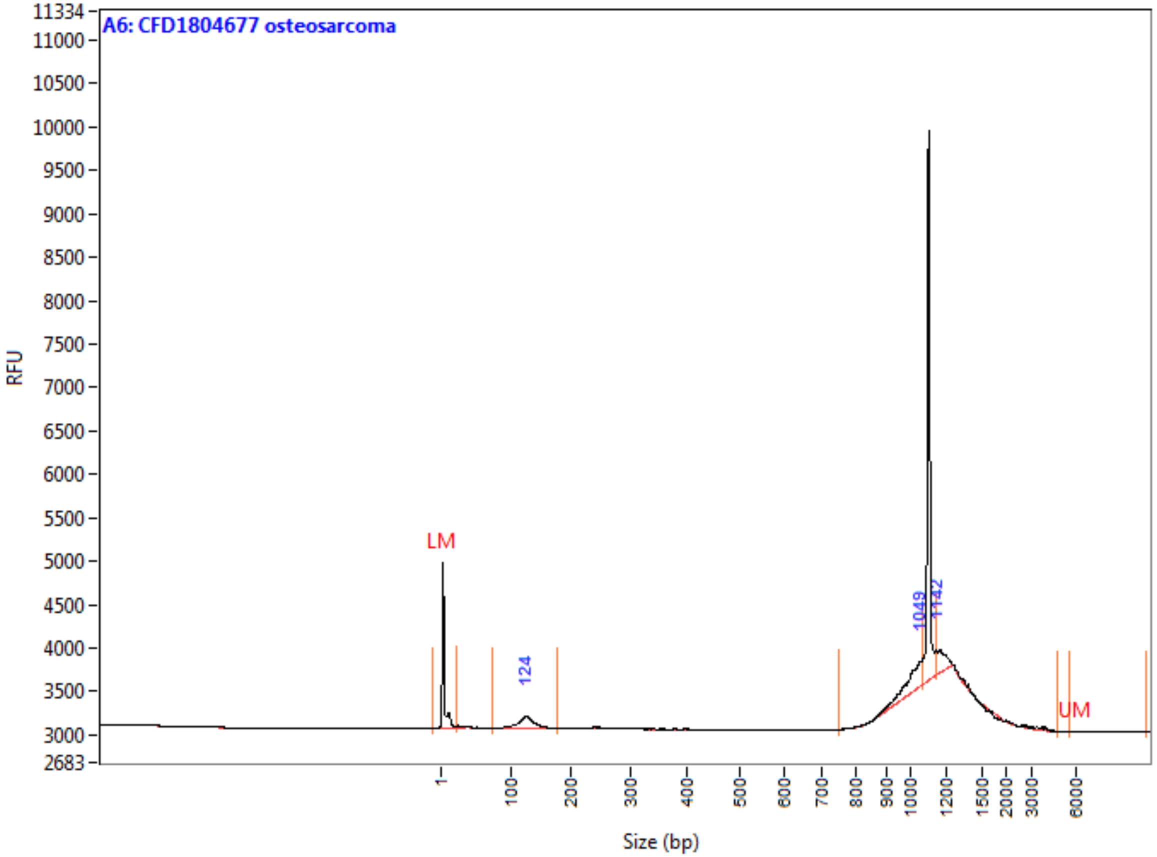

**Peak Table**

| | Size (bp) | ng/uL | % (Conc.) (ng/uL) | nmole/L | From (bp) | To (bp) | Avg. Size | CV% |
|---|---|---|---|---|---|---|---|---|
| 1 | 1 (LM) | 0.0004 | | 0.3273 | 0 | 23 | 2 | 191.97 |
| 2 | 124 | 0.0018 | 32.3 | 0.0235 | 74 | 176 | 123 | 11.58 |
| 3 | 1049 | 0.0022 | 41.1 | 0.0037 | 751 | 1066 | 996 | 5.17 |
| 4 | 1142 | 0.0014 | 26.6 | 0.0016 | 1142 | 4788 | 1526 | 52.07 |
| 5 | 6000 (UM) | 0.0000 | | 0.0000 | 5576 | 10718 | 7212 | 18.07 |
| | TIC: | 0.0054 | ng/uL | | | | | |
| | TIM: | 0.0288 | nmole/L | | | | | |
| | Total Conc.: | 0.0138 | ng/uL | | | | | |

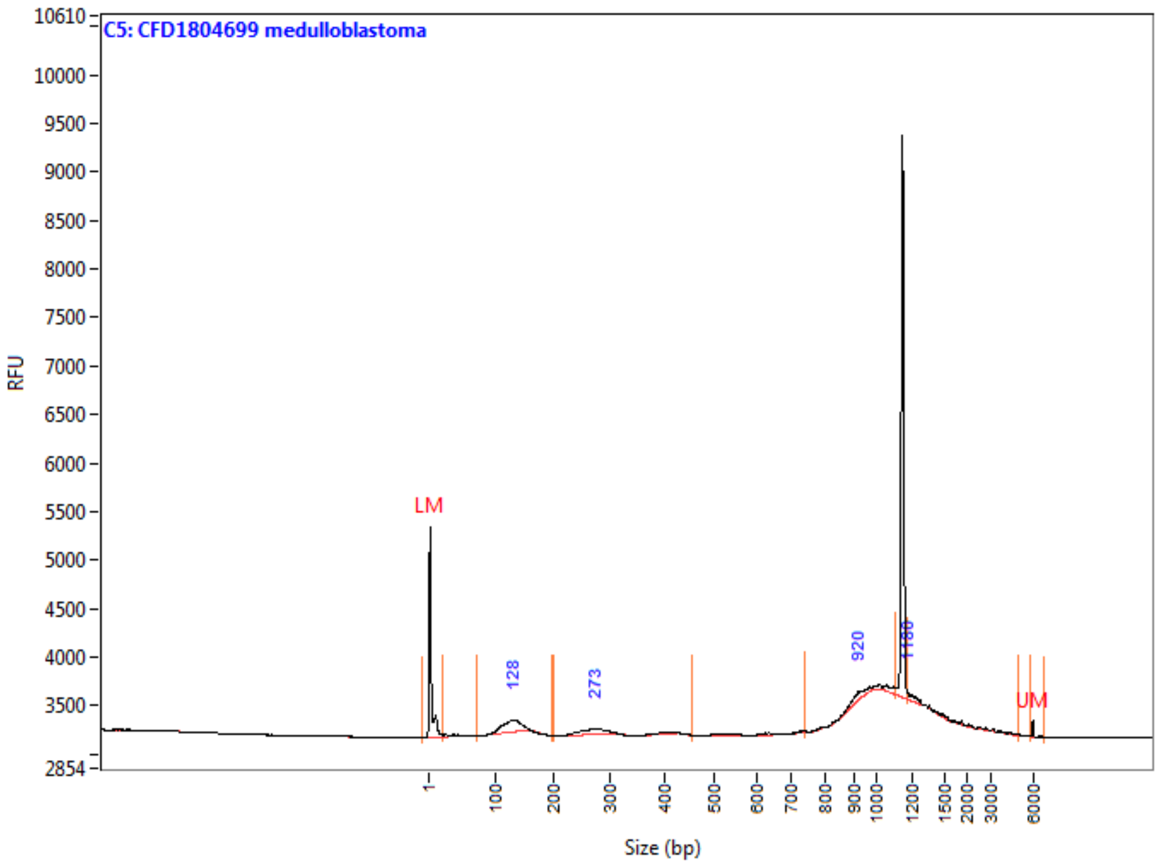

**Peak Table**

| | Size (bp) | ng/uL | % (Conc.) (ng/uL) | nmole/L | From (bp) | To (bp) | Avg. Size | CV% |
|---|---|---|---|---|---|---|---|---|
| 1 | 1 (LM) | 0.0004 | | 0.3108 | 0 | 21 | 2 | 209.34 |
| 2 | 128 | 0.0016 | 40.7 | 0.0219 | 72 | 196 | 123 | 11.29 |
| 3 | 273 | 0.0010 | 25.2 | 0.0059 | 202 | 455 | 285 | 17.21 |
| 4 | 920 | 0.0009 | 21.7 | 0.0015 | 738 | 1108 | 964 | 8.56 |
| 5 | 1180 | 0.0005 | 12.5 | 0.0004 | 1180 | 4938 | 1925 | 51.65 |
| 6 | 6000 (UM) | 0.0000 | | 0.0000 | 5875 | 6743 | 6026 | 3.06 |
| | TIC: | 0.0040 | ng/uL | | | | | |
| | TIM: | 0.0297 | nmole/L | | | | | |
| | Total Conc.: | 0.0102 | ng/uL | | | | | |

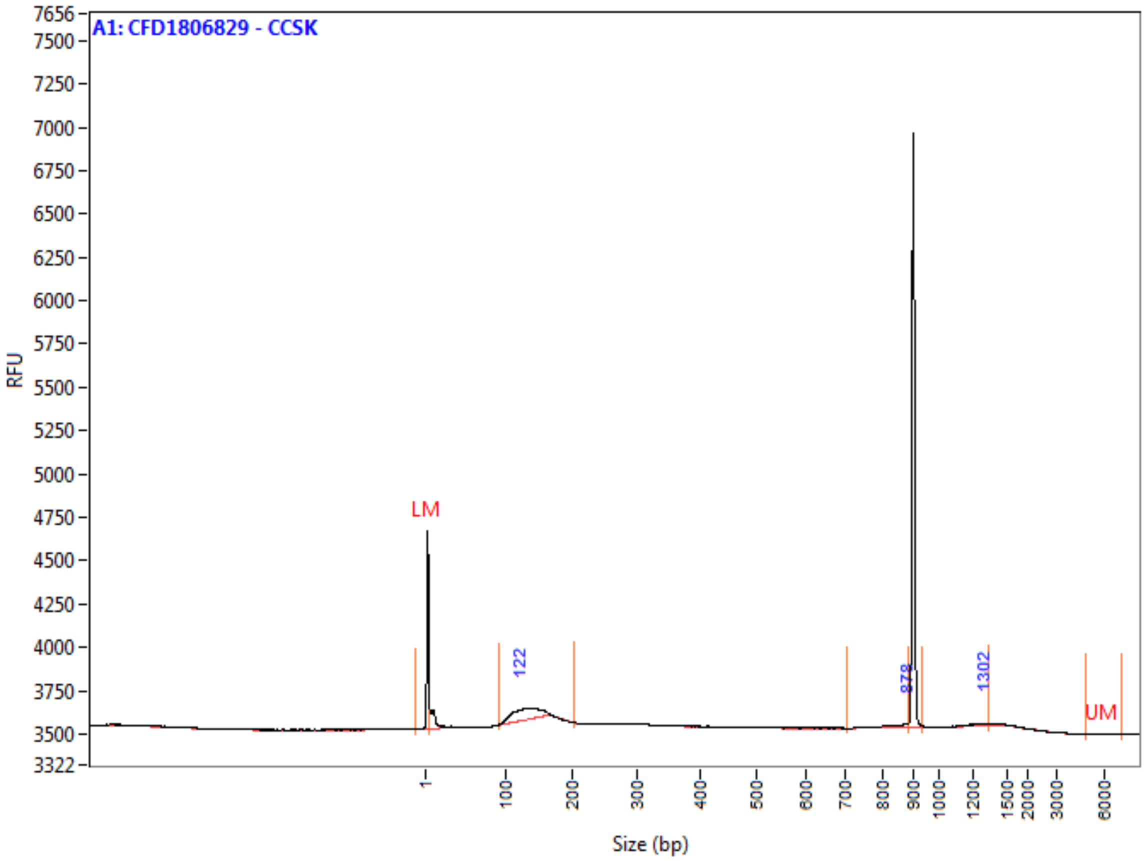

**Peak Table**

| | Size (bp) | ng/uL | % (Conc.) (ng/uL) | nmole/L | From (bp) | To (bp) | Avg. Size | CV% |
|---|---|---|---|---|---|---|---|---|
| 1 | 1 (LM) | 0.0007 | | 0.9609 | 0 | 6 | 1 | 135.64 |
| 2 | 122 | 0.0050 | 86.6 | 0.0640 | 93 | 204 | 128 | 12.95 |
| 3 | 878 | 0.0004 | 7.5 | 0.0008 | 706 | 885 | 838 | 4.77 |
| 4 | 1302 | 0.0003 | 5.9 | 0.0005 | 937 | 1344 | 1192 | 10.17 |
| 5 | 6000 (UM) | 0.0000 | | 0.0000 | 4878 | 7228 | 6163 | 7.88 |
| | TIC: | 0.0058 | ng/uL | | | | | |
| | TIM: | 0.0653 | nmole/L | | | | | |
| | Total Conc.: | 0.0210 | ng/uL | | | | | |

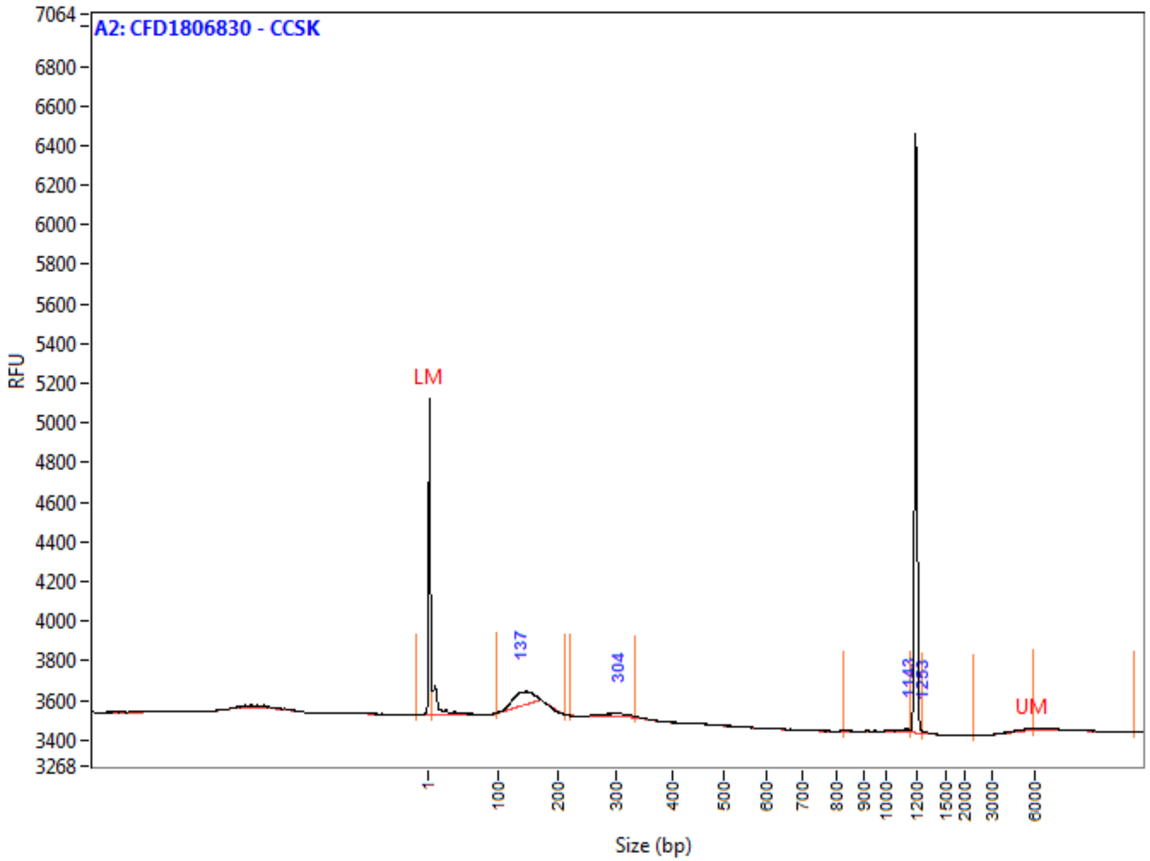

**Peak Table**

| | Size (bp) | ng/uL | % (Conc.) (ng/uL) | nmole/L | From (bp) | To (bp) | Avg. Size | CV% |
|---|---|---|---|---|---|---|---|---|
| 1 | 1 (LM) | 0.0007 | | 0.9609 | 0 | 6 | 1 | 164.93 |
| 2 | 137 | 0.0027 | 73.0 | 0.0322 | 98 | 212 | 137 | 10.14 |
| 3 | 304 | 0.0007 | 19.1 | 0.0039 | 219 | 335 | 295 | 6.75 |
| 4 | 1143 | 0.0002 | 6.1 | 0.0004 | 826 | 1161 | 1027 | 11.02 |
| 5 | 1253 | 0.0001 | 1.7 | 0.0001 | 1253 | 2318 | 1378 | 13.61 |
| 6 | 6000 (UM) | 0.0000 | | 0.0000 | 6000 | 13301 | 7163 | 22.08 |
| | TIC: | 0.0037 | ng/uL | | | | | |
| | TIM: | 0.0365 | nmole/L | | | | | |
| | Total Conc.: | 0.0137 | ng/uL | | | | | |

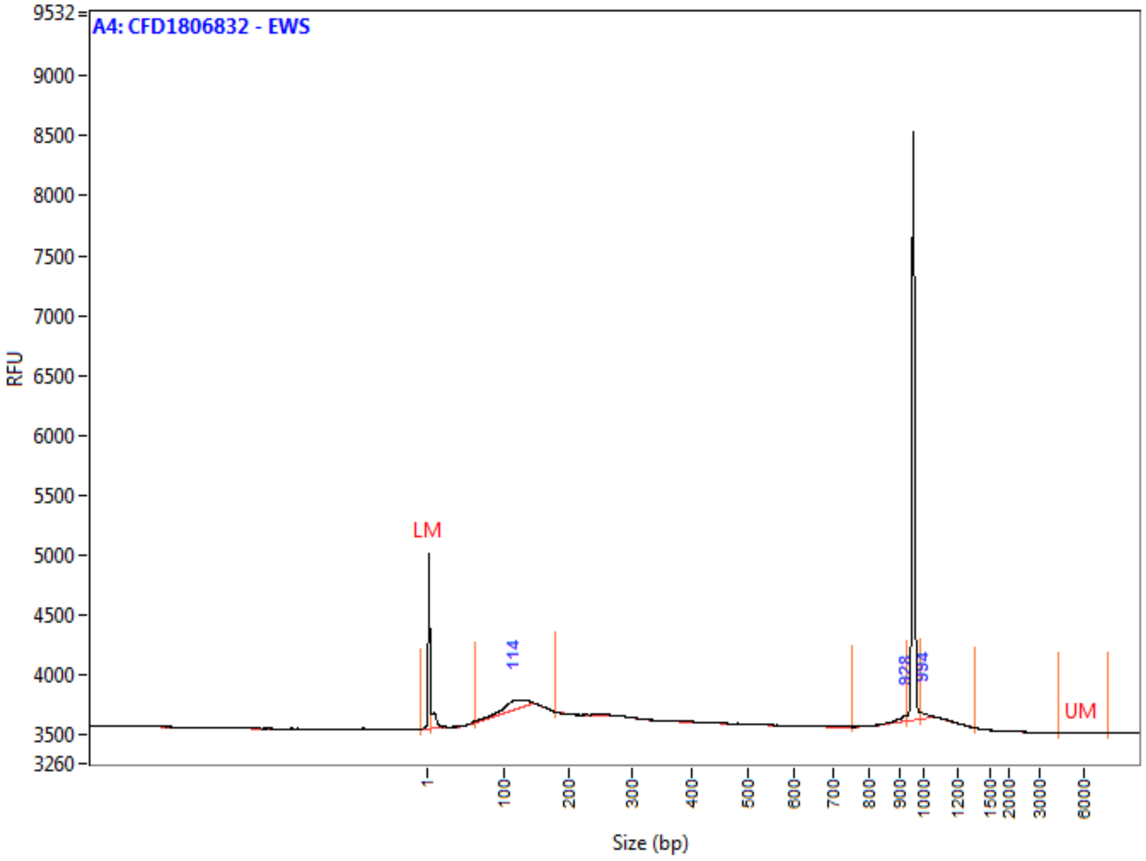

**Peak Table**

| | Size (bp) | ng/uL | % (Conc.) (ng/uL) | nmole/L | From (bp) | To (bp) | Avg. Size | CV% |
|---|---|---|---|---|---|---|---|---|
| 1 | 1 (LM) | 0.0007 | | 0.9609 | 0 | 5 | 1 | 157.30 |
| 2 | 114 | 0.0041 | 80.3 | 0.0612 | 63 | 180 | 109 | 16.71 |
| 3 | 928 | 0.0006 | 11.0 | 0.0010 | 751 | 931 | 889 | 4.57 |
| 4 | 994 | 0.0004 | 8.7 | 0.0007 | 989 | 1370 | 1027 | 4.47 |
| 5 | 6000 (UM) | 0.0000 | | 0.0000 | 4352 | 7683 | 6084 | 10.64 |
| | TIC: | 0.0051 | ng/uL | | | | | |
| | TIM: | 0.0629 | nmole/L | | | | | |
| | Total Conc.: | 0.0240 | ng/uL | | | | | |

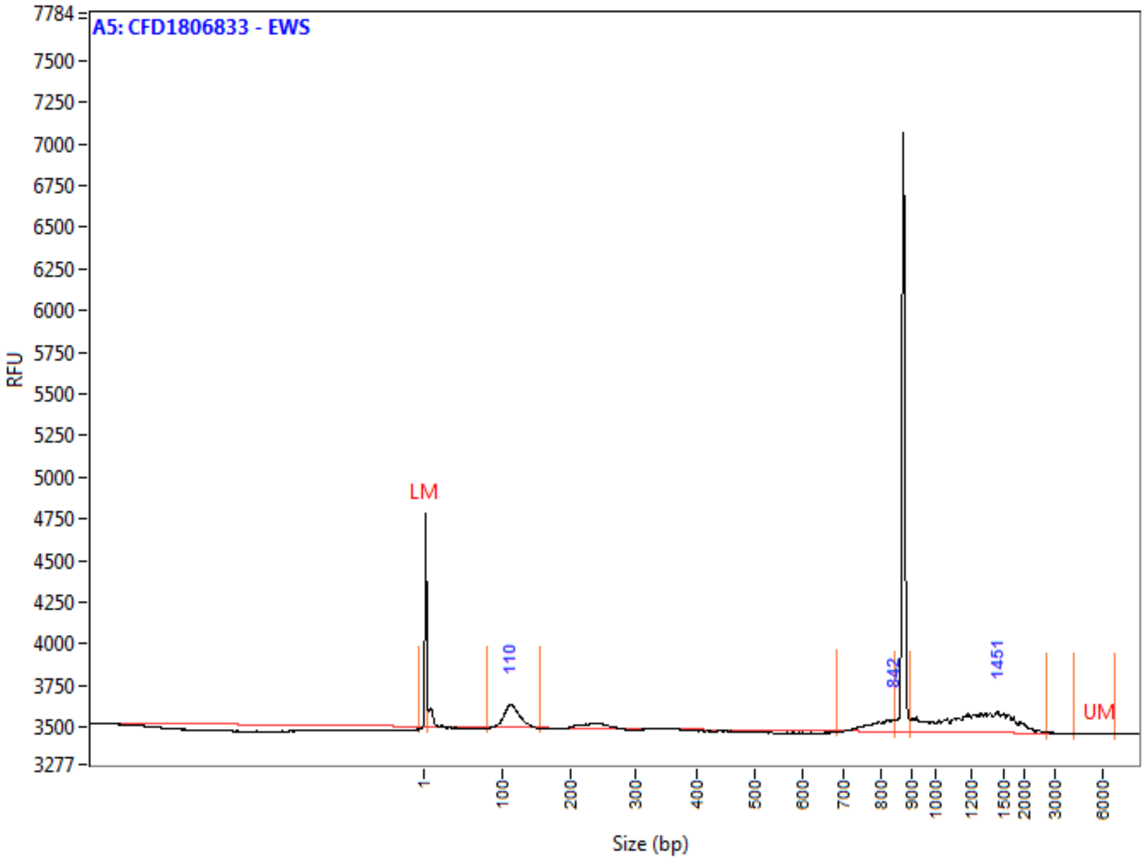

**Peak Table**

| | Size (bp) | ng/uL | % (Conc.) (ng/uL) | nmole/L | From (bp) | To (bp) | Avg. Size | CV% |
|---|---|---|---|---|---|---|---|---|
| 1 | 1 (LM) | 0.0007 | | 0.9609 | 0 | 5 | 1 | 117.16 |
| 2 | 110 | 0.0052 | 29.3 | 0.0764 | 80 | 154 | 112 | 9.47 |
| 3 | 842 | 0.0022 | 12.2 | 0.0045 | 685 | 847 | 796 | 4.22 |
| 4 | 1451 | 0.0104 | 58.5 | 0.0131 | 897 | 2741 | 1304 | 24.68 |
| 5 | 6000 (UM) | 0.0000 | | 0.0000 | 4260 | 6933 | 5536 | 11.45 |
| | TIC: | 0.0178 | ng/uL | | | | | |
| | TIM: | 0.0940 | nmole/L | | | | | |
| | Total Conc.: | 0.0362 | ng/uL | | | | | |

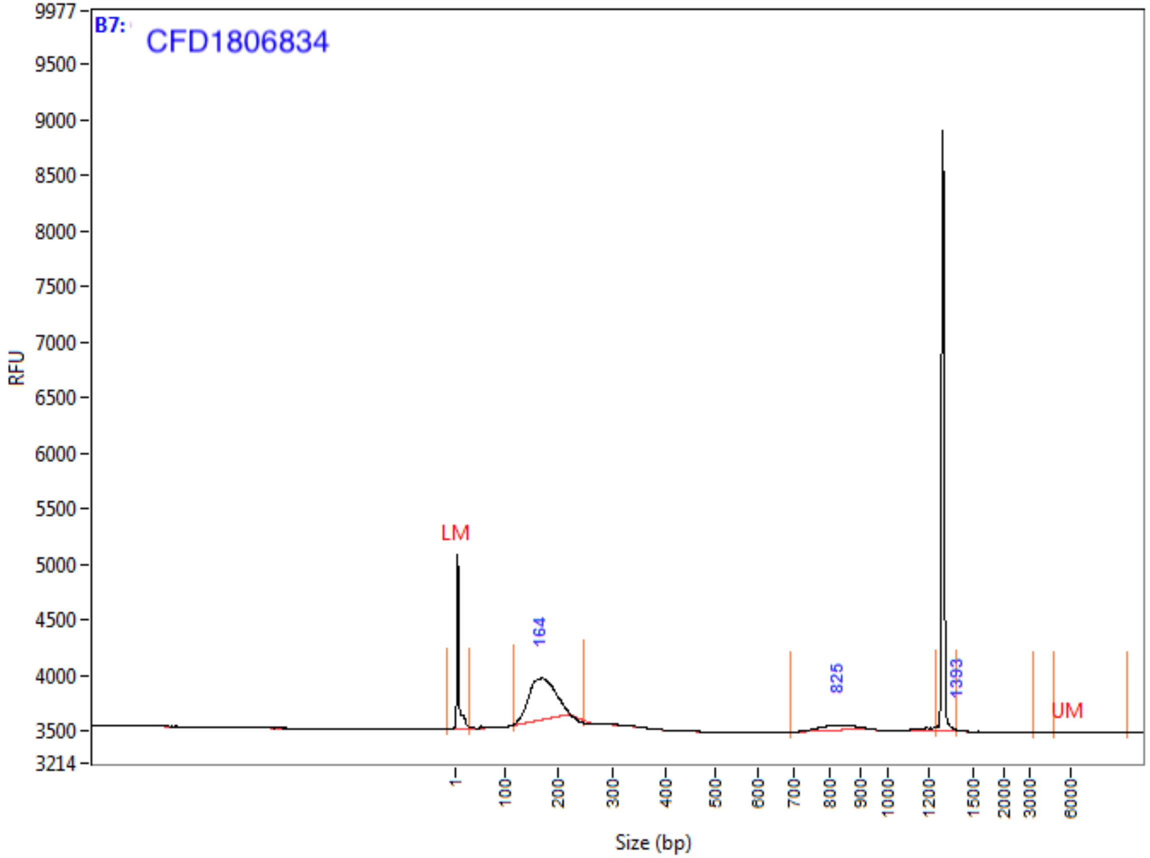

**Peak Table**

| | Size (bp) | ng/uL | % (Conc.) (ng/uL) | nmole/L | From (bp) | To (bp) | Avg. Size | CV% |
|---|---|---|---|---|---|---|---|---|
| 1 | 1 (LM) | 0.0013 | | 0.9492 | 0 | 26 | 2 | 227.41 |
| 2 | 164 | 0.0314 | 89.7 | 0.3094 | 116 | 245 | 167 | 12.37 |
| 3 | 825 | 0.0035 | 10.1 | 0.0065 | 692 | 1252 | 891 | 17.00 |
| 4 | 1393 | 0.0001 | 0.3 | 0.0001 | 1393 | 3304 | 1720 | 21.18 |
| 5 | 6000 (UM) | 0.0000 | | 0.0000 | 4819 | 10183 | 7507 | 19.48 |
| | TIC: | 0.0351 | ng/uL | | | | | |
| | TIM: | 0.3160 | nmole/L | | | | | |
| | Total Conc.: | 0.0578 | ng/uL | | | | | |

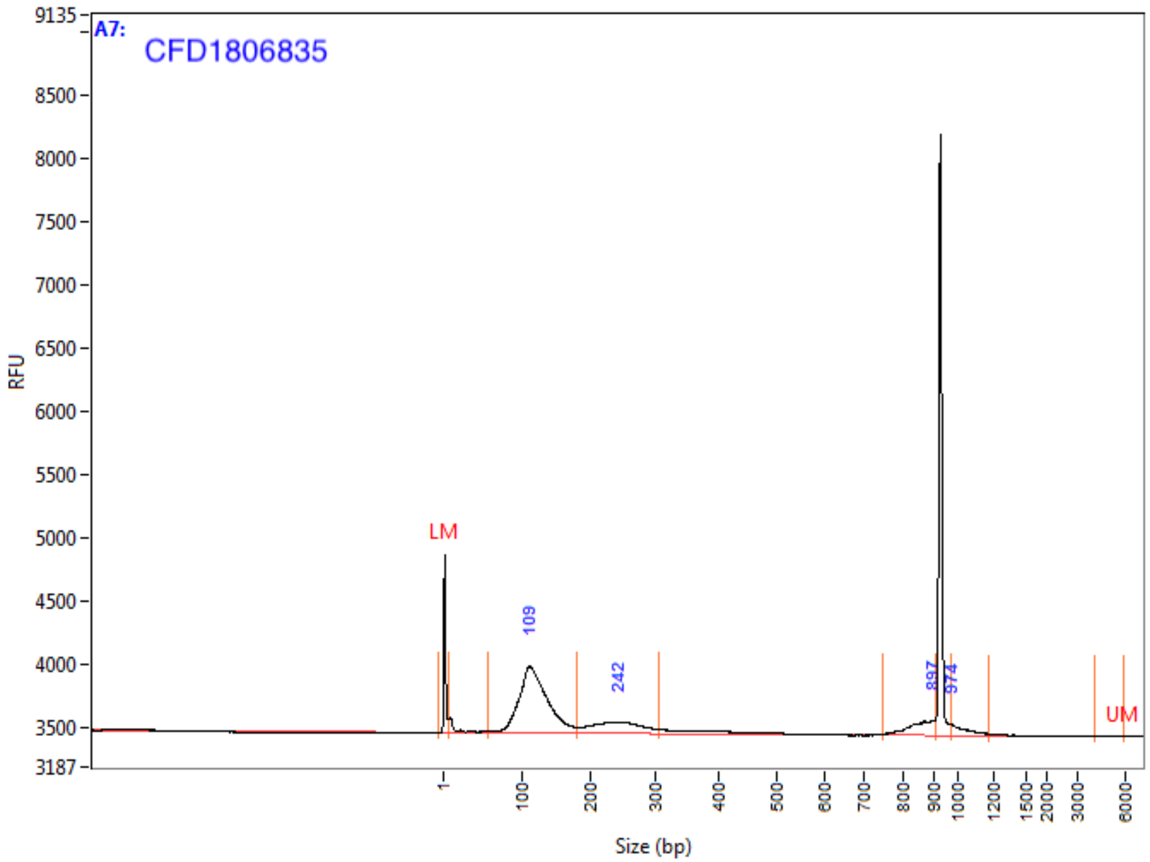

Peak Table

| | Size (bp) | ng/uL | % (Conc.) (ng/uL) | nmole/L | From (bp) | To (bp) | Avg. Size | CV% |
|---|---|---|---|---|---|---|---|---|
| 1 | 1 (LM) | 0.0007 | | 0.9609 | 0 | 7 | 1 | 180.54 |
| 2 | 109 | 0.0322 | 71.1 | 0.4521 | 57 | 181 | 117 | 18.58 |
| 3 | 242 | 0.0083 | 18.3 | 0.0569 | 181 | 306 | 240 | 13.24 |
| 4 | 897 | 0.0033 | 7.3 | 0.0064 | 750 | 911 | 853 | 4.53 |
| 5 | 974 | 0.0014 | 3.2 | 0.0023 | 974 | 1180 | 1041 | 5.34 |
| 6 | 6000 (UM) | 0.0000 | | 0.0000 | 4146 | 6023 | 5940 | 2.40 |
| | TIC: | 0.0453 | ng/uL | | | | | |
| | TIM: | 0.5177 | nmole/L | | | | | |
| | Total Conc.: | 0.0665 | ng/uL | | | | | |

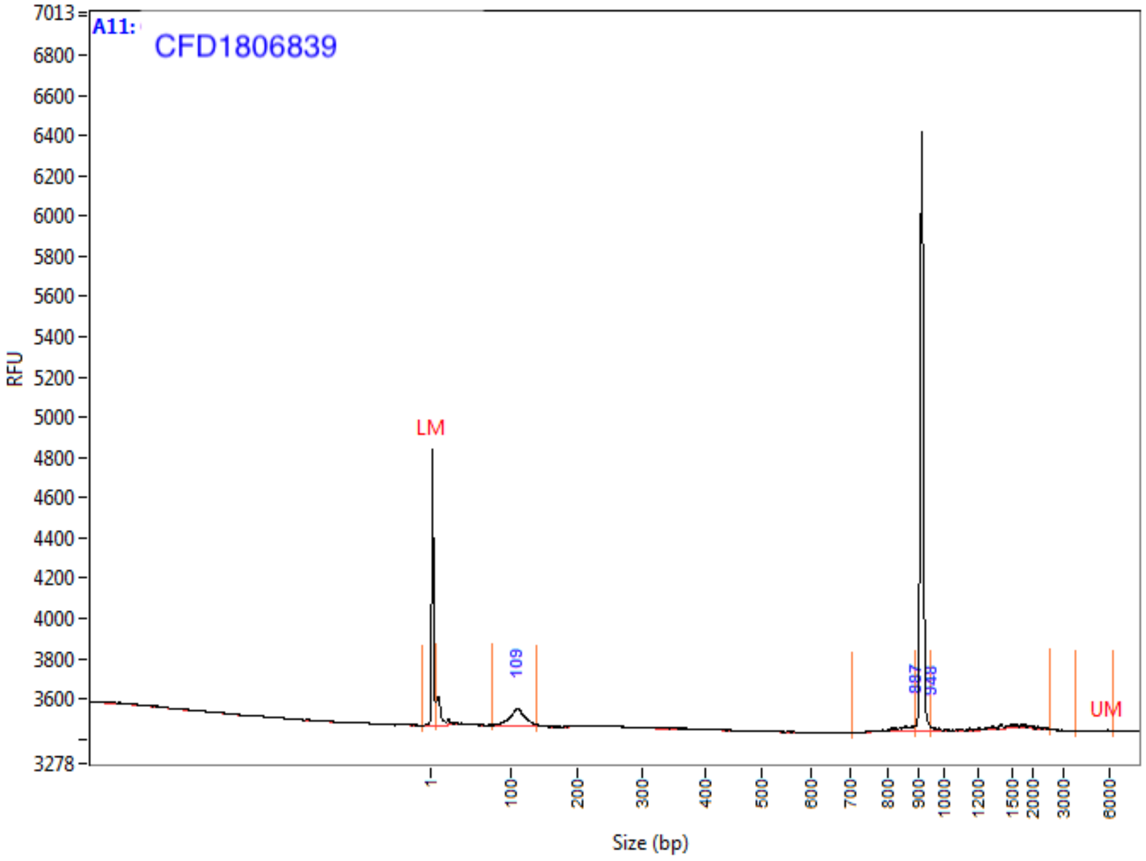

Peak Table

| | Size (bp) | ng/uL | % (Conc.) (ng/uL) | nmole/L | From (bp) | To (bp) | Avg. Size | CV% |
|---|---|---|---|---|---|---|---|---|
| 1 | 1 (LM) | 0.0007 | | 0.9609 | 0 | 7 | 1 | 137.00 |
| 2 | 109 | 0.0027 | 67.8 | 0.0406 | 78 | 140 | 109 | 10.53 |
| 3 | 887 | 0.0005 | 11.6 | 0.0009 | 705 | 888 | 840 | 4.76 |
| 4 | 948 | 0.0008 | 20.6 | 0.0009 | 948 | 2612 | 1432 | 28.15 |
| 5 | 6000 (UM) | 0.0000 | | 0.0000 | 3871 | 6318 | 5709 | 8.92 |
| | TIC: | 0.0040 | ng/uL | | | | | |
| | TIM: | 0.0424 | nmole/L | | | | | |
| | Total Conc.: | 0.0172 | ng/uL | | | | | |

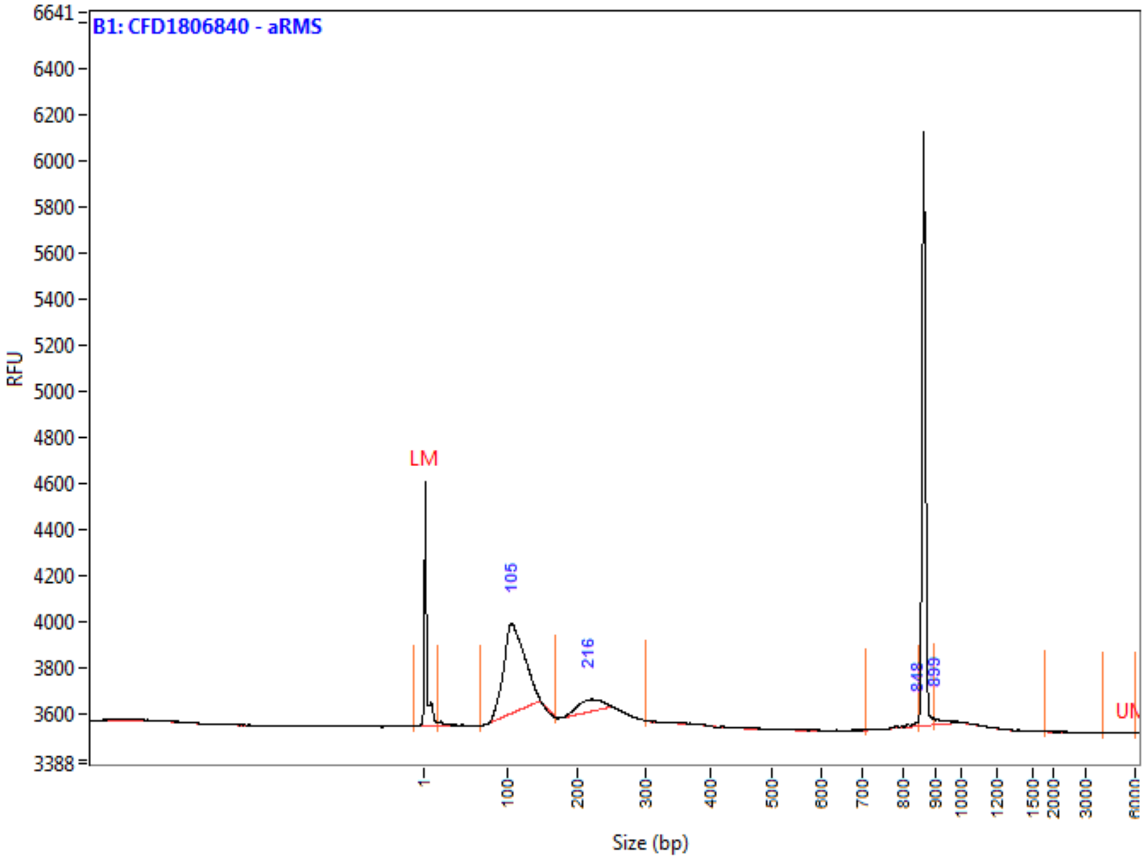

**Peak Table**

| | Size (bp) | ng/uL | % (Conc.) (ng/uL) | nmole/L | From (bp) | To (bp) | Avg. Size | CV% |
|---|---|---|---|---|---|---|---|---|
| 1 | 1 (LM) | 0.0007 | | 0.5357 | 0 | 18 | 2 | 180.03 |
| 2 | 105 | 0.0189 | 86.0 | 0.2824 | 68 | 170 | 110 | 12.21 |
| 3 | 216 | 0.0025 | 11.6 | 0.0194 | 168 | 303 | 216 | 6.94 |
| 4 | 848 | 0.0002 | 1.0 | 0.0004 | 708 | 850 | 812 | 4.32 |
| 5 | 899 | 0.0003 | 1.5 | 0.0006 | 899 | 1816 | 959 | 12.41 |
| 6 | 6000 (UM) | 0.0000 | | 0.0000 | 4123 | 6136 | 5413 | 10.94 |
| | TIC: | 0.0220 | ng/uL | | | | | |
| | TIM: | 0.3028 | nmole/L | | | | | |
| | Total Conc.: | 0.0329 | ng/uL | | | | | |

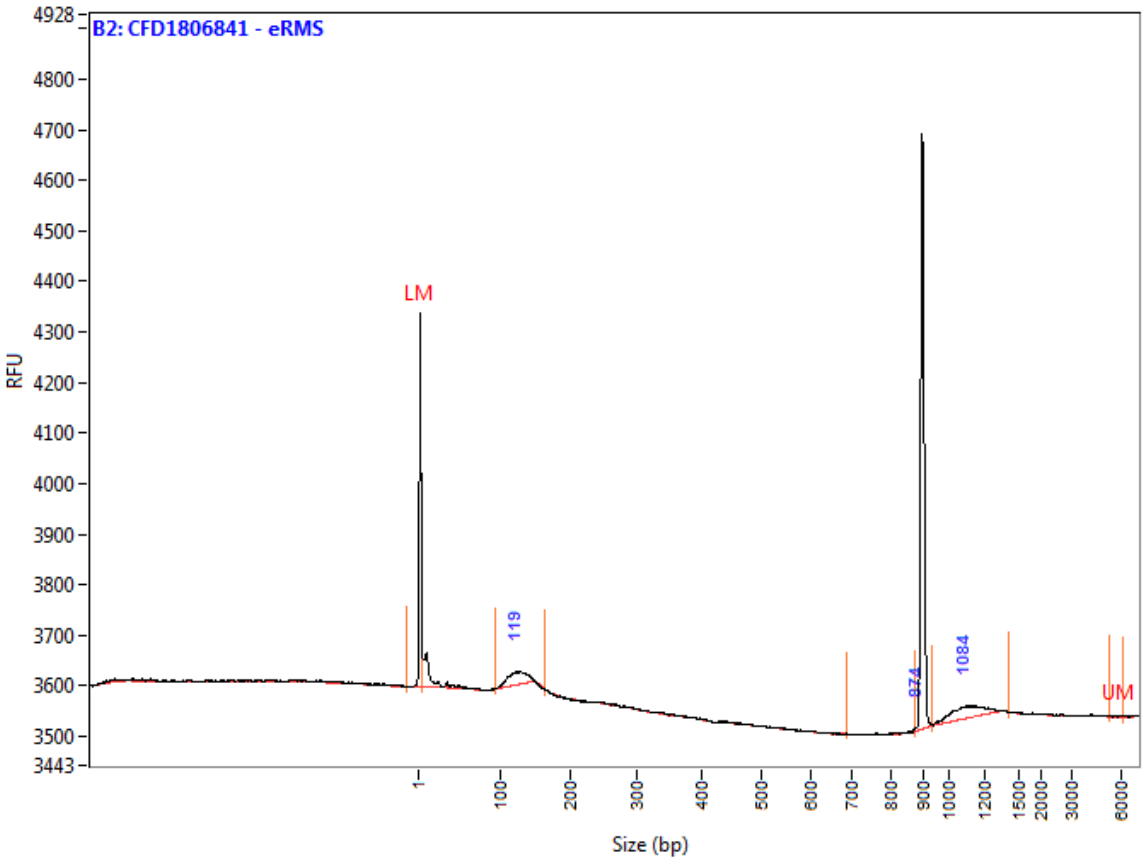

**Peak Table**

| | Size (bp) | ng/uL | % (Conc.) (ng/uL) | nmole/L | From (bp) | To (bp) | Avg. Size | CV% |
|---|---|---|---|---|---|---|---|---|
| 1 | 1 (LM) | 0.0007 | | 0.9609 | 0 | 6 | 1 | 184.05 |
| 2 | 119 | 0.0019 | 54.9 | 0.0258 | 94 | 165 | 121 | 9.26 |
| 3 | 874 | 0.0001 | 2.4 | 0.0002 | 687 | 875 | 799 | 6.80 |
| 4 | 1084 | 0.0015 | 42.7 | 0.0022 | 935 | 1412 | 1094 | 7.25 |
| 5 | 6000 (UM) | 0.0000 | | 0.0000 | 5382 | 6250 | 6056 | 2.51 |
| | TIC: | 0.0035 | ng/uL | | | | | |
| | TIM: | 0.0282 | nmole/L | | | | | |
| | Total Conc.: | 0.0142 | ng/uL | | | | | |

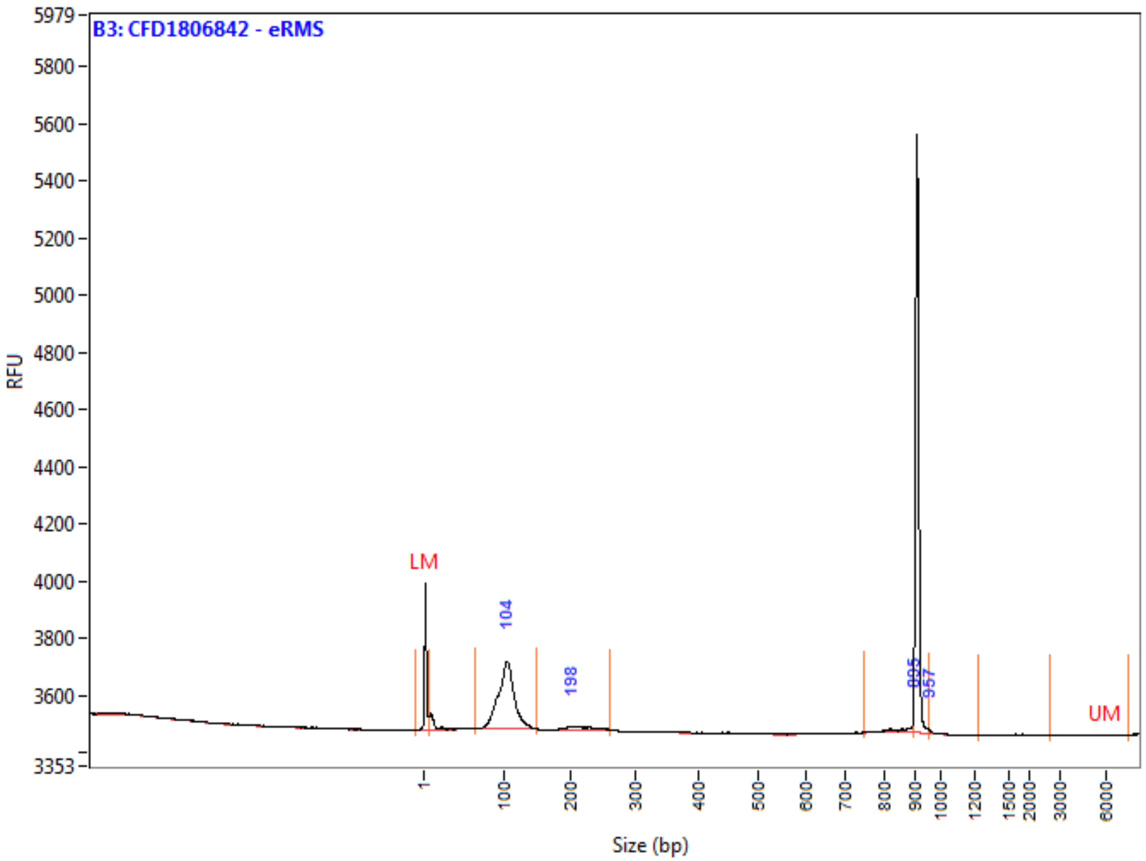

**Peak Table**

| | Size (bp) | ng/uL | % (Conc.) (ng/uL) | nmole/L | From (bp) | To (bp) | Avg. Size | CV% |
|---|---|---|---|---|---|---|---|---|
| 1 | 1 (LM) | 0.0007 | | 0.9609 | 0 | 7 | 1 | 170.81 |
| 2 | 104 | 0.0217 | 90.5 | 0.3486 | 64 | 148 | 102 | 11.70 |
| 3 | 198 | 0.0015 | 6.2 | 0.0116 | 150 | 259 | 212 | 9.01 |
| 4 | 895 | 0.0006 | 2.5 | 0.0012 | 749 | 897 | 863 | 4.59 |
| 5 | 957 | 0.0002 | 0.8 | 0.0003 | 957 | 1234 | 1033 | 7.39 |
| 6 | 6000 (UM) | 0.0000 | | 0.0000 | 2718 | 7478 | 5023 | 29.78 |
| | TIC: | 0.0239 | ng/uL | | | | | |
| | TIM: | 0.3616 | nmole/L | | | | | |
| | Total Conc.: | 0.0470 | ng/uL | | | | | |

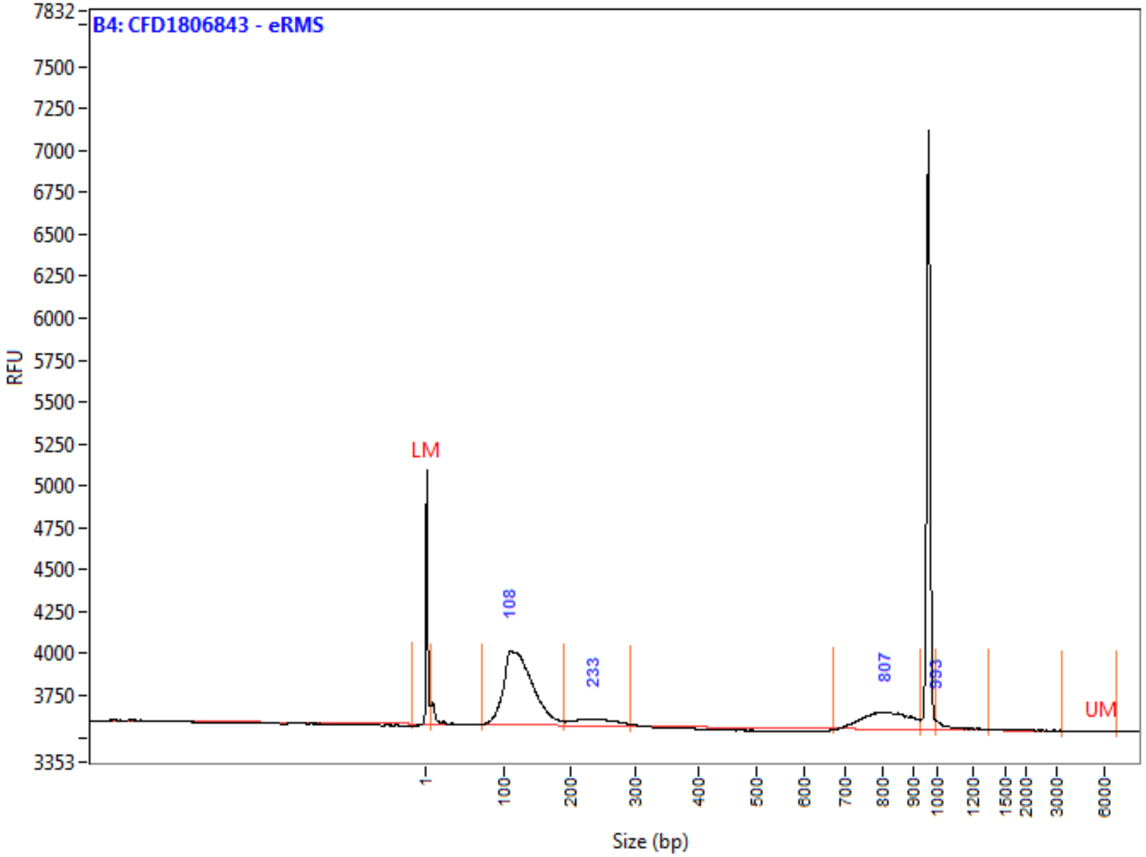

**Peak Table**

| | Size (bp) | ng/uL | % (Conc.) (ng/uL) | nmole/L | From (bp) | To (bp) | Avg. Size | CV% |
|---|---|---|---|---|---|---|---|---|
| 1 | 1 (LM) | 0.0007 | | 0.9609 | 0 | 7 | 1 | 179.32 |
| 2 | 108 | 0.0251 | 75.0 | 0.3375 | 72 | 189 | 122 | 17.12 |
| 3 | 233 | 0.0028 | 8.3 | 0.0194 | 189 | 292 | 235 | 10.90 |
| 4 | 807 | 0.0050 | 14.9 | 0.0100 | 672 | 931 | 815 | 7.01 |
| 5 | 993 | 0.0006 | 1.9 | 0.0009 | 993 | 1350 | 1084 | 8.50 |
| 6 | 6000 (UM) | 0.0000 | | 0.0000 | 3367 | 6933 | 5112 | 20.40 |
| | TIC: | 0.0334 | ng/uL | | | | | |
| | TIM: | 0.3678 | nmole/L | | | | | |
| | Total Conc.: | 0.0478 | ng/uL | | | | | |

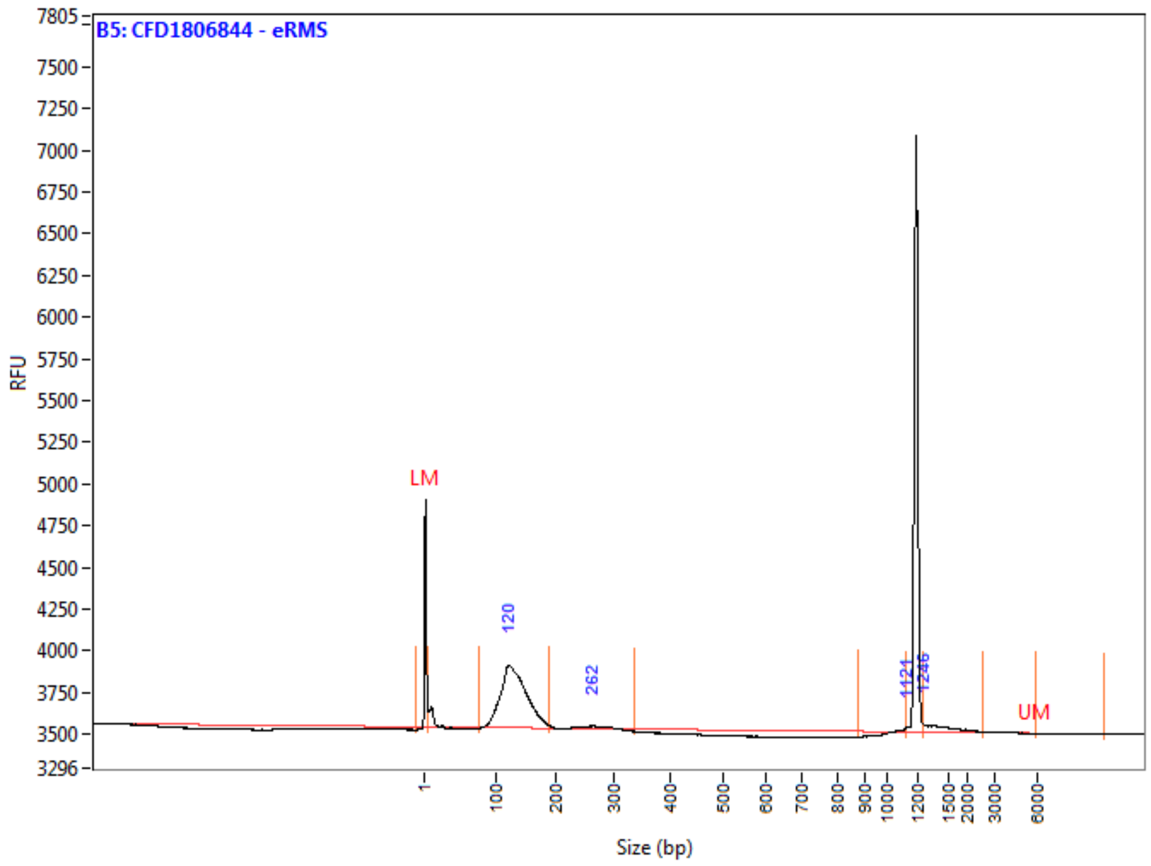

**Peak Table**

| | Size (bp) | ng/uL | % (Conc.) (ng/uL) | nmole/L | From (bp) | To (bp) | Avg. Size | CV% |
|---|---|---|---|---|---|---|---|---|
| 1 | 1 (LM) | 0.0007 | | 0.9609 | 0 | 6 | 1 | 131.77 |
| 2 | 120 | 0.0229 | 91.5 | 0.2891 | 78 | 189 | 130 | 15.61 |
| 3 | 262 | 0.0007 | 2.9 | 0.0046 | 189 | 337 | 261 | 8.17 |
| 4 | 1121 | 0.0001 | 0.4 | 0.0001 | 877 | 1123 | 1100 | 1.58 |
| 5 | 1246 | 0.0013 | 5.3 | 0.0014 | 1246 | 2600 | 1533 | 18.83 |
| 6 | 6000 (UM) | | | | 6000 | 10890 | NaN | NaN |
| | TIC: | 0.0250 | ng/uL | | | | | |
| | TIM: | 0.2952 | nmole/L | | | | | |
| | Total Conc.: | 0.0423 | ng/uL | | | | | |

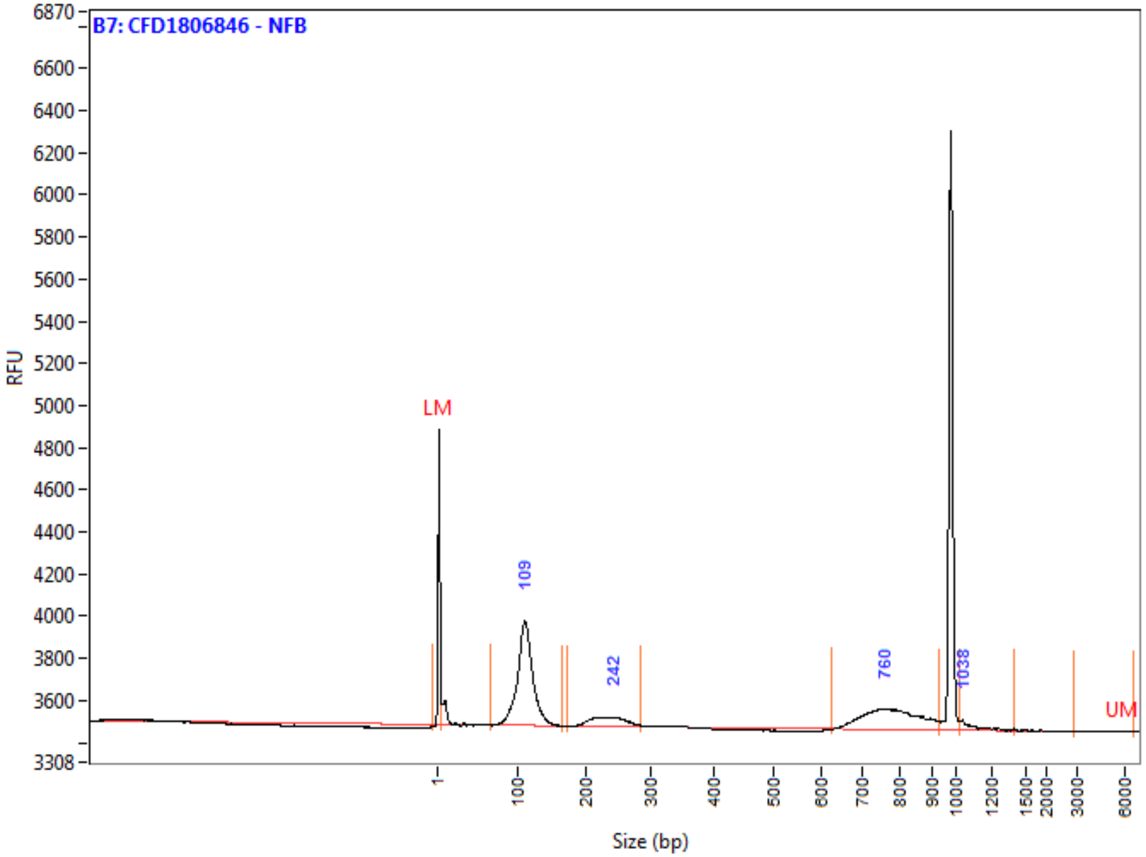

**Peak Table**

| | Size (bp) | ng/uL | % (Conc.) (ng/uL) | nmole/L | From (bp) | To (bp) | Avg. Size | CV% |
|---|---|---|---|---|---|---|---|---|
| 1 | 1 (LM) | 0.0007 | | 0.9609 | 0 | 5 | 1 | 191.18 |
| 2 | 109 | 0.0170 | 61.8 | 0.2558 | 67 | 166 | 109 | 11.00 |
| 3 | 242 | 0.0029 | 10.6 | 0.0206 | 172 | 283 | 233 | 9.26 |
| 4 | 760 | 0.0069 | 24.9 | 0.0144 | 625 | 933 | 781 | 8.68 |
| 5 | 1038 | 0.0007 | 2.6 | 0.0011 | 1019 | 1409 | 1115 | 8.34 |
| 6 | 6000 (UM) | 0.0000 | | 0.0000 | 2929 | 6660 | 5391 | 21.47 |
| | TIC: | 0.0275 | ng/uL | | | | | |
| | TIM: | 0.2919 | nmole/L | | | | | |
| | Total Conc.: | 0.0421 | ng/uL | | | | | |

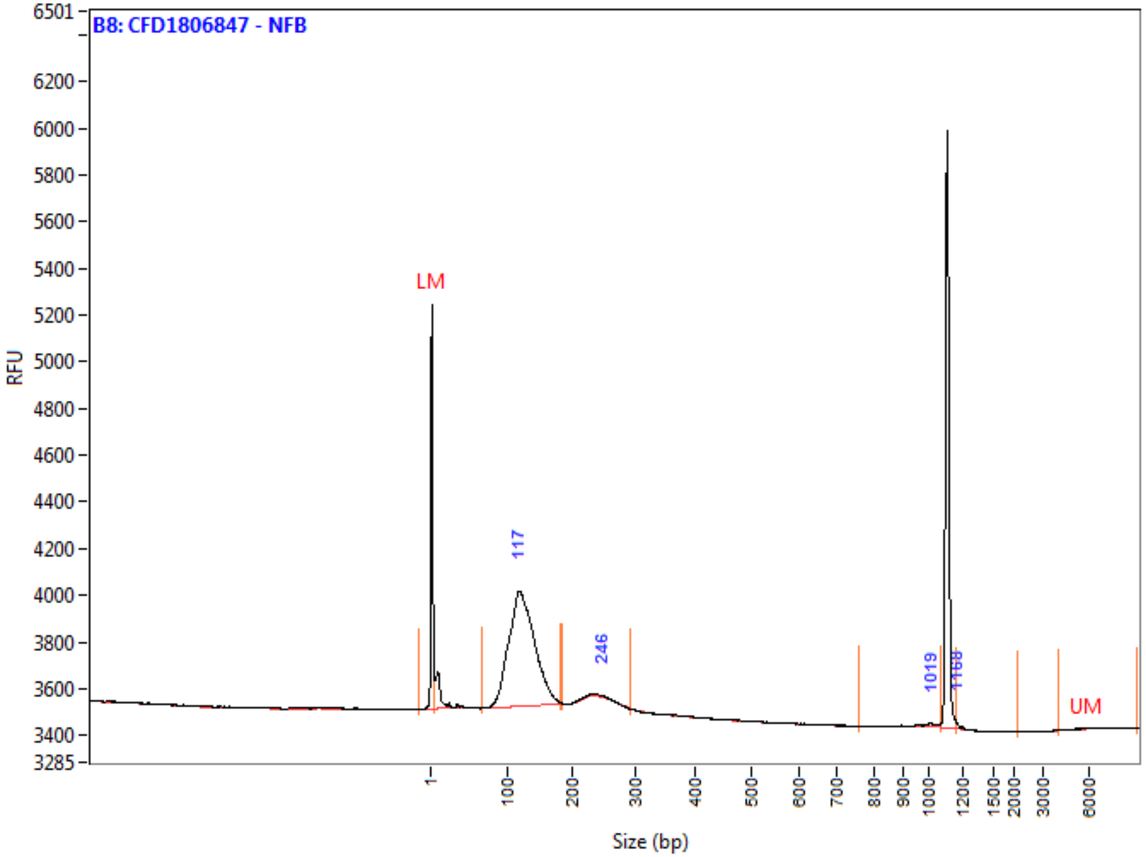

**Peak Table**

| | Size (bp) | ng/uL | % (Conc.) (ng/uL) | nmole/L | From (bp) | To (bp) | Avg. Size | CV% |
|---|---|---|---|---|---|---|---|---|
| 1 | 1 (LM) | 0.0007 | | 0.9609 | 0 | 7 | 1 | 144.87 |
| 2 | 117 | 0.0217 | 98.2 | 0.2928 | 68 | 183 | 122 | 15.28 |
| 3 | 246 | 0.0001 | 0.6 | 0.0010 | 186 | 294 | 233 | 5.76 |
| 4 | 1019 | 0.0002 | 0.9 | 0.0003 | 760 | 1076 | 978 | 8.43 |
| 5 | 1168 | 0.0001 | 0.3 | 0.0001 | 1168 | 2154 | 1237 | 11.68 |
| 6 | 6000 (UM) | 0.0000 | | 0.0000 | 4031 | 9298 | 6978 | 17.17 |
| | TIC: | 0.0222 | ng/uL | | | | | |
| | TIM: | 0.2942 | nmole/L | | | | | |
| | Total Conc.: | 0.0316 | ng/uL | | | | | |

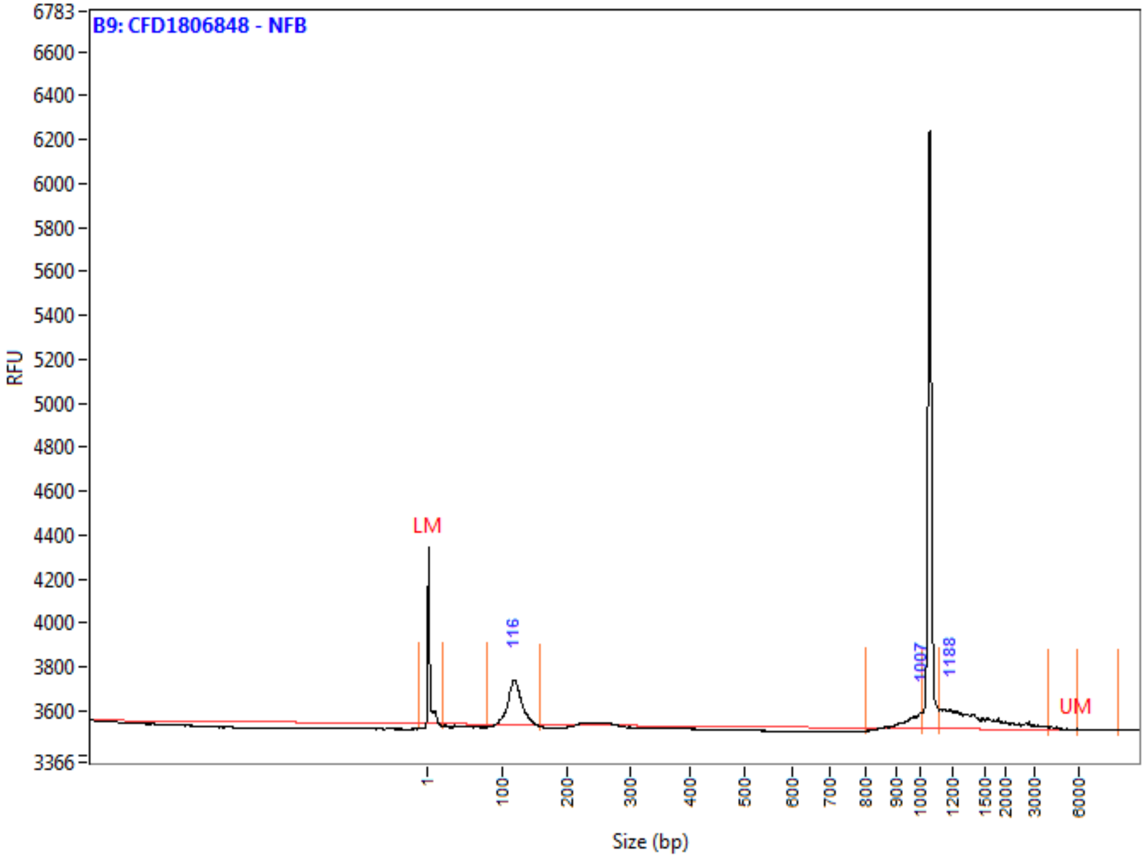

**Peak Table**

| | Size (bp) | ng/uL | % (Conc.) (ng/uL) | nmole/L | From (bp) | To (bp) | Avg. Size | CV% |
|---|---|---|---|---|---|---|---|---|
| 1 | 1 (LM) | 0.0007 | | 0.5357 | 0 | 20 | 2 | 151.67 |
| 2 | 116 | 0.0095 | 52.8 | 0.1340 | 79 | 157 | 117 | 8.83 |
| 3 | 1007 | 0.0017 | 9.5 | 0.0029 | 800 | 1017 | 966 | 3.78 |
| 4 | 1188 | 0.0068 | 37.7 | 0.0070 | 1121 | 4008 | 1599 | 37.07 |
| 5 | 6000 (UM) | 0.0000 | | 0.0000 | 5977 | 8752 | 6836 | 15.05 |
| | TIC: | 0.0181 | ng/uL | | | | | |
| | TIM: | 0.1439 | nmole/L | | | | | |
| | Total Conc.: | 0.0375 | ng/uL | | | | | |

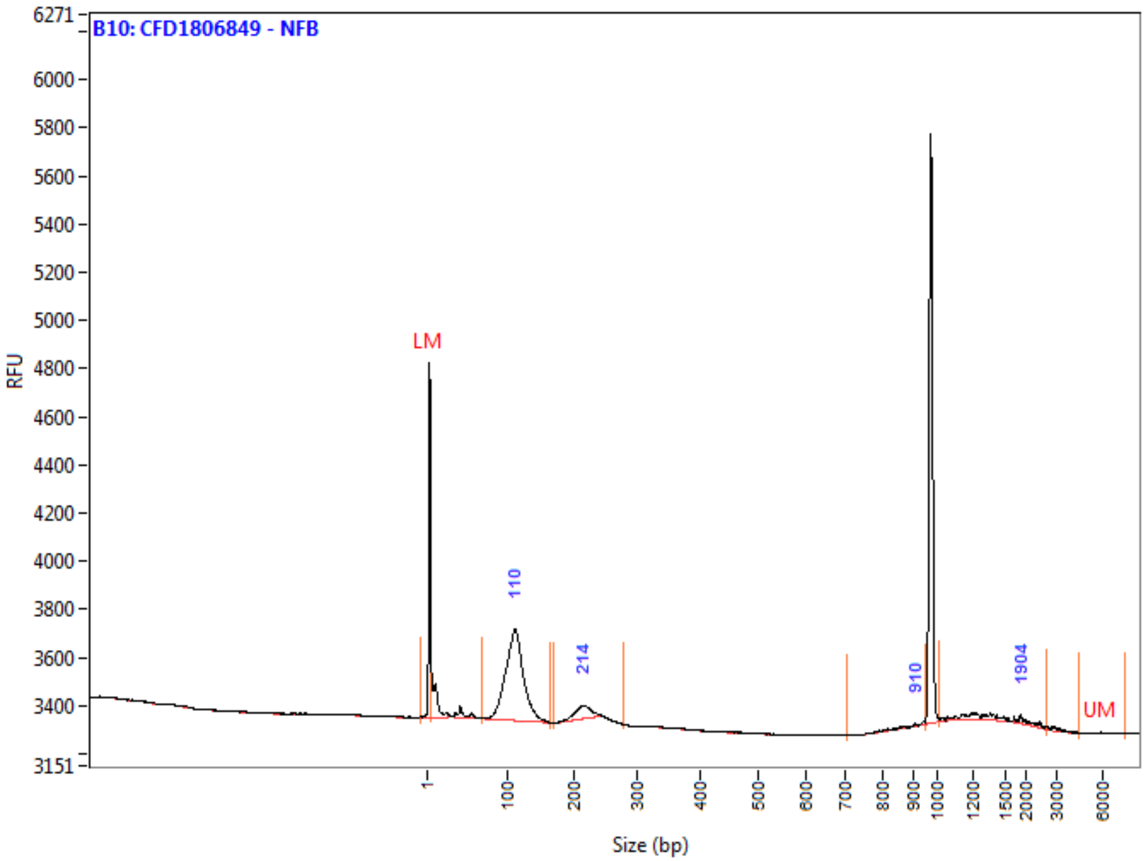

**Peak Table**

| | Size (bp) | ng/uL | % (Conc.) (ng/uL) | nmole/L | From (bp) | To (bp) | Avg. Size | CV% |
|---|---|---|---|---|---|---|---|---|
| 1 | 1 (LM) | 0.0007 | | 0.9609 | 0 | 5 | 1 | 167.43 |
| 2 | 110 | 0.0145 | 82.5 | 0.2168 | 68 | 166 | 110 | 12.16 |
| 3 | 214 | 0.0017 | 9.4 | 0.0128 | 169 | 277 | 213 | 5.73 |
| 4 | 910 | 0.0002 | 1.0 | 0.0003 | 707 | 953 | 870 | 7.11 |
| 5 | 1904 | 0.0012 | 7.1 | 0.0014 | 1011 | 2718 | 1512 | 28.89 |
| 6 | 6000 (UM) | 0.0000 | | 0.0000 | 4558 | 7569 | 5715 | 13.33 |
| | TIC: | 0.0176 | ng/uL | | | | | |
| | TIM: | 0.2313 | nmole/L | | | | | |
| | Total Conc.: | 0.0299 | ng/uL | | | | | |

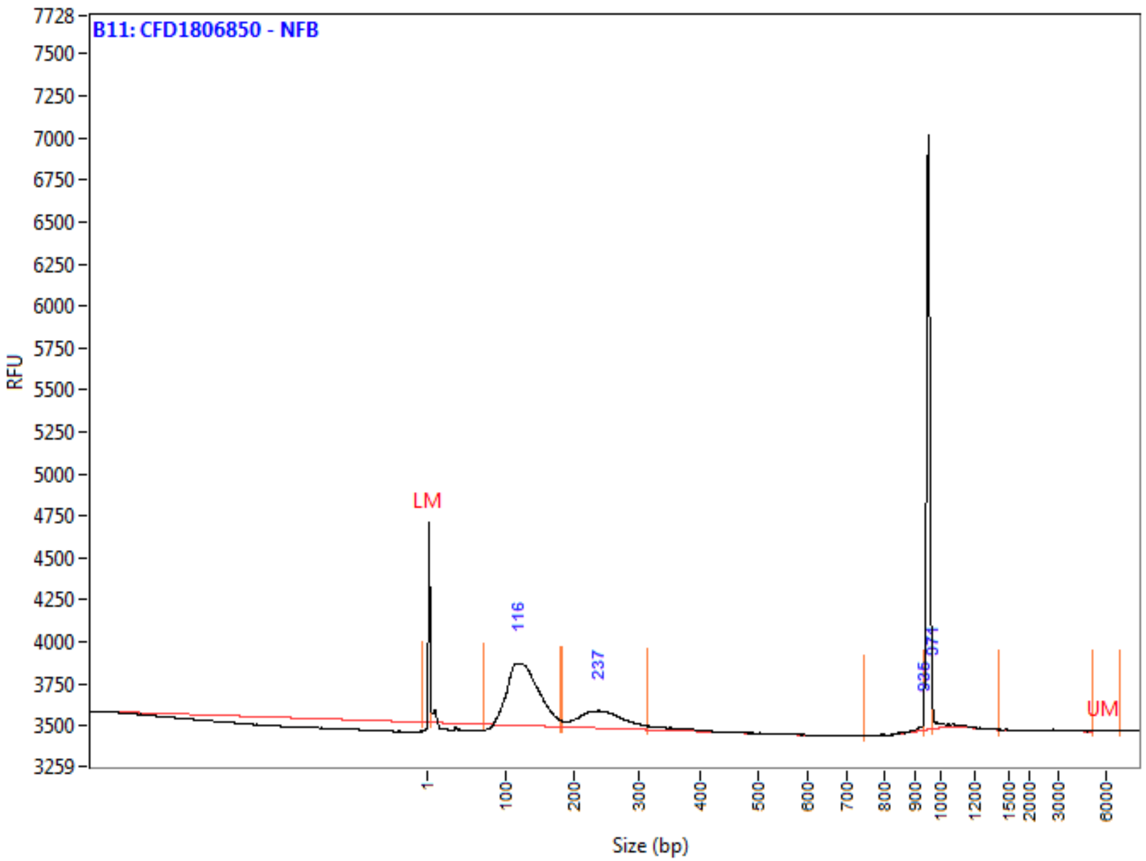

**Peak Table**

| | Size (bp) | ng/uL | % (Conc.) (ng/uL) | nmole/L | From (bp) | To (bp) | Avg. Size | CV% |
|---|---|---|---|---|---|---|---|---|
| 1 | 1 (LM) | 0.0007 | | 0.9609 | 0 | 6 | 1 | 114.75 |
| 2 | 116 | 0.0324 | 72.8 | 0.4254 | 72 | 181 | 125 | 16.54 |
| 3 | 237 | 0.0108 | 24.2 | 0.0729 | 184 | 316 | 243 | 12.46 |
| 4 | 935 | 0.0004 | 0.8 | 0.0007 | 747 | 937 | 892 | 5.89 |
| 5 | 971 | 0.0010 | 2.2 | 0.0016 | 971 | 1425 | 1034 | 8.16 |
| 6 | 6000 (UM) | 0.0000 | | 0.0000 | 5222 | 7001 | 6067 | 9.23 |
| | TIC: | 0.0445 | ng/uL | | | | | |
| | TIM: | 0.5006 | nmole/L | | | | | |
| | Total Conc.: | 0.0644 | ng/uL | | | | | |

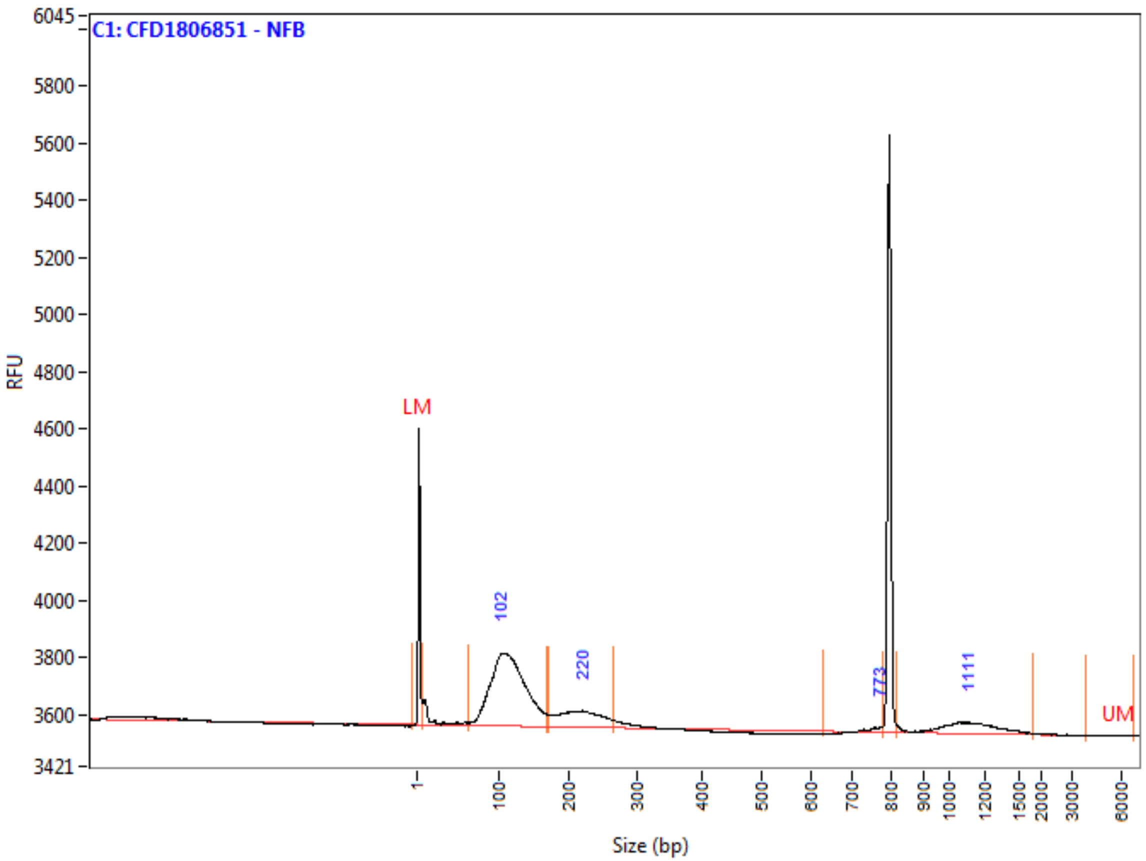

**Peak Table**

| | Size (bp) | ng/uL | % (Conc.) (ng/uL) | nmole/L | From (bp) | To (bp) | Avg. Size | CV% |
|---|---|---|---|---|---|---|---|---|
| 1 | 1 (LM) | 0.0007 | | 0.9487 | 0 | 6 | 1 | 149.08 |
| 2 | 102 | 0.0264 | 73.3 | 0.3866 | 62 | 168 | 112 | 19.74 |
| 3 | 220 | 0.0062 | 17.2 | 0.0474 | 171 | 265 | 215 | 11.67 |
| 4 | 773 | 0.0003 | 0.8 | 0.0007 | 629 | 778 | 755 | 2.19 |
| 5 | 1111 | 0.0031 | 8.6 | 0.0045 | 816 | 1825 | 1129 | 15.78 |
| 6 | 6000 (UM) | 0.0000 | | 0.0000 | 3871 | 6864 | 5711 | 14.37 |
| | TIC: | 0.0360 | ng/uL | | | | | |
| | TIM: | 0.4392 | nmole/L | | | | | |
| | Total Conc.: | 0.0509 | ng/uL | | | | | |

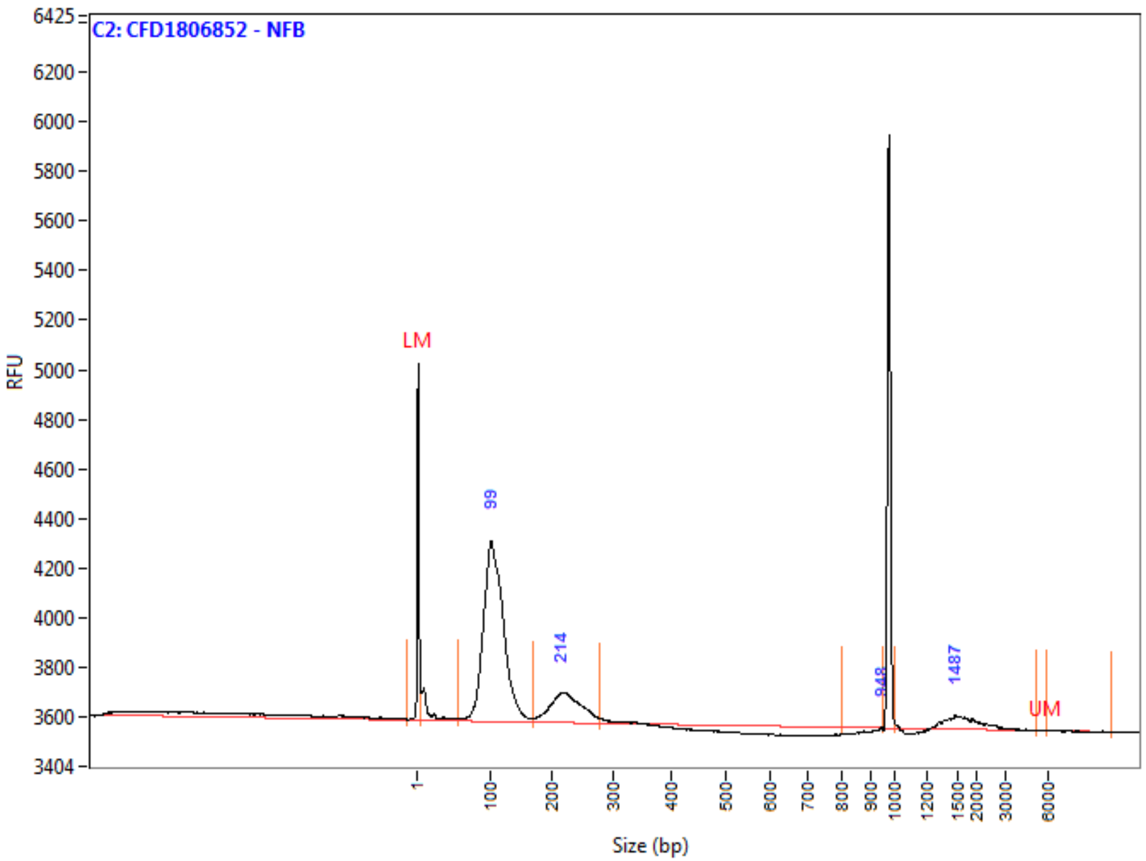

**Peak Table**

| | Size (bp) | ng/uL | % (Conc.) (ng/uL) | nmole/L | From (bp) | To (bp) | Avg. Size | CV% |
|---|---|---|---|---|---|---|---|---|
| 1 | 1 (LM) | 0.0007 | | 0.9487 | 0 | 6 | 1 | 176.80 |
| 2 | 99 | 0.0324 | 78.2 | 0.5068 | 56 | 167 | 105 | 15.19 |
| 3 | 214 | 0.0071 | 17.2 | 0.0527 | 169 | 279 | 222 | 10.84 |
| 4 | 948 | 0.0000 | 0.0 | 0.0000 | 799 | 953 | 947 | 0.09 |
| 5 | 1487 | 0.0019 | 4.6 | 0.0018 | 1007 | 5245 | 1806 | 28.62 |
| 6 | 6000 (UM) | 0.0000 | | 0.0000 | 5954 | 10458 | 7204 | 22.19 |
| | TIC: | 0.0414 | ng/uL | | | | | |
| | TIM: | 0.5612 | nmole/L | | | | | |
| | Total Conc.: | 0.0526 | ng/uL | | | | | |

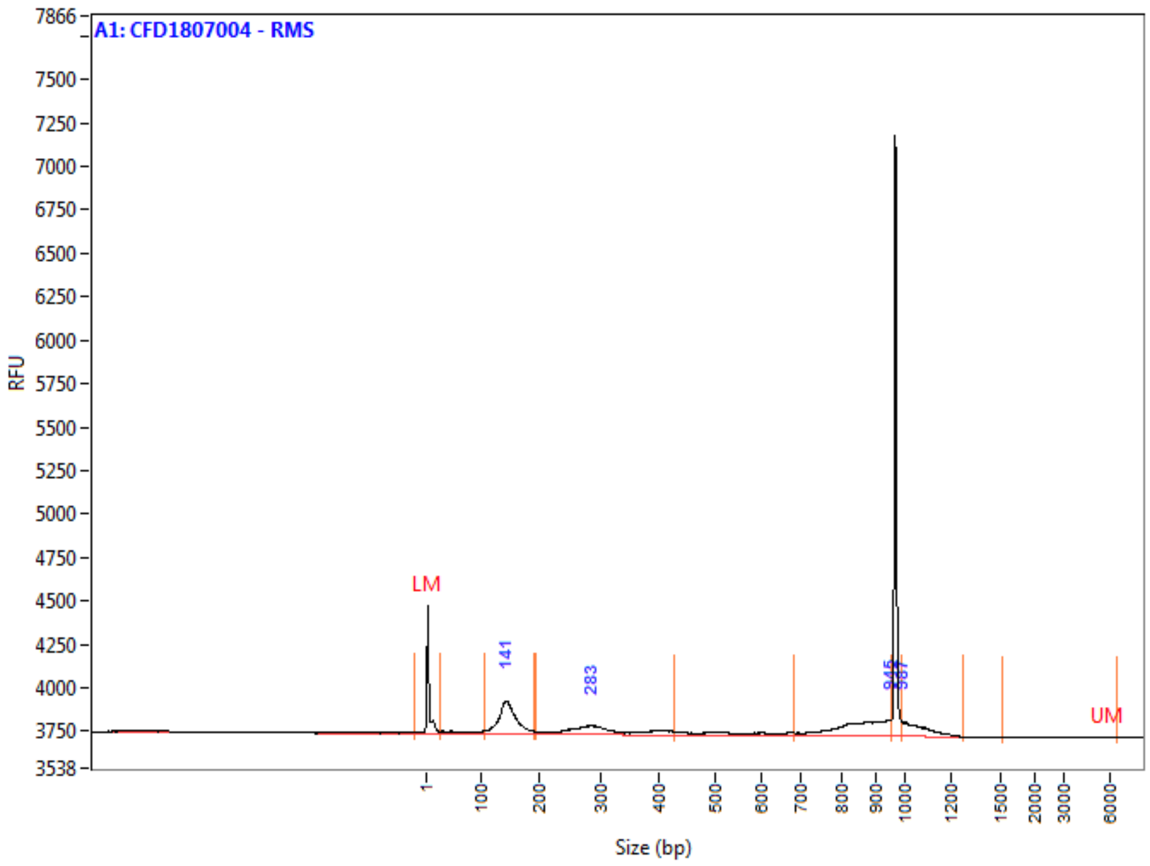

**Peak Table**

| | Size (bp) | ng/uL | % (Conc.) (ng/uL) | nmole/L | From (bp) | To (bp) | Avg. Size | CV% |
|---|---|---|---|---|---|---|---|---|
| 1 | 1 (LM) | 0.0013 | | 0.9492 | 0 | 25 | 2 | 233.80 |
| 2 | 141 | 0.0205 | 37.6 | 0.2339 | 107 | 191 | 144 | 11.17 |
| 3 | 283 | 0.0135 | 24.8 | 0.0732 | 194 | 427 | 304 | 19.54 |
| 4 | 945 | 0.0146 | 26.9 | 0.0285 | 683 | 953 | 846 | 8.01 |
| 5 | 987 | 0.0058 | 10.7 | 0.0090 | 987 | 1272 | 1061 | 5.48 |
| 6 | 6000 (UM) | 0.0000 | | 0.0000 | 1522 | 6542 | 5844 | 6.49 |
| | TIC: | 0.0545 | ng/uL | | | | | |
| | TIM: | 0.3447 | nmole/L | | | | | |
| | Total Conc.: | 0.0883 | ng/uL | | | | | |

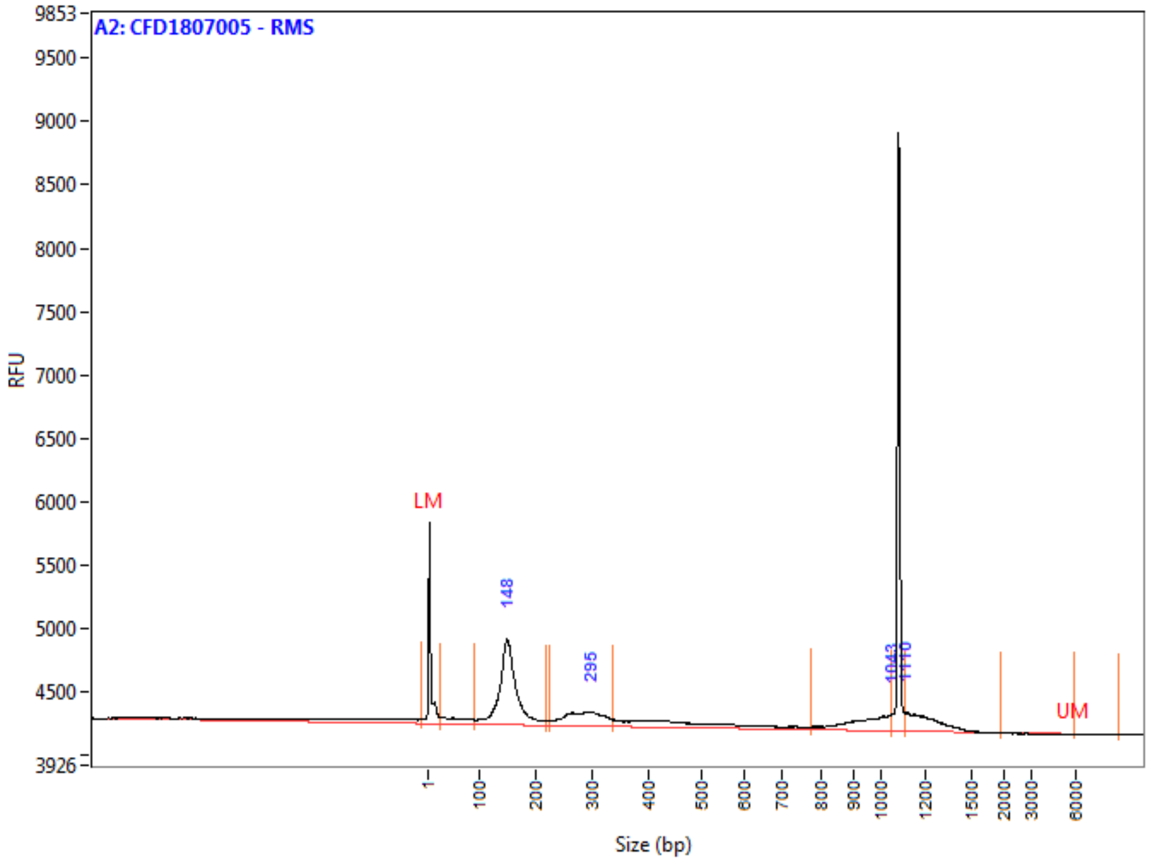

**Peak Table**

| | Size (bp) | ng/uL | % (Conc.) (ng/uL) | nmole/L | From (bp) | To (bp) | Avg. Size | CV% |
|---|---|---|---|---|---|---|---|---|
| 1 | 1 (LM) | 0.0013 | | 0.6580 | 0 | 25 | 3 | 222.60 |
| 2 | 148 | 0.0307 | 57.4 | 0.3385 | 89 | 217 | 149 | 13.78 |
| 3 | 295 | 0.0102 | 19.2 | 0.0598 | 226 | 335 | 282 | 9.80 |
| 4 | 1043 | 0.0065 | 12.2 | 0.0115 | 775 | 1045 | 935 | 7.60 |
| 5 | 1110 | 0.0060 | 11.2 | 0.0081 | 1110 | 1966 | 1223 | 7.45 |
| 6 | 6000 (UM) | | | | 6000 | 9100 | NaN | NaN |
| | TIC: | 0.0535 | ng/uL | | | | | |
| | TIM: | 0.4178 | nmole/L | | | | | |
| | Total Conc.: | 0.0845 | ng/uL | | | | | |

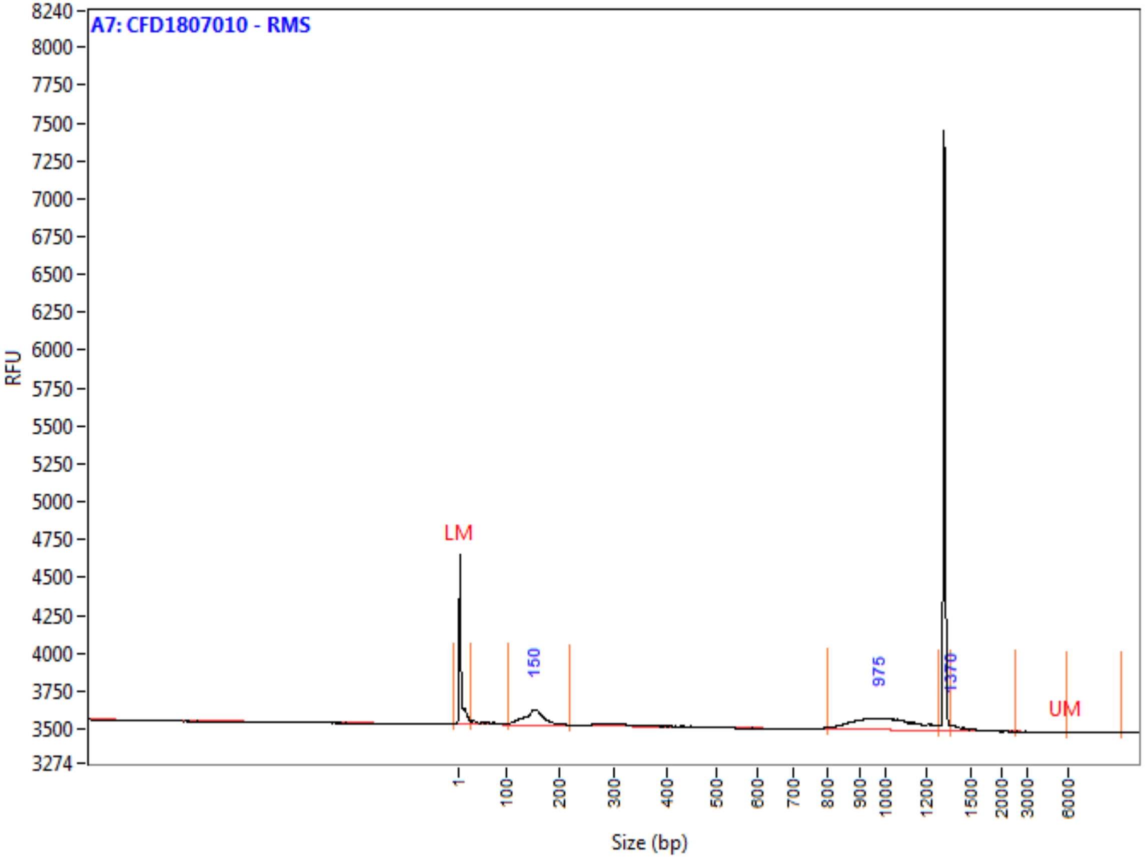

**Peak Table**

| | Size (bp) | ng/uL | % (Conc.) (ng/uL) | nmole/L | From (bp) | To (bp) | Avg. Size | CV% |
|---|---|---|---|---|---|---|---|---|
| 1 | 1 (LM) | 0.0013 | | 0.6580 | 0 | 25 | 3 | 208.31 |
| 2 | 150 | 0.0090 | 40.9 | 0.0997 | 103 | 217 | 149 | 13.93 |
| 3 | 975 | 0.0121 | 54.8 | 0.0198 | 799 | 1286 | 1008 | 11.48 |
| 4 | 1370 | 0.0010 | 4.3 | 0.0011 | 1367 | 2597 | 1462 | 8.65 |
| 5 | 6000 (UM) | | | | 6000 | 10033 | 10033 | 0.00 |
| | TIC: | 0.0221 | ng/uL | | | | | |
| | TIM: | 0.1206 | nmole/L | | | | | |
| | Total Conc.: | 0.0466 | ng/uL | | | | | |

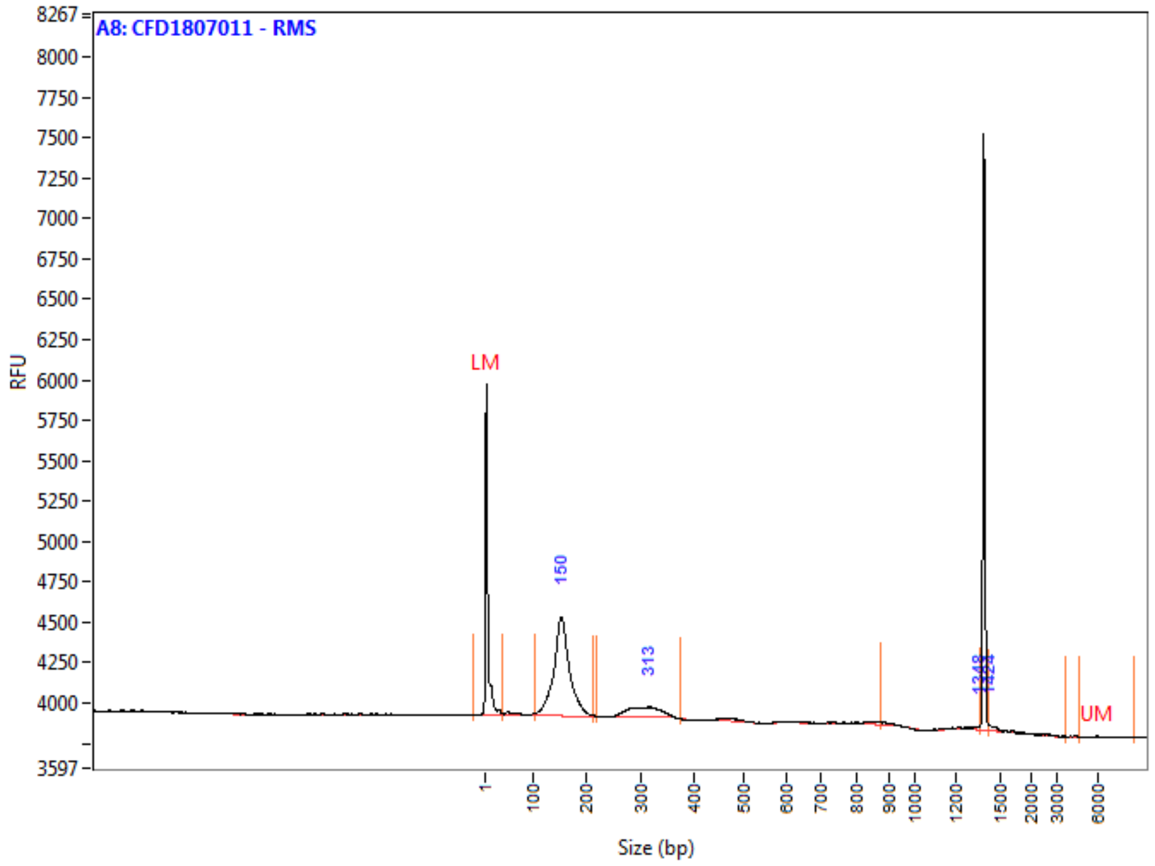

**Peak Table**

| | Size (bp) | ng/uL | % (Conc.) (ng/uL) | nmole/L | From (bp) | To (bp) | Avg. Size | CV% |
|---|---|---|---|---|---|---|---|---|
| 1 | 1 (LM) | 0.0013 | | 0.6580 | 0 | 37 | 3 | 222.05 |
| 2 | 150 | 0.0234 | 82.6 | 0.2550 | 103 | 212 | 151 | 11.16 |
| 3 | 313 | 0.0038 | 13.5 | 0.0210 | 219 | 376 | 301 | 8.14 |
| 4 | 1348 | 0.0006 | 2.1 | 0.0009 | 875 | 1365 | 1122 | 16.22 |
| 5 | 1424 | 0.0005 | 1.8 | 0.0005 | 1424 | 3667 | 1827 | 30.81 |
| 6 | 6000 (UM) | 0.0000 | | 0.0000 | 4697 | 8678 | 6233 | 13.83 |
| | TIC: | 0.0284 | ng/uL | | | | | |
| | TIM: | 0.2774 | nmole/L | | | | | |
| | Total Conc.: | 0.0404 | ng/uL | | | | | |

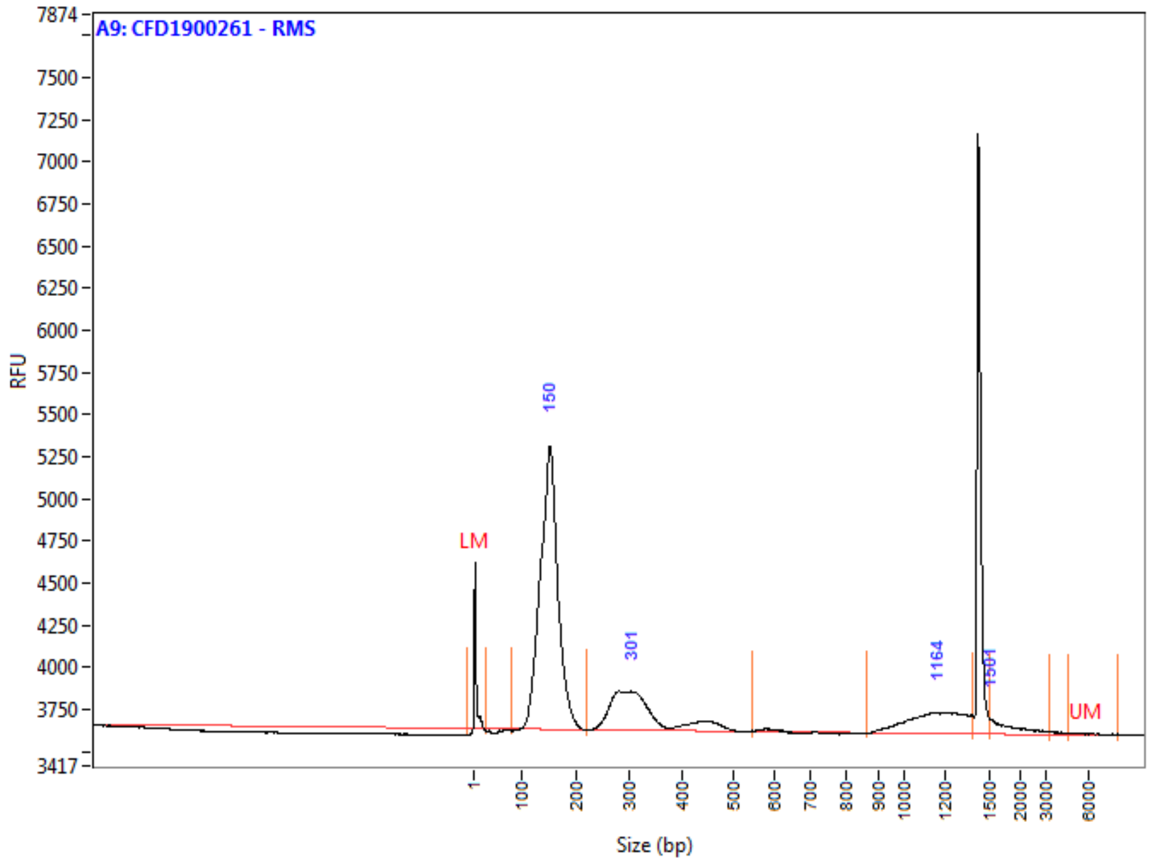

**Peak Table**

| | Size (bp) | ng/uL | % (Conc.) (ng/uL) | nmole/L | From (bp) | To (bp) | Avg. Size | CV% |
|---|---|---|---|---|---|---|---|---|
| 1 | 1 (LM) | 0.0013 | | 1.7025 | 0 | 26 | 1 | 273.60 |
| 2 | 150 | 0.1686 | 68.5 | 1.8853 | 77 | 219 | 147 | 11.52 |
| 3 | 301 | 0.0496 | 20.1 | 0.2547 | 219 | 545 | 320 | 18.58 |
| 4 | 1164 | 0.0225 | 9.1 | 0.0320 | 866 | 1387 | 1156 | 10.83 |
| 5 | 1501 | 0.0056 | 2.3 | 0.0048 | 1501 | 3274 | 1938 | 21.69 |
| 6 | 6000 (UM) | 0.0001 | | 0.0000 | 4637 | 8287 | 5986 | 16.91 |
| | TIC: | 0.2463 | ng/uL | | | | | |
| | TIM: | 2.1768 | nmole/L | | | | | |
| | Total Conc.: | 0.2887 | ng/uL | | | | | |

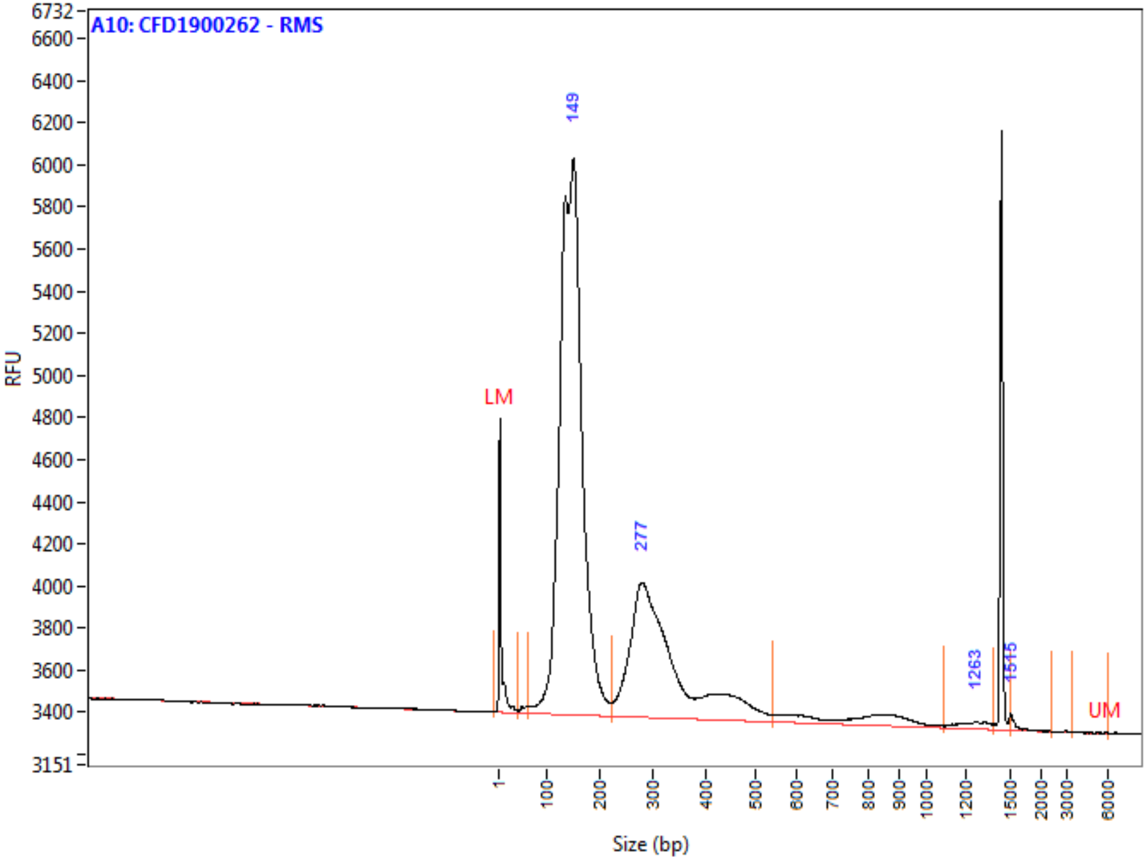

**Peak Table**

| | Size (bp) | ng/uL | % (Conc.) (ng/uL) | nmole/L | From (bp) | To (bp) | Avg. Size | CV% |
|---|---|---|---|---|---|---|---|---|
| 1 | 1 (LM) | 0.0013 | | 0.5036 | 0 | 39 | 4 | 199.31 |
| 2 | 149 | 0.2076 | 70.3 | 2.3691 | 62 | 221 | 144 | 14.80 |
| 3 | 277 | 0.0853 | 28.9 | 0.4370 | 221 | 541 | 321 | 21.22 |
| 4 | 1263 | 0.0019 | 0.6 | 0.0025 | 1089 | 1387 | 1245 | 6.50 |
| 5 | 1515 | 0.0007 | 0.2 | 0.0007 | 1508 | 2436 | 1613 | 8.71 |
| 6 | 6000 (UM) | 0.0000 | | 0.0000 | 3455 | 6120 | 5365 | 12.09 |
| | TIC: | 0.2955 | ng/uL | | | | | |
| | TIM: | 2.8093 | nmole/L | | | | | |
| | Total Conc.: | 0.3196 | ng/uL | | | | | |

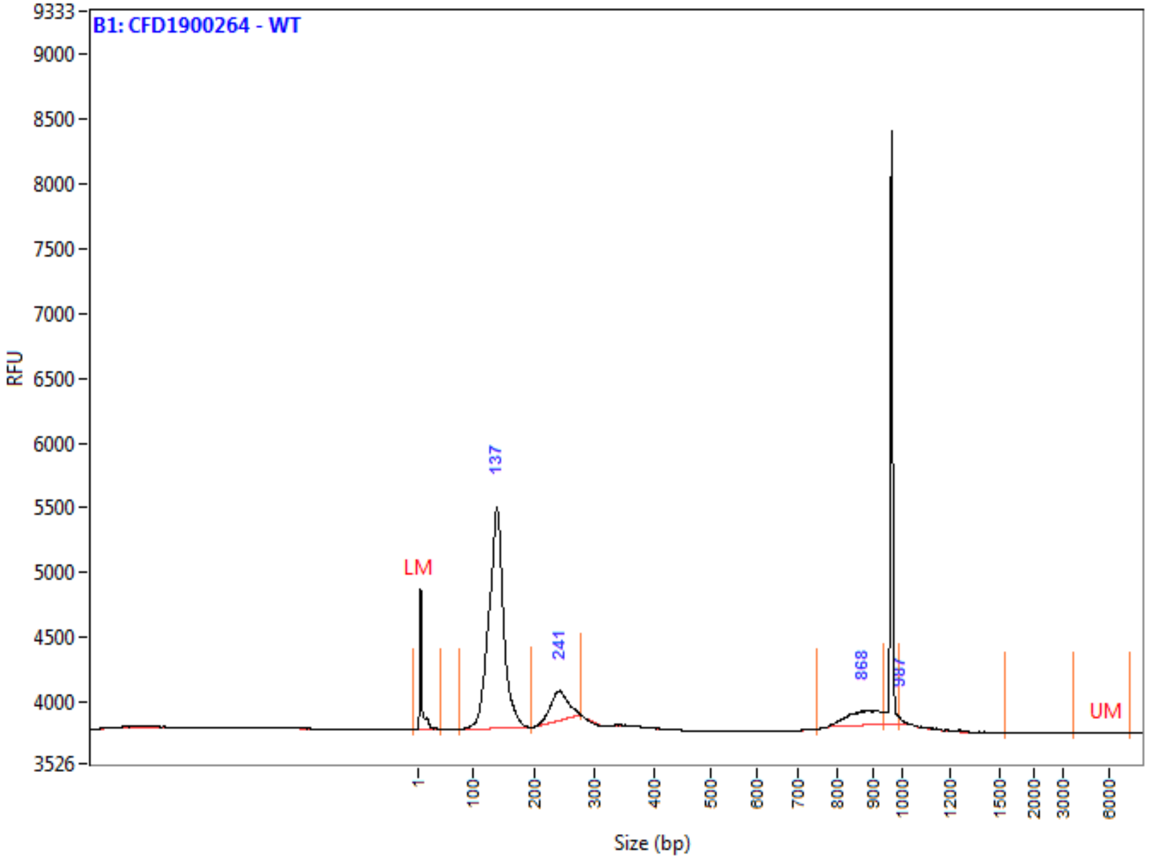

**Peak Table**

| | Size (bp) | ng/uL | % (Conc.) (ng/uL) | nmole/L | From (bp) | To (bp) | Avg. Size | CV% |
|---|---|---|---|---|---|---|---|---|
| 1 | 1 (LM) | 0.0013 | | 0.6580 | 0 | 41 | 3 | 202.51 |
| 2 | 137 | 0.1090 | 81.6 | 1.3372 | 72 | 195 | 134 | 10.22 |
| 3 | 241 | 0.0151 | 11.3 | 0.1039 | 195 | 276 | 239 | 5.66 |
| 4 | 868 | 0.0085 | 6.4 | 0.0162 | 748 | 934 | 867 | 4.52 |
| 5 | 987 | 0.0009 | 0.7 | 0.0014 | 987 | 1597 | 1122 | 13.58 |
| 6 | 6000 (UM) | 0.0000 | | 0.0000 | 3698 | 7414 | 5590 | 14.89 |
| | TIC: | 0.1336 | ng/uL | | | | | |
| | TIM: | 1.4586 | nmole/L | | | | | |
| | Total Conc.: | 0.1590 | ng/uL | | | | | |

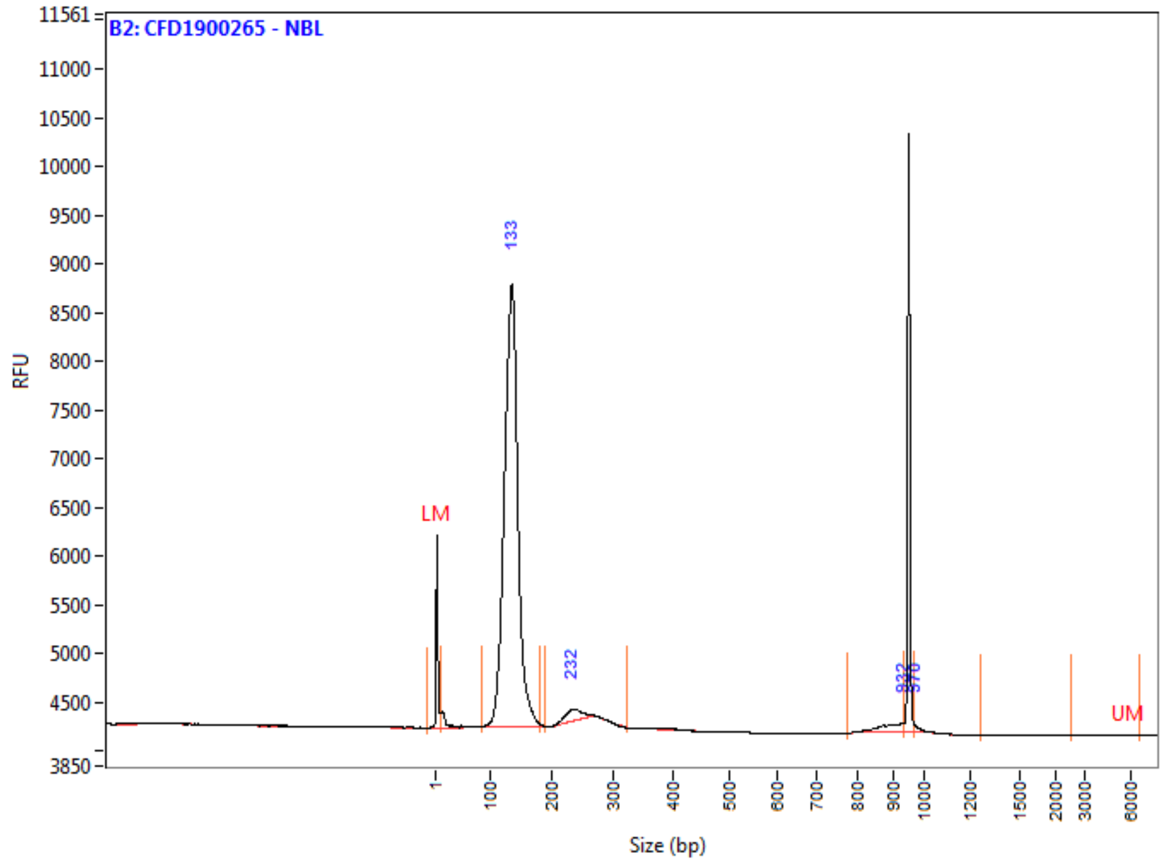

**Peak Table**

| | Size (bp) | ng/uL | % (Conc.) (ng/uL) | nmole/L | From (bp) | To (bp) | Avg. Size | CV% |
|---|---|---|---|---|---|---|---|---|
| 1 | 1 (LM) | 0.0013 | | 1.7025 | 0 | 9 | 1 | 295.65 |
| 2 | 133 | 0.1790 | 95.7 | 2.2450 | 85 | 182 | 131 | 8.97 |
| 3 | 232 | 0.0051 | 2.7 | 0.0356 | 190 | 323 | 235 | 5.91 |
| 4 | 932 | 0.0025 | 1.3 | 0.0047 | 775 | 936 | 888 | 3.56 |
| 5 | 970 | 0.0005 | 0.3 | 0.0008 | 970 | 1266 | 1009 | 6.93 |
| 6 | 6000 (UM) | 0.0000 | | 0.0000 | 2581 | 6632 | 4674 | 31.86 |
| | TIC: | 0.1871 | ng/uL | | | | | |
| | TIM: | 2.2861 | nmole/L | | | | | |
| | Total Conc.: | 0.2115 | ng/uL | | | | | |

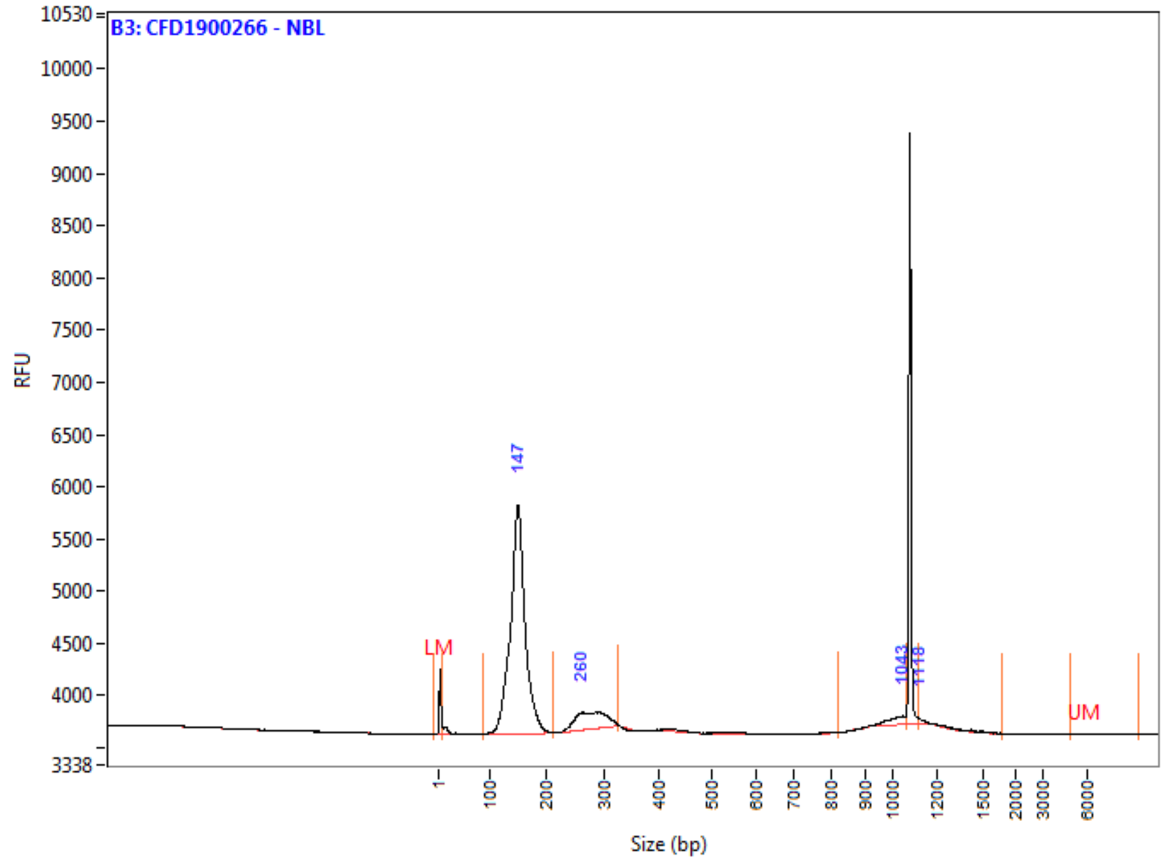

**Peak Table**

| | Size (bp) | ng/uL | % (Conc.) (ng/uL) | nmole/L | From (bp) | To (bp) | Avg. Size | CV% |
|---|---|---|---|---|---|---|---|---|
| 1 | 1 (LM) | 0.0013 | | 8.2518 | 0 | 6 | 0 | 1448.4[illegible] |
| 2 | 147 | 0.2950 | 85.4 | 3.3209 | 86 | 212 | 146 | 10.21 |
| 3 | 260 | 0.0405 | 11.7 | 0.2433 | 212 | 327 | 274 | 7.94 |
| 4 | 1043 | 0.0067 | 2.0 | 0.0110 | 819 | 1063 | 1007 | 4.07 |
| 5 | 1118 | 0.0030 | 0.9 | 0.0039 | 1118 | 1802 | 1285 | 13.09 |
| 6 | 6000 (UM) | 0.0000 | | 0.0000 | 4940 | 9702 | 6610 | 20.74 |
| | TIC: | 0.3453 | ng/uL | | | | | |
| | TIM: | 3.5791 | nmole/L | | | | | |
| | Total Conc.: | 0.4173 | ng/uL | | | | | |

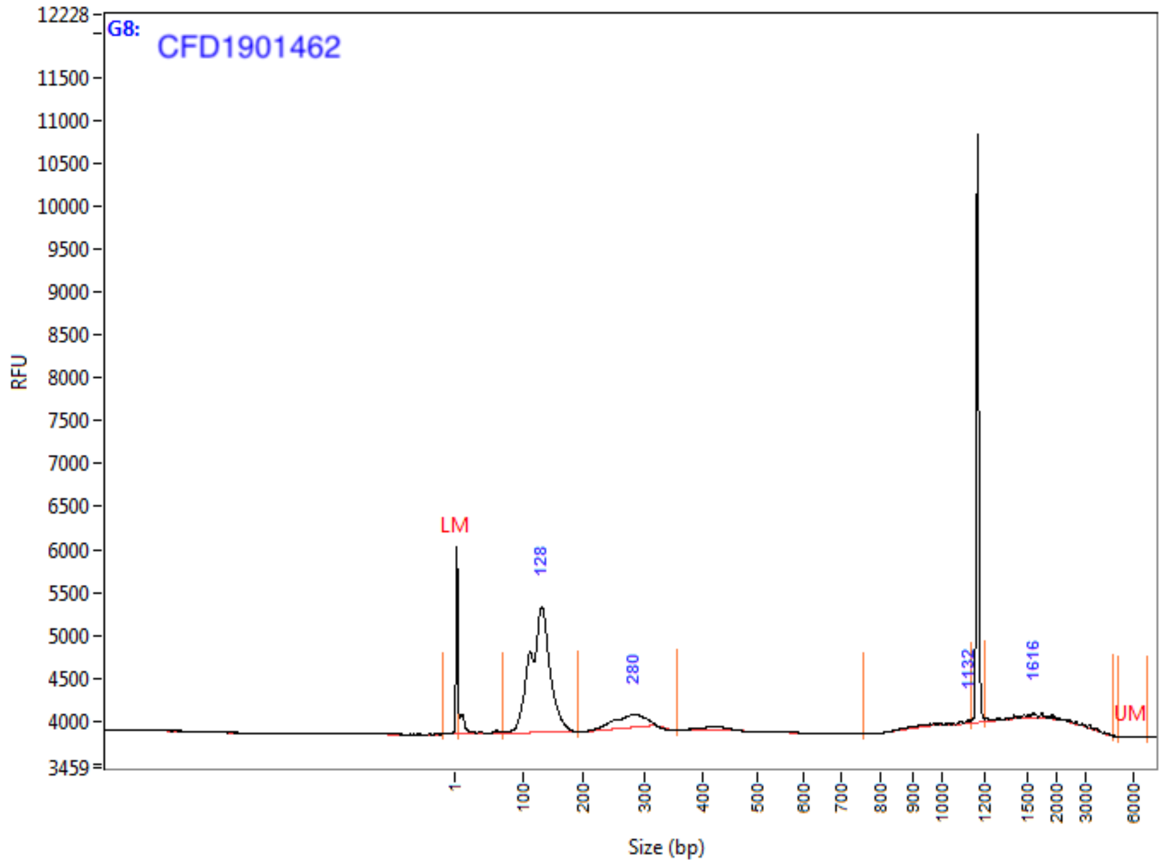

**Peak Table**

| | Size (bp) | ng/uL | % (Conc.) (ng/uL) | nmole/L | From (bp) | To (bp) | Avg. Size | CV% |
|---|---|---|---|---|---|---|---|---|
| 1 | 1 (LM) | 0.0003 | | 0.4320 | 0 | 5 | 1 | 226.00 |
| 2 | 128 | 0.0185 | 84.4 | 0.2447 | 68 | 191 | 124 | 13.78 |
| 3 | 280 | 0.0025 | 11.5 | 0.0153 | 191 | 357 | 269 | 8.80 |
| 4 | 1132 | 0.0003 | 1.6 | 0.0006 | 756 | 1141 | 980 | 9.95 |
| 5 | 1616 | 0.0006 | 2.5 | 0.0005 | 1211 | 4849 | 2009 | 42.40 |
| 6 | 6000 (UM) | 0.0000 | | 0.0000 | 5182 | 7022 | 5977 | 6.40 |
| | TIC: | 0.0219 | ng/uL | | | | | |
| | TIM: | 0.2610 | nmole/L | | | | | |
| | Total Conc.: | 0.0289 | ng/uL | | | | | |

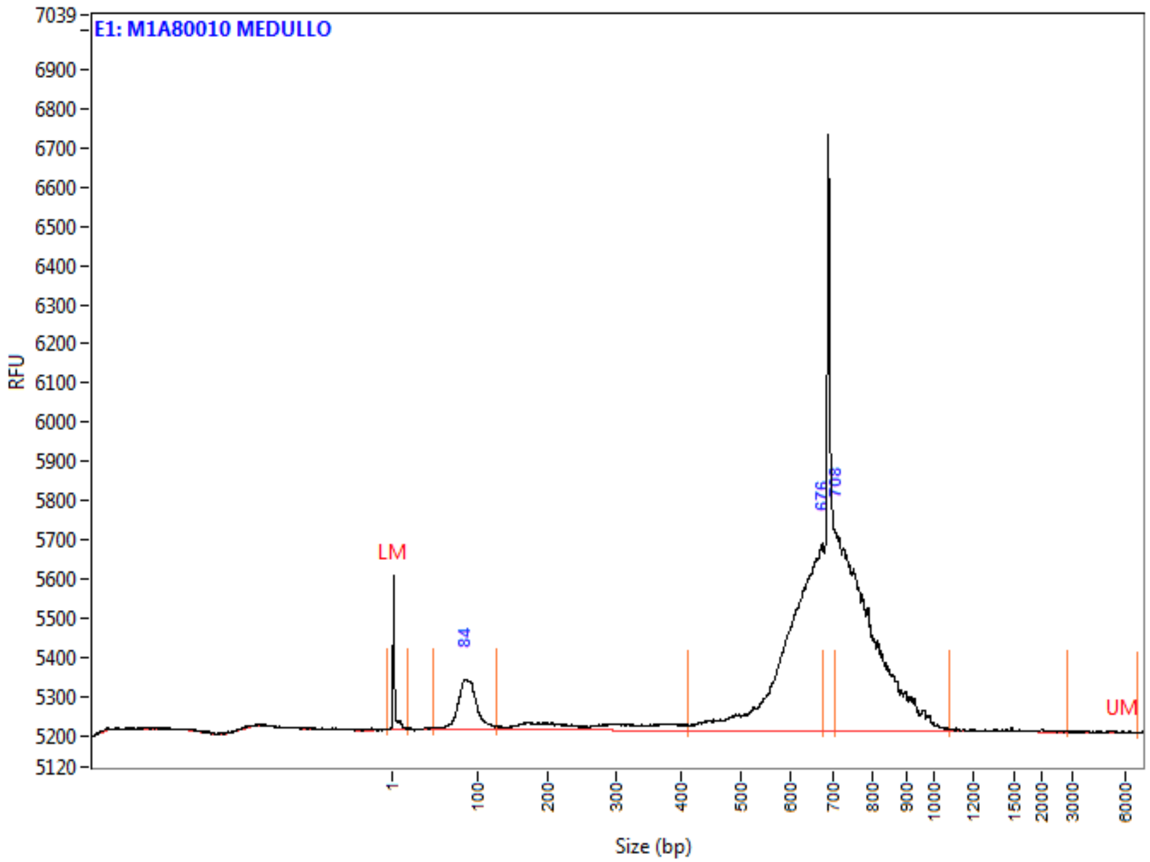

**Peak Table**

| | Size (bp) | ng/uL | % (Conc.) (ng/uL) | nmole/L | From (bp) | To (bp) | Avg. Size | CV% |
|---|---|---|---|---|---|---|---|---|
| 1 | 1 (LM) | 0.0005 | | 0.3322 | 0 | 20 | 2 | 161.24 |
| 2 | 84 | 0.0137 | 11.7 | 0.2576 | 49 | 128 | 87 | 13.32 |
| 3 | 676 | 0.0506 | 43.2 | 0.1374 | 411 | 677 | 606 | 9.57 |
| 4 | 708 | 0.0529 | 45.1 | 0.1108 | 707 | 1080 | 785 | 8.47 |
| 5 | 6000 (UM) | 0.0000 | | 0.0000 | 2898 | 6807 | 5092 | 17.95 |
| | TIC: | 0.1171 | ng/uL | | | | | |
| | TIM: | 0.5058 | nmole/L | | | | | |
| | Total Conc.: | 0.1477 | ng/uL | | | | | |
